# Validation of a stereological method for estimating particle size and density from 2D projections with high accuracy

**DOI:** 10.1101/2022.10.21.513285

**Authors:** Jason Seth Rothman, Carolina Borges-Merjane, Noemi Holderith, Peter Jonas, R. Angus Silver

## Abstract

Stereological methods for estimating the 3D particle size and density from 2D projections are essential to many research fields. These methods are, however, prone to errors arising from undetected particle profiles due to sectioning and limited resolution, known as ‘lost caps’. A potential solution by Keiding et al. (1972) accounts for lost caps by quantifying the smallest detectable profiles in terms of their limiting section angle (ϕ). However, this simple solution has not been widely adopted nor validated. Rather, model-independent design-based stereological methods, which do not explicitly account for lost caps, have come to the fore. Here, we provide the first experimental validation of the Keiding model by quantifying ϕ of synaptic vesicles using electron-tomography 3D reconstructions. This analysis reveals a Gaussian distribution for ϕ rather than a single value. Nevertheless, curve fits of the Keiding model to the 2D diameter distribution accurately estimate the mean ϕ and 3D diameter distribution. While systematic testing using Monte Carlo simulations revealed an upper limit to determining ϕ, our analysis shows that mean ϕ can be used to estimate the 3D particle density from the 2D density under a wide range of conditions, and this method is potentially more accurate than minimum-size-based lost-cap corrections and disector methods. We applied the Keiding model to estimate the size and density of somata, nuclei and vesicles in rodent cerebella, where high packing density can be problematic for design-based methods.

## 1 Introduction

Estimating the size and density of particles from their orthogonal projection, such as a 2D image, is a common stereological endeavour in the fields of biosciences, petrography, materials science and astronomy^1^. This approach is particularly valuable in the field of biosciences where the size and density of biological structures, such as cells and organelles, are often compared before and after drug perturbations, between normal and disease conditions, species, ages or critical periods of development^2^. Moreover, measures of particle size and density, or the equivalent measure of volume fraction (VF; see Definition of Key Terms), form the basis of our understanding of a wide range of biological phenomena. For example, the density of synaptic vesicles near the active zone has been related to measures of synaptic plasticity^3–5^, the density of cerebellar granule cells (GCs) and mossy fiber terminals (MFTs) has been used to estimate the amount of information transferred across the input layer of the cerebellum^6^ and the amount of energy expenditure at the cellular and subcellular level^7^. Stereological measures are also commonly used to assay disease states, such as that of the lung, liver, kidney and brain^8–11^. Hence, stereological methods for estimating particle size and density have wide application and are of great practical utility.

Recent advances in high-resolution volumetric imaging have significantly improved the morphological information available about cells and tissue structure^12–15^ making them ideal for 3D analysis. However, these technologies are expensive and full reconstructions are both labour and computationally intensive. The use of stereological methods for analysing a relatively small sample of 2D projections is therefore still the most time efficient and practical solution for most laboratories.

Stereological methods for estimating size and density have developed along two distinct approaches: a model-based approach that makes basic assumptions about the geometry of the particle of interest, e.g. Wicksell’s transformation^16^ and the Abercrombie correction^17^ that assume a spherical geometry, versus a design-based approach that makes no assumption about particle geometry, e.g. the nucleator, rotator, physical and optical disector^10,18,19^. The ability to function for particles with an arbitrary shape is one of the reasons design-based methods are often referred to as ‘assumption-free’ or ‘unbiased’ and considered the superior approach^2,11,19–23^. However, design-based methods are not free of assumptions and may contain biases^24–28^. Moreover, design-based methods can be labour intensive and costly^26,28–30^ and are not appropriate for particles with a high density^18^. A high particle density occurs in many types of preparations, including vesicles in synapses^31,32^, granules in chromaffin and mast cells^33,34^, granule cells in the cerebellum and hippocampus^6,35^ and corneal epithelial basal cells^36^. Model-based methods, on the other hand, not only have the potential to be more efficient, but can offer more information about particle size^37^, higher levels of accuracy^30^ and can be applied to particles with either a low or high density.

A classic problem addressed by model-based stereological approaches is the estimation of particle size from a 2D projection, such as an image. This was first studied by Wicksell^16^ and is known as Wicksell’s corpuscle problem (Figure 1A). Wicksell’s problem was to infer the true 3D diameter distribution (F(d)) of secondary follicles from their 2D diameter distribution (G(d)) measured from images of planar sections of the human spleen, where the thickness of the sections (T) was much thinner than the mean particle diameter (μ_D_). Wicksell’s solution to this inverse problem was to use a model-based approach to derive an analytical solution for G(d) with respect to F(d) and then use a finite-difference unfolding algorithm to estimate F(d) from G(d).

**Figure 1.**
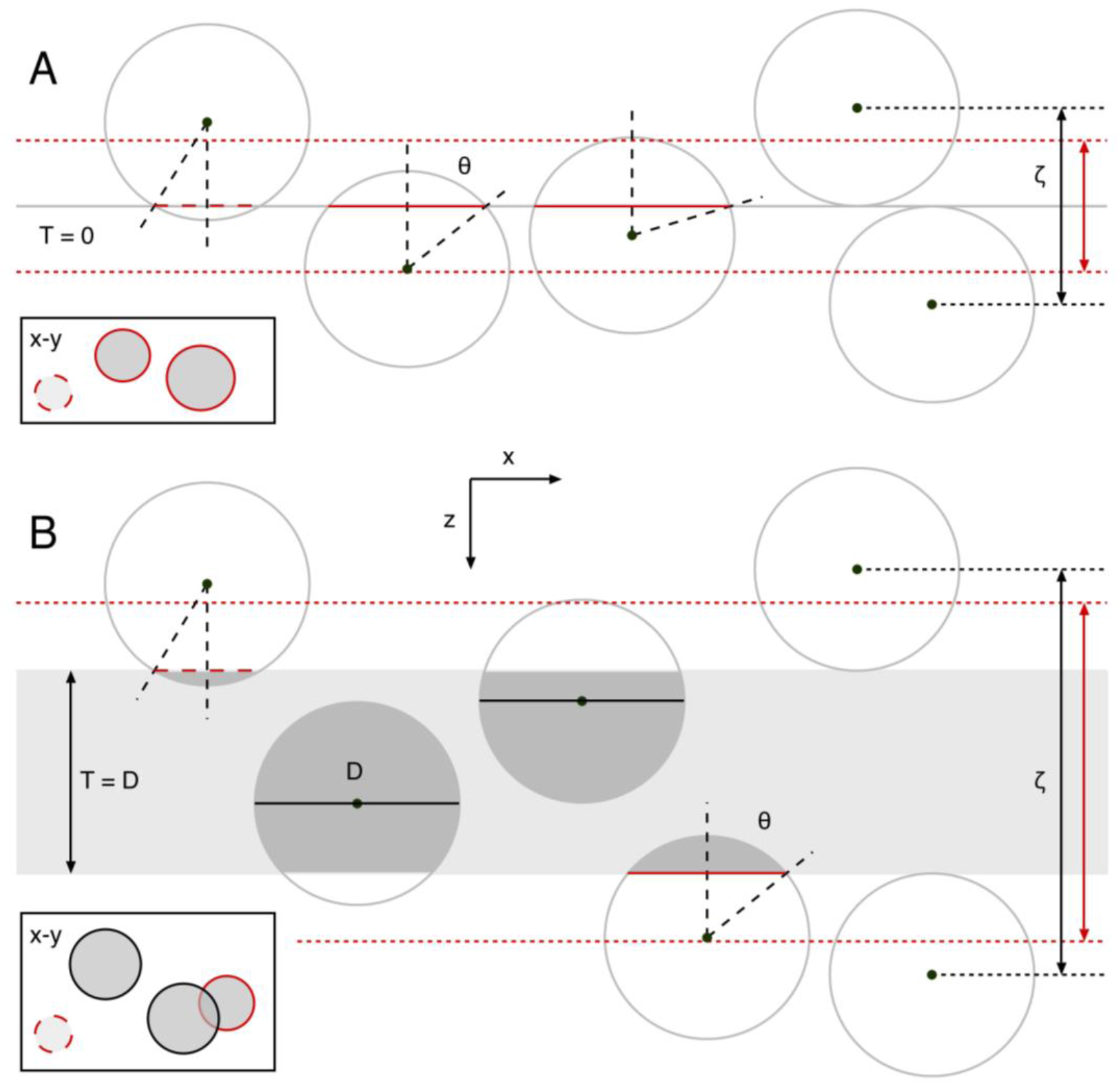
Illustration of observed circular profiles when spherical particles are orthogonally viewed from above a planar or thick section. (A) Side view of a planar section (T = 0; gray line) transecting spherical particles (red solid lines). For simplicity, all particles have the same 3D diameter (D). Particles with their center above/below the section within a distance D/2 are observed as circular ‘caps’ in a horizontal projection (bottom rectangle, red circles) with apparent diameter d < D, where d = D·sin and θ is the cap angle (black dashed lines; 0 < θ < 90°). Hence, particles appearing within the projection have their center confined within a depth ζ = D (black double-headed arrow). However, due to experimental limitations, the smallest caps are not apparent, i.e. lost (red dashed lines). To account for lost caps, the Keiding model sets a minimum limit on θ (ϕ) such that caps are observed in the projection only if ϕ < θ < 90°. In this case, ζ = D·cosϕ (red double-headed arrow; Equation 2). For planar sections, there are no projection overlaps and the total area fraction (AF) of the projections approximately equals the 3D volume fraction (VF) of the particles so long as there are relatively few lost caps (Appendix B). Because all particle centers fall above/below the planar section, all particles are considered caps. (B) Same as (A) for a thick section (T = D). In this case, particles with their center within the section have a circular projection with d = D (black circles) and ζ = T + D·cosϕ. For thick sections and a high particle density, there are usually projection overlaps that make counting/outlining the projections more difficult; moreover, AF > VF, a condition known as overprojection.

A perhaps more common scenario than measuring particle profiles from a planar section is that of measuring particle profiles observed through a transparent section of thickness T (Figure 1B). The analytical solution for G(d) with respect to F(d) for T ≥ 0 was derived by Bach^38^ and can be described as the weighted sum of two components^39^: the diameter distribution of those particles with their center points contained within the section, in which case their G(d) = F(d), and the diameter distribution of those particles with their center points just above and below the section, by less than one radius, in which case their G(d) is a distorted version of F(d) as defined by Wicksell’s analytical solution^16^. These latter particles whose north and south poles appear on the bottom and top of the section are known as ‘caps’. Besides distorting the diameter distribution, caps also introduce a distortion of the apparent density, an effect known as overprojection, the split-cell error or the Holmes effect^17,40–43^.

A fundamental limitation of the Wicksell^16^ and Bach^38^ models, however, is that they assume all caps are resolvable. While this might be true for the largest caps with diameters on the order of F(d), the smallest caps are usually unresolvable, falling below the limits of resolution and contrast or blending in with their surrounding environment^44,45^. Other caps might simply not exist if they fall off the surfaces of the sections or if the microtome fails to transect the particles during sectioning^27,46–48^. Wicksell noted that lost caps could be accounted for by a post-hoc correction of his unfolding algorithm, whereby the missing probabilities of the smallest bins of F(d) are estimated via extrapolation from the smallest non-zero probability down to the origin. However, this approach is problematic since it relies on a small number of outlier observations. Indeed, when the number of observations within the smallest bins are insufficient, the unfolding algorithm can generate erroneous negative probabilities.

It was not until the 1970s that a key innovation for accounting for lost caps was developed by Keiding et al.^49^ whereby lost caps are defined with respect to a lower limit of the angle subtended by the observable caps (cap-angle limit, ϕ; Figure 1). In this conceptual model, ϕ is independent of particle diameter, which is important since a distribution of limiting cap sizes can arise when particle size varies while specimen contrast, rather than microscope resolution, limits cap detection. Incorporating ϕ into the Wicksell-Bach model, Keiding et al. derived the following relationship between F(d) and G(d) for spherical particles:

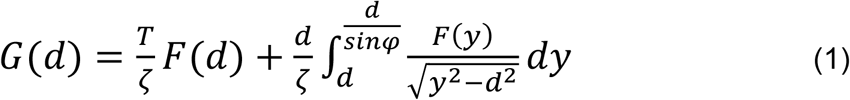

where d is the 2D particle diameter, *y* is the variable of integration and ϕ can vary from 0°, where no caps are lost, to 90°, where all caps are lost. Here, ζ is the mean axial length spanning from below to above the section that contains the center points of those particles observed within the projection (Figure 1), defined as follows:

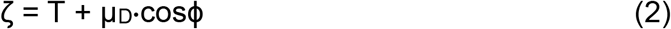

As expected, Equation 1 reduces to Bach’s analytical solution when ϕ = 0° and to Wicksell’s analytical solution when ϕ = 0° and T = 0.

Another innovation of Keiding et al.^49^ was to estimate both F(d) and ϕ from G(d) using a maximum likelihood estimation (MLE) algorithm, rather than using an unfolding algorithm, thereby providing a better quantification of ϕ. Knowing ϕ is particularly useful since it can be used via Equation 2 to estimate the 3D particle density (λ_3D_) from the measured 2D particle density (λ_2D_) as follows^50^:

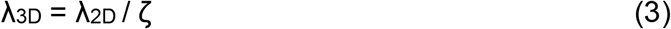

(Appendix A). This ‘correction’ method for estimating λ_3D_ is potentially more accurate than using the classic Abercrombie correction^17^, which assumes no caps are lost (i.e. ϕ = 0°), or the Floderus^42^ and Konigsmark^47^ corrections that use the minimum cap penetration depth (h_min_) or equivalent minimum cap diameter (d_min_), both of which are likely to be outlier measures. Unfortunately, despite potentially providing the most accurate description of the lost-cap distribution via ϕ, the Keiding model has not been widely adopted nor validated. Validation of model-based approaches is important since they are based on simplifying assumptions of particle geometry^22,24,28,51^.

To investigate whether the Keiding model can provide a simple and accurate method for estimating size and density of spherical particles such as synaptic vesicles and nuclei, we used a distribution-based least-squares estimation (LSE) algorithm to systematically test the model’s performance in estimating F(d) and ϕ from G(d) computed from 3D Monte Carlo simulations and electron-tomography (ET) reconstructions. This analysis confirmed the accuracy of the Keiding model in estimating the ‘true’ F(d) and ϕ over a range of conditions. However, the accuracy of estimated ϕ was limited by the sample size, spread of F(d) and the number (i.e. distribution) of lost caps. Finally, we tested the accuracy of Equation 3 for estimating λ_3D_ from the measured λ_2D_ using the same 3D simulations and reconstructions and found this method to be more accurate than the Abercrombie^17^ and d_min_ corrections^47^ and the widely used disector method^18,52^. To facilitate the adoption of the Keiding model in stereological applications, we provide an analysis workflow for estimating F(d), ϕ and λ_3D_ from 2D projections and provide guidelines for optimising particle cap detection. Moreover, we incorporated our numerical solution of Equation 1, including LSE curve-fit functions, into the open-source software toolkit package NeuroMatic^53^ that works within Igor Pro (Key Resources).

## 2 Results

### 2.1 Properties of 2D diameter distributions from images of somata, nuclei and vesicles in the cerebellar cortex

To investigate the properties of G(d) over a range of experimental conditions, including particle size, section thickness, imaging technique and spatial resolution, we quantified the 2D diameters of GC somata and nuclei and MFT vesicles using confocal and transmission electron microscopy (TEM) images of cerebellar sections. These preparations encompassed both planar sections (where T << μ_D_; Figure 1A) and thick sections (where T ≈ μ_D_; Figure 1B).

First, we computed G(d) for cerebellar GC somata from confocal images of rat brain sections from a previous study^6^ (Figure 2A1). In these images, GC somata were visible due to Kv4.2 immunolabeling. However, because the GC somata had an opaque staining and were tightly packed together, there was a high probability of partial and complete overlaps of the 2D profiles, especially since the sections were not planar (T ≈ 1.8 μm; Table 1). To compute G(d), we drew outlines around the GC somata and computed a normalised histogram from the equivalent diameters of the areas of the outlines (d_area_). As predicted by Wicksell^16^, G(d) of the GC somata had an asymmetrical shape with a negative skew (Figure 2A2). However, the negative ‘tail’ of the distribution only extended to ∼2 μm rather than 0 μm, since we were unable to detect GC somata profiles with d < 2 μm (i.e. lost caps). In total, we computed 9 G(d) of GC somata that spanned 2–8 μm with a mean 2D diameter μ_d_ = 4.96–5.83 μm and standard deviation σ_d_ = 0.61–0.90 μm (n = 3 rats, 2–3 cerebellar sections per rat, 494–638 diameters per G(d)).

**Table 1.**
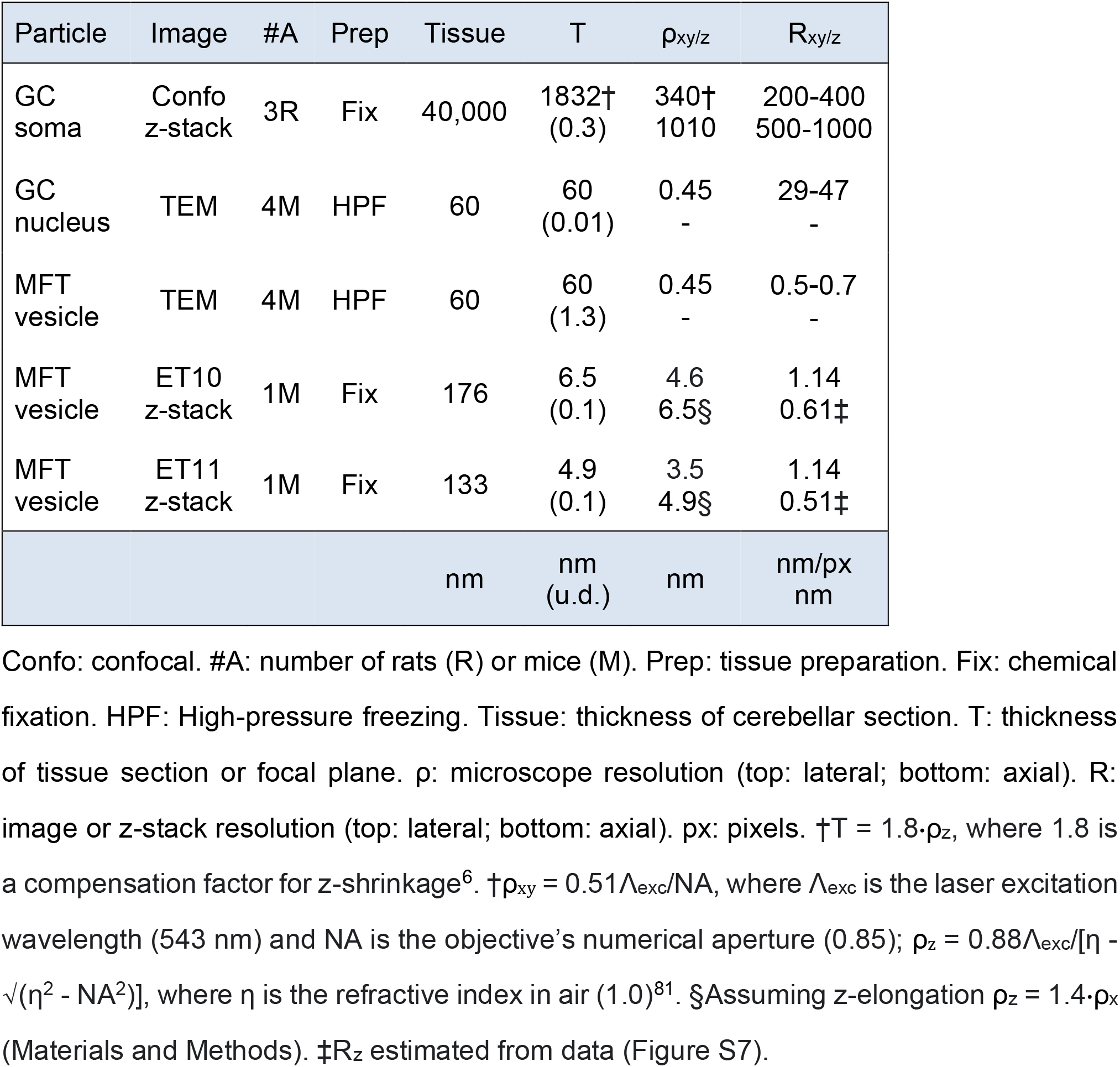
Experimental conditions for confocal, TEM and ET imaging.

**Figure 2.**
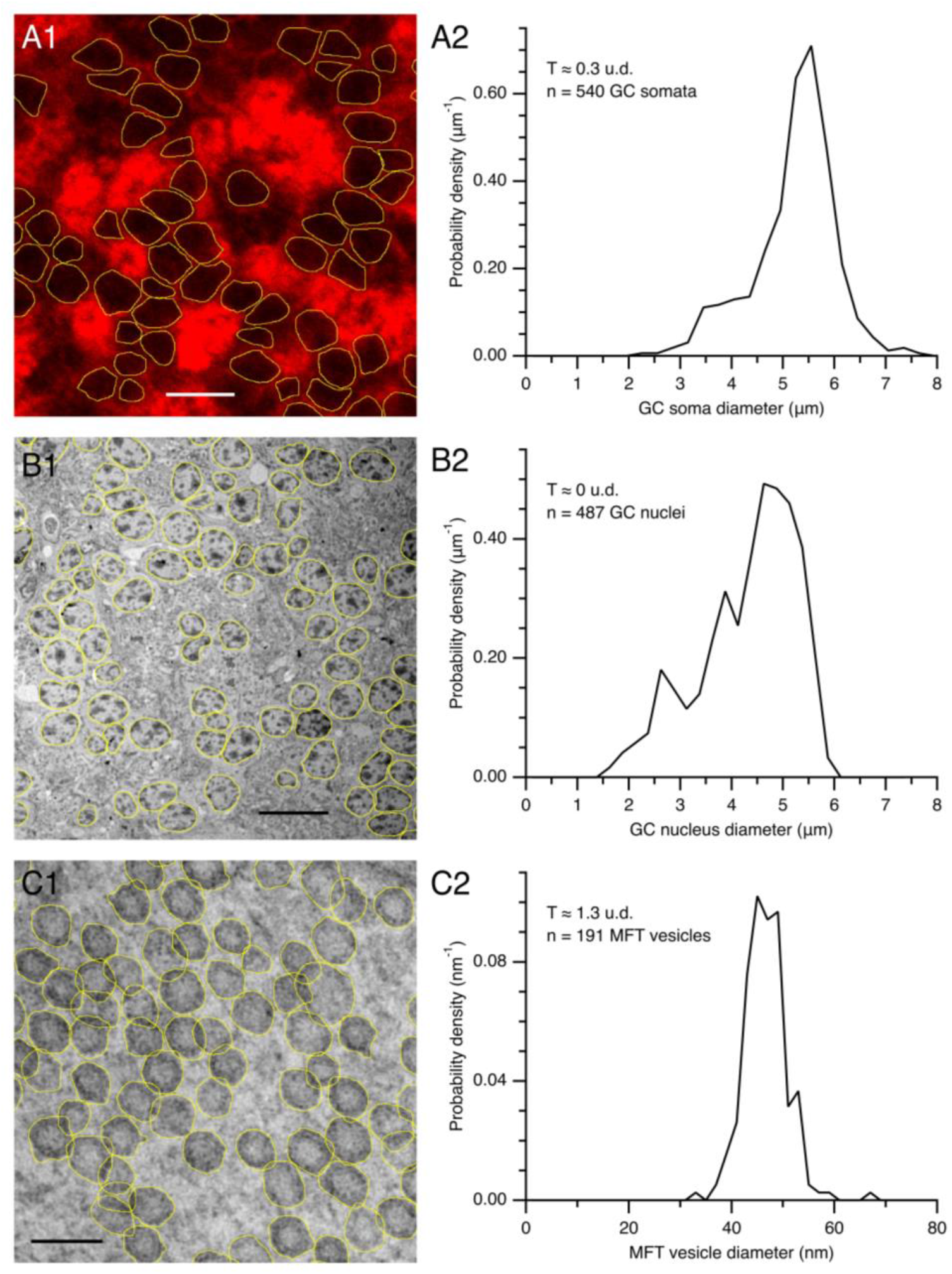
Computing G(d) of GC somata and nuclei and MFT vesicles. (A1) Confocal image of a cerebellar section of a WT rat (P30). GC somatic plasma membranes were delineated via immunolabeling for Kv4.2. Outlines were drawn around those GC somata that were well delineated (yellow) and an equivalent diameter was computed from the area of each outline (d_area_). T ≈ 1.8 μm. Scale bar 10 μm. Image ID R5.SL2.1. (A2) Probability density of 2D diameters (G(d)) computed from the GC somata diameters (0.30 μm bins) measured from the image in (A1) plus 1 other image from the same z-stack. (B1) Low-magnification TEM image of a cerebellar section of a WT mouse (P31). Outlines were drawn around the outer contour of visually identified GC nuclei (yellow). T ≈ 60 nm. Scale bar 10 μm. Image ID M18.N2.51. (B2) G(d) computed from nucleus diameters (0.25 μm bins) measured from the image in (B1) plus 6 other images from the same mouse. (C1) High-magnification TEM image of a MFT in the GC layer of the same mouse in (B1). Outlines were drawn around the outer contour of the synaptic vesicles (yellow). T ≈ 60 nm. Because vesicles are semi-transparent, 2D overlaps do not necessarily preclude drawing their outline or counting. Scale bar 80 nm. Image ID M18.N2.03. (C2) G(d) computed from the vesicle diameters (2 nm bins) measured from the TEM image in (C1). For (A1), (B1) and (C1) only a subregion of the outline analysis is shown.

To examine whether a more complete G(d) could be obtained from images where the resolution is higher, we computed G(d) for GCs in TEM images of mouse cerebellar sections (Figure 2B1; Table 1). Because the sections for this preparation were ∼60 nm thick, the sections were essentially planar (T << μ_D_) with no overlap of 2D profiles. Here, we drew outlines around the outer contour of the GC nuclei rather than the somatic membrane since the nuclei were easier to identify due to their spotted appearance created by dark patches of heterochromatin. Similar to G(d) of the GC somata, G(d) of the GC nuclei had an asymmetrical shape negatively skewed (Figure 2B2), similar to that previously reported for rat hepatocyte nuclei in planar sections^54^. However, G(d) extended to 1 μm rather than 2 μm since smaller caps were easier to resolve in the TEM images compared to the confocal images. In total, we computed 26 G(d) of GC nuclei from 4 mice, 1–3 cerebellar sections per mouse, 6–7 TEM images per mouse. Comparison of G(d) within mice showed no significant differences (Kolmogorov-Smirnov, KS, test); hence, G(d) within mice were pooled. The resulting 4 nuclei G(d) spanned 1–6 μm with μ_d_ = 3.61–4.32 μm and σ_d_ = 0.85–0.98 μm (416–519 diameters per G(d)). To allow a comparison of GC somata and nuclei across species, described further below, we computed equivalent-area diameters of both the soma (d_soma_) and nucleus (d_nucleus_) of individual GCs using high-magnification TEM images of the same cerebellar sections of mice and found the d_soma_-versus-d_nucleus_ relation was well described by the following linear relation: d_soma_ = 0.952·d_nucleus_ + 1.016 μm (Pearson correlation coefficient (PCC) = 0.96, goodness-of-fit r_2_ = 0.92; n = 175 GCs from 4 mice).

Finally, to investigate G(d) when the section thickness is comparable to the size of the particles, we measured 2D diameters of synaptic vesicles in MFTs in high-magnification TEM images of the same cerebellar sections of mice, where T ≈ μ_D_ (Figure 2C1; Table 1). Because the vesicle membrane was not always apparent, we drew outlines around the outer contour of the vesicles rather than attempt to outline the inner or outer membrane leaflet. Moreover, we did not assume vesicle outlines were circular or oval, but rather followed the irregular contours of the vesicles that included membrane proteins, which are known to add at least 2 nm to the diameter of the vesicles^55^. Depending on the vesicle density and section thickness, vesicles aligned in the axial axis may show different degrees of overlap in the projection^54^ (Figure 1B). Although our TEM images of vesicles in MFTs exhibited numerous overlaps, this did not necessarily prevent outlining the vesicles since they were semi-transparent. Interestingly, G(d) of MFT vesicles were quite different to that of GC somata and nuclei, having a Gaussian shape with no negative skew and a large number of lost caps with d < 30 nm (Figure 2C2), similar to that previously reported for synaptic vesicles in thick sections^54,56^. In total, we computed 8 G(d) of MFT vesicles for 4 mice, 1 section per mouse, 2 MFTs per section, 152–428 vesicles per MFT. Comparison of the 8 G(d) showed the majority were significantly different from each other (KS test), even within mice comparisons, supporting previous findings of synapse-to-synapse variation of synaptic vesicle size^54,56^. Analysis of the 8 G(d) of MFT vesicles showed all had Gaussian shapes spanning 23–82 nm with μ_d_ = 43.1–47.3 nm and σ_d_ = 4.2–6.2 nm. These results highlight how the shape of G(d), especially the cap tail, depends on the relative size of the particles compared to the section thickness and the imaging method used to acquire the projections.

### 2.2 Exploration of the effects of section thickness and lost caps on G(d) of the Keiding model

To better understand how section thickness and lost caps affect the shape of G(d), we computed numerical solutions of the Keiding model (Equation 1) for different section thicknesses (T) and cap-angle limits (ϕ). To do this, we assumed F(d) was a Gaussian distribution (Equation 5) with normalised mean, i.e. a mean of one unit diameter (u.d.), and standard deviation of 0.09 u.d. to mimic the coefficient of variation of our experimental data (CV_D_ = σ_D_/μ_D_ ≈ 0.07 for GC somata, 0.08 for GC nuclei and 0.09 for MFT vesicles). For a planar section (T = 0 u.d.; Figure 1A) with no lost caps (ϕ = 0°), the numerical solution of G(d) had a skewed distribution with pronounced tail descending to 0 u.d. due to caps (where d < D; Figure 3A). In contrast, when the section thickness equaled the mean particle diameter (T = 1 u.d.; Figure 1B), G(d) had a larger peak and less pronounced tail, since most particles (57%) had their widest central region falling within the section (where d = D). To examine how G(d) is affected by lost caps, we computed the same numerical solutions for ϕ > 0°. When ϕ = 20°, the cap tails of G(d) now descended to 0.3 u.d. (Figure 3B), resembling those of the GC somata and nuclei G(d) in Figure 2A2 and B2. When ϕ = 40°, the cap tails of G(d) descended to 0.5 u.d. (Figure 3C). Interestingly, when ϕ = 70°, the cap tails of G(d) were no longer apparent and G(d) ≈ F(d) (Figure 3D). In fact, G(d) ≈ F(d) for ϕ ≥ 55°, in which case the two distributions were nearly indistinguishable (Figure 3E). The absence of cap tails in these G(d) is reminiscent of the G(d) for the MFT vesicles in Figure 2C2.

**Figure 3.**
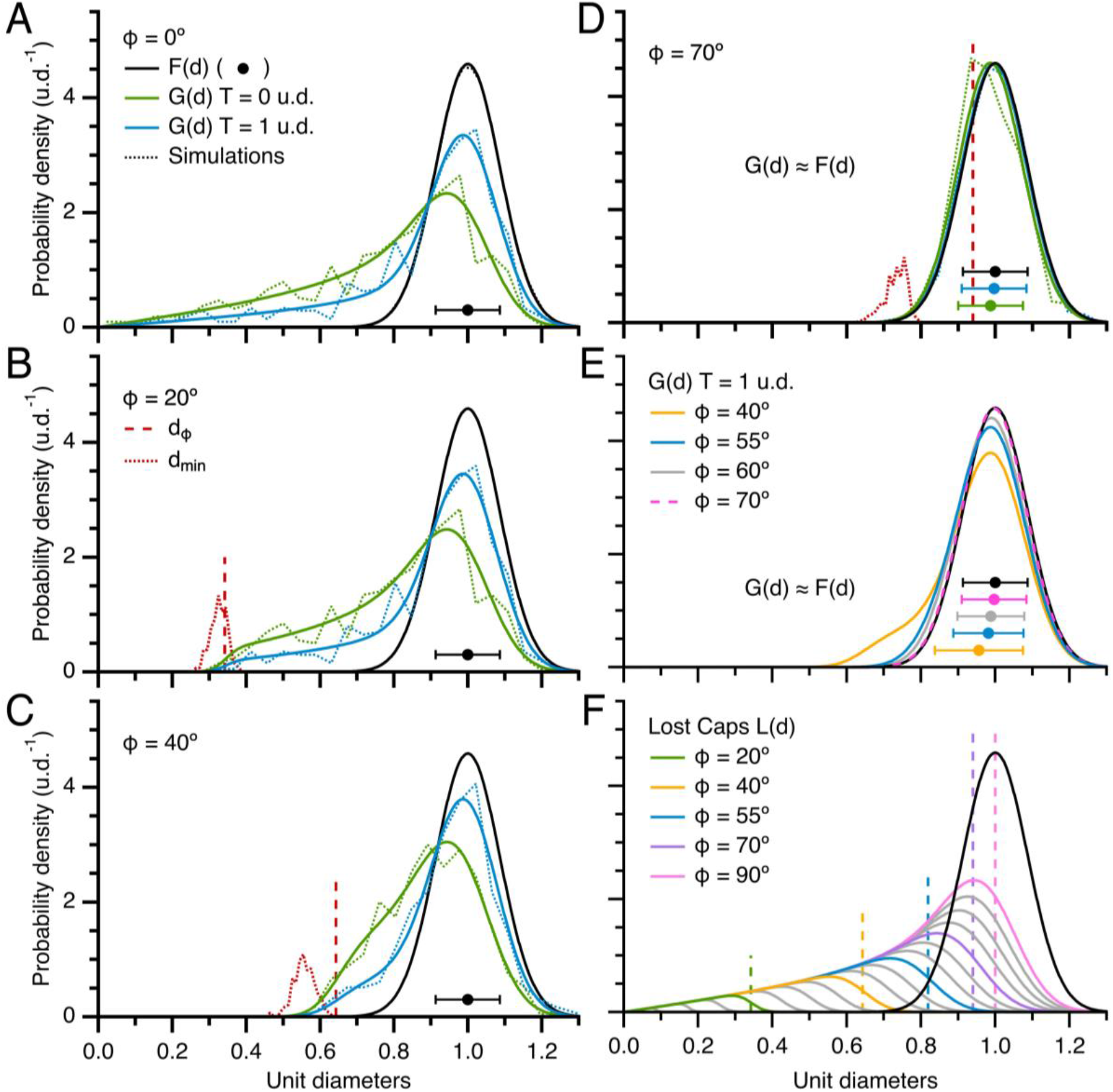
Effect of section thickness and lost caps on G(d). (A) Probability density of 2D diameters (G(d)) computed via Equation 1, where ϕ = 0° and the probability density of 3D diameters (F(d)) is a Gaussian distribution with normalised mean (Equation 5; μ_D_ ± σ_D_ = 1.00 ± 0.09 u.d.; black line and circle ± error bars). For T = 0 u.d. (green line) conditions are that of Wicksell’s model^16^ (Figure 1A) and for T = 1 u.d. (blue line) conditions are that of Bach’s model^38^ (Figure 1B). Because no caps are lost, both G(d) have tails extending to d = 0 u.d. Dotted lines denote G(d) of equivalent Monte Carlo simulations computed from ∼500 diameters using 0.04 u.d. bins. (B, C and D) Same as (A) for ϕ = 20, 40 and 70°. Here, the tails of G(d) are limited to 0.3, 0.5 and 0.7 u.d. For ϕ = 70°, G(d) ≈ F(d) (green and blue circles denote μ_d_ ± σ_d_) since most caps are lost. Comparison of the distribution of the minimum observed 2D diameter (d_min_, red dotted line, computed from simulations for both T = 0 and 1 u.d., probability densities scaled by 0.07) to d_ϕ_ = μ_D_·sinϕ (vertical red dashed line) shows d_min_ < d_ϕ_, especially at larger ϕ. (E) For ϕ ≥ 55°, G(d) ≈ F(d). (F) Distribution of lost caps, L(d), for ϕ = 5–90° in steps of 5° (gray and colored solid lines; Materials and Methods) compared to F(d) (black line). Vertical dashed lines denote d_ϕ_. Note, L(d, ϕ = 90°) = G(d, ϕ = 0°).

To quantify the relationship between G(d), F(d) and lost caps, we computed the distribution of lost caps (L(d)) for ϕ = 5–90° and compared L(d) to F(d) (Figure 3F). Results showed L(d) had a negatively skewed Gaussian shape with tail descending to 0 u.d. The upper limit of L(d) showed a dispersion, rather than a hard cutoff, since a variation in particle size in combination with a fixed-ϕ limit resulted in limiting caps of different size. For ϕ < 55°, there was a clear separation between L(d) and F(d), i.e. nearly all lost caps had a diameter smaller than those that define F(d). For ϕ > 55°, on the other hand, L(d) overlapped F(d), in which case there was a large number of lost caps with diameters equivalent to those that define F(d). At the most extreme condition where all caps are lost (i.e. ϕ = 90°) L(d) spanned from 0 to the largest diameter of F(d). Hence, at these larger ϕ, L(d) and F(d) cannot be delineated by diameter size.

### 2.3 Estimating the 3D particle size and cap-angle limit using the Keiding model

Next, we tested the Keiding model’s capacity to estimate F(d) and ϕ from G(d). To do this, we used a 3D Monte Carlo simulation package to compute virtual projections of spherical particles for planar and thick sections, creating lost caps according to the Keiding model (i.e. a cap was removed from the projection when its θ < ϕ; Figure S2; Materials and Methods). To make a direct comparison to our numerical solutions of Equation 1, we matched F(d) of the simulated particles to that used in the numerical solutions. Moreover, we set the number of measured 2D diameters per projection (∼500) to match the average sample size of our experimental G(d) of GC somata and nuclei, the two datasets with the largest number of measured diameters (Table 2). From the 2D diameters of the simulated particles, we computed G(d) for T = 0 and 1 u.d. and ϕ = 0, 20, 40 and 70°, all of which matched their equivalent numerical solution (Figure 3A–D). Using an LSE routine, we then curve fitted Equation 1 to the simulated G(d) and compared the resulting estimates of μ_D_, σ_D_ and ϕ to their true values. The comparison showed that, when true ϕ < 55°, the Keiding model accurately estimated μ_D_, σ_D_ and ϕ with only a small positive bias for σ_D_ (Figure 4A and C). On the other hand, when ϕ > 55°, estimates of μ_D_ and σ_D_ were less accurate, with positive and negative biases, respectively, and estimates of ϕ often had a large negative bias (Figure 4B and C). In this case, the LSE routine had difficulty estimating true ϕ since G(d) ≈ F(d) as ϕ approached 90°. The similarity between G(d) and F(d) at true ϕ > 55° was greatest for thick sections in which case fitting a simple Gaussian function (Equation 5) to G(d), which is equivalent to curve fitting the Keiding model with ϕ = 90°, where all caps are lost, resulted in similar estimates of μ_D_ and σ_D_, as did simply using the 2D measures μ_d_ and σ_d_ as estimates. Moreover, the estimates of μ_D_ and σ_D_ of the three approaches all showed relatively small biases (< 2 and 5%, respectively, for T = 1 u.d.). An overall comparison revealed that thick sections were moderately better for estimating μ_D_ and σ_D_, and planar sections were moderately better for estimating ϕ. Repeating the error analysis for G(d) computed from ∼2000 diameters gave qualitatively similar results, except μ_D_, σ_D_ and ϕ had smaller biases and confidence intervals, where the confidence intervals followed a 1/√n relation (Figure S5).

**Table 2.**
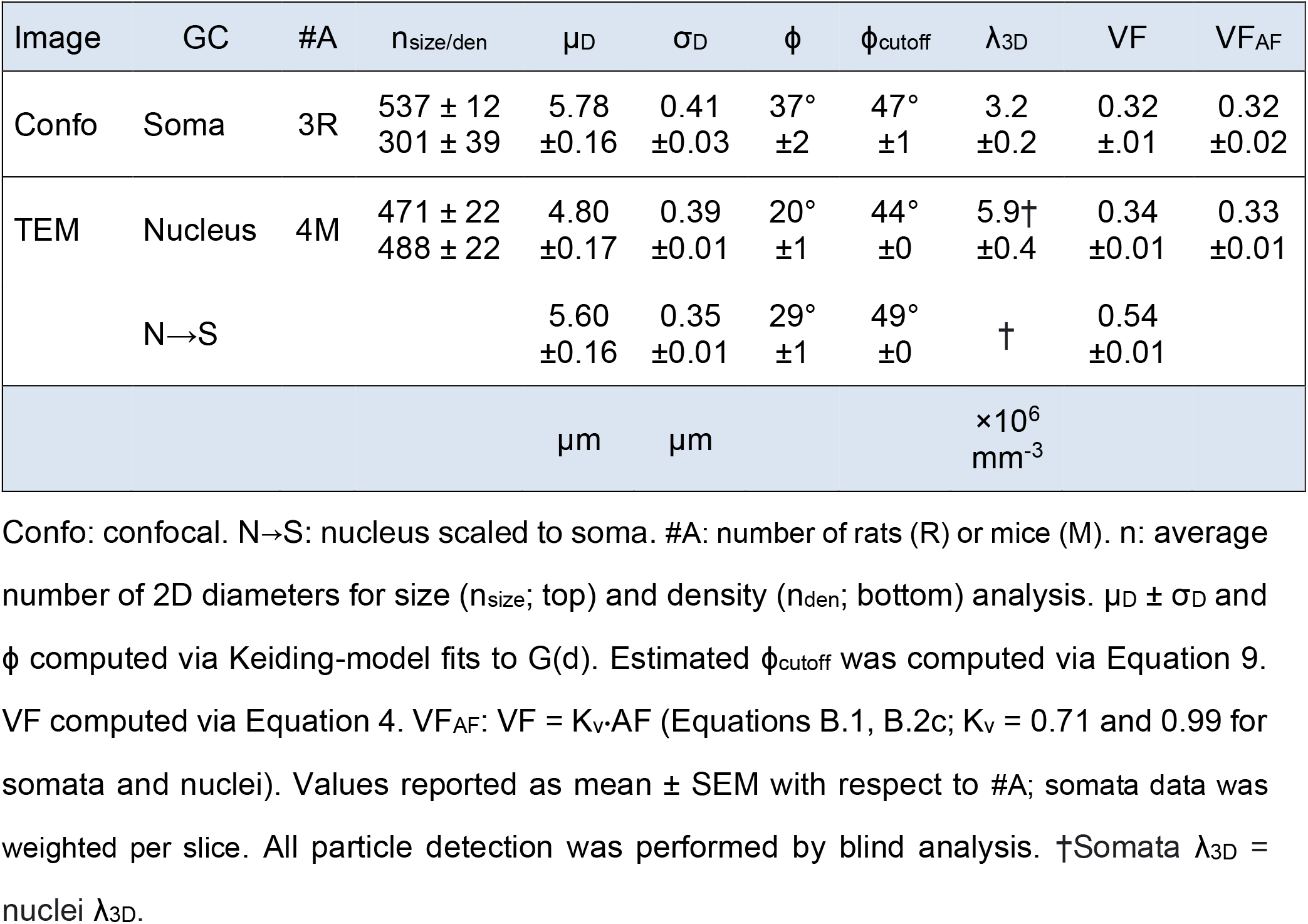
Summary of size and density analysis of cerebellar GC somata and nuclei.

**Figure 4.**
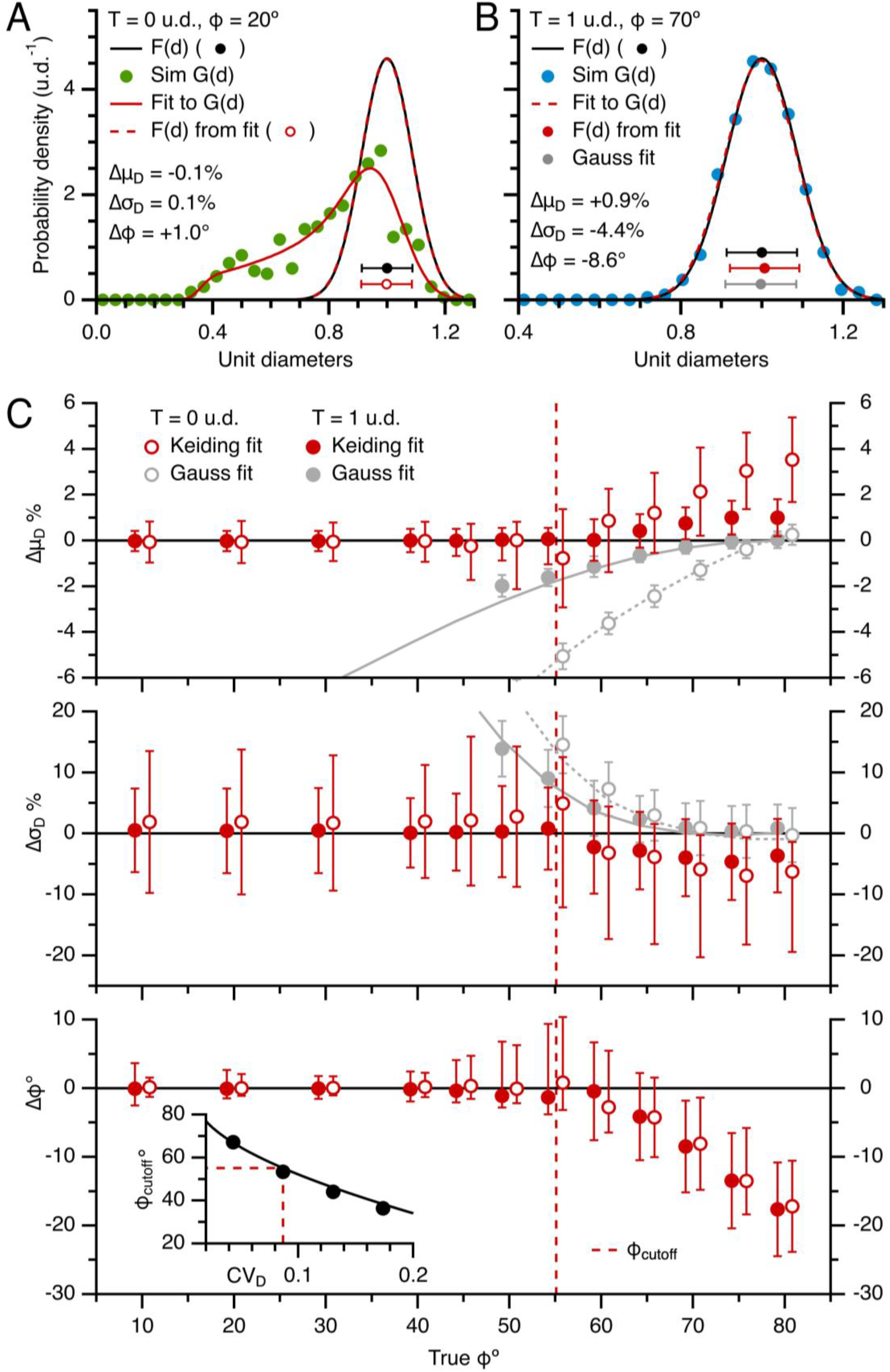
The Keiding model accurately estimates F(d) and ϕ from G(d) for true ϕ < ϕ_cutoff_ (Part I. simulations). (A) Curve fit of Equation 1 (red solid line) to G(d) of the simulation in Figure 3B where T = 0 u.d. and ϕ = 20° (green circles; ∼500 diameters). F(d) derived from the fit (red dashed line and circle) matches the true simulation F(d) (black line and circle) and fit ϕ matches true ϕ. (B) Same as (A) for G(d) of the simulation in Figure 3D where T = 1 u.d. and ϕ = 70° (blue circles). Although there is a good match between estimated and true F(d), estimation errors Δμ_D_ and Δσ_D_ are larger than those in (A) and estimated ϕ < true ϕ. (C) Average estimation errors Δμ_D_, Δσ_D_ and Δϕ of Keiding-model fits to simulated G(d), as in (A) and (B), for true ϕ = 10–80°, T = 0 and 1 u.d., CV_D_ = 0.09 (red open and closed circles; μ_Δ_ ± σ_Δ_ for 100 repetitions per ϕ). Red dashed lines denote ϕ_cutoff_ (∼55°; Equation 8) above which G(d) ≈ F(d) and true ϕ becomes indeterminable. For comparison, results are shown for Gaussian fits to the same G(d) (Equation 5; gray circles) and 2D statistics μ_d_ ± σ_d_ (gray lines). Data shifted ±0.8° to avoid overlap. Asymmetrical error bars indicate skewed distributions (Materials and Methods). Inset: ϕ_cutoff_ vs. CV_D_ for simulations (black circles) and Equation 8 (black line; n = 500 diameters).

While the above results show the Keiding model works well in estimating μ_D_, σ_D_ and ϕ, estimates were most accurate when ϕ < 55° (Figure 4C). However, this upper limit of ϕ (ϕ_cutoff_) is dependent on the spread of F(d) and the number of measured diameters, both of which were set in our Monte Carlo simulations to match our experimental data (CV_D_ = 0.09, ∼500 diameters). For a wider F(d) and/or smaller number of diameters, accurate parameter estimation will be limited to a smaller range of true ϕ, and vice versa (Figures 4C and S5, insets). To quantify this effect, we computed ϕ_cutoff_ over a range of CV_D_ (0.04–0.17) and number of diameters (n ≈ 200–2000) for our Monte Carlo simulations (T = 0 u.d.; Materials and Methods). Results showed sinϕ_cutoff_ fit well to a bivariate polynomial with respect to CV_D_ and 1/√n (Equation 8). Next, we investigated whether this ϕ_cutoff_ equation could be used to test the accuracy of estimated ϕ using estimated μ_D_ and σ_D_, since the estimation errors of μ_D_ and σ_D_ are relatively small even when true ϕ > ϕ_cutoff_ (Figures 4C and S5). Results showed that indeed this was possible with relatively small adjustments to the sinϕ_cutoff_ relations (Equation 9). Hence, if one requires an accurate estimate of ϕ, for example when computing particle density as discussed further below, and one only has estimates of μ_D_ and σ_D_, then Equation 9 can be used as an accuracy test, i.e. by requiring estimated ϕ < estimated ϕ_cutoff_ (Figure S6). Although the fit error of ϕ can also be used as a measure of accuracy, ∼10% of our curve fits to simulated G(d) for T = 0 u.d. and true ϕ > 55° showed small fit errors (< 1°) but large estimation errors (|Δϕ| > 5°). Hence, a combination of the fit error and Equation 9 can be used as an accuracy test for ϕ.

### 2.4 Overlapping particle projections in thick sections

A potential source of error of the Bach^38^ and Keiding^49^ models for thick sections is that the models assume particles near the bottom of the section can be identified and outlined in a projection as well as those toward the top of the section. While this may be true for thin sections, it is unlikely to be true for thick sections with overlapping particle projections^2,57^. To investigate what effects overlapping projections might have on the ability to accurately estimate F(d) and ϕ from G(d), we used simulations to compute the sum of 2D projection overlaps between a given particle and particles higher in a thick section, where particles had a random distribution and a modest to high VF (0.15–0.45). Results of the analysis showed that, as expected, particles toward the bottom of the section experienced more overlaps in a projection, and this effect was greater for thicker sections and higher particle density (Figure 5A). To investigate what effect projection overlaps might have on the shape of G(d), we set an upper limit to the amount of projection overlaps an observable particle can have, i.e. particles with overlaps above the set limit (ψ) were considered hidden from view, i.e. lost, and excluded from the projection. Results of these ‘semi-transparent’ particle simulations showed that, for thick sections and high particle density, a large number of particles near the bottom of the section were excluded from the projection, thereby reducing the effective section thickness (T) for a given projection (Figure 5B; ψ = 0.25). Nevertheless, excluding bottom-dwelling particles from the projection had minimal effect on the shape of G(d), and therefore had little effect on estimates of F(d), as long as the top-dwelling particles were simulated as transparent, i.e. small degrees of projection overlap did not interfere with counting the projections or computing their size (Figure 5C). This is in contrast to when particles were simulated as opaque such that overlapping projections reduced the number of observed projections and increased their size; in this case, opaque particles created a positive skew in G(d) (Figure 5D). However, the distortions in G(d) produced only modest changes in estimates of F(d). Hence, these results indicate overlapping projections of semi-transparent and opaque particles in thick sections create only small biases in estimates of F(d). However, the overlapping projections do have the potential to reduce the effective section thickness (Figure 5B) and therefore affect the estimates of the 3D density (see below).

**Figure 5.**
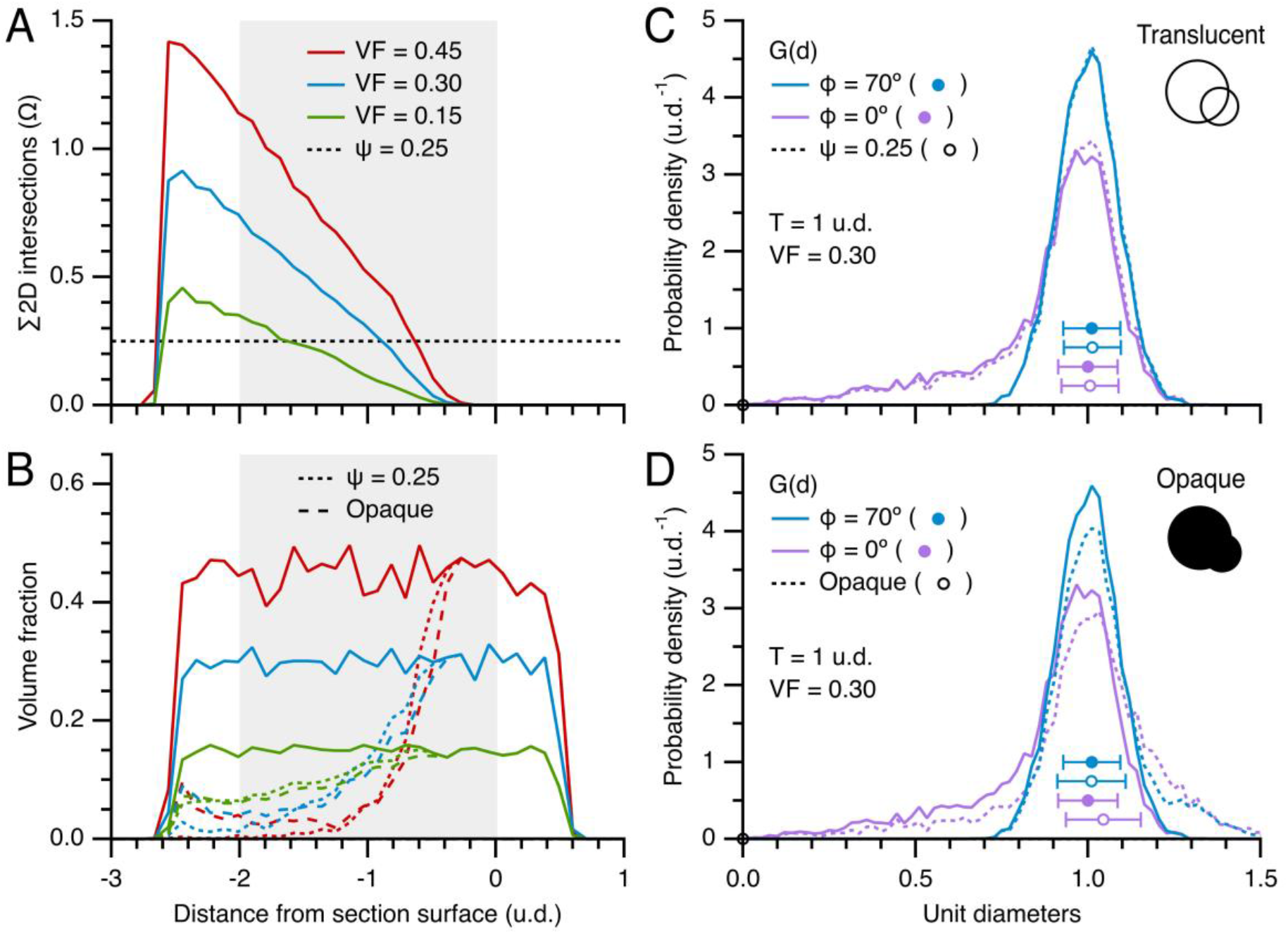
Effects of projection overlaps for thick sections. (A) Sum of 2D projection overlaps (Ω) as a function of distance from the surface of a simulated section for particle VF = 0.15, 0.30 and 0.45 (green, blue and red lines) where Ω > 1 indicates a particle’s projection is likely to be completely overlapping with projections of other particles closer to the section surface. Histograms were computed using particle z-center points and 0.1 u.d. bins. Gray background denotes section thickness (T = 2 u.d.). The distributions extend 0.5 u.d. below the section because of caps (Figure 1B). Black dotted line denotes upper limit ψ = 0.25 for (B) and (C). For simplicity, ϕ = 0°. (B) VF of those particles appearing in a simulated projection as a function of distance from the surface of each section (T = 2 u.d.) for transparent particles (solid lines; control, ψ = ∞), semi-transparent particles (dotted lines; a particle is removed from the projection if its Ω > 0.25) and opaque particles (dashed lines; bottom-dwelling particles are merged with top-dwelling particles if -1 < α < 0; Materials and Methods). VF = 0.15, 0.30 and 0.45 as in (A). The effect of semi-transparent and opaque particles is to reduce the effective T. Histograms were computed using particle z-center points and 0.1 u.d. bins; counts were converted to VF using the equivalent bin volume (bin z-width times geometry Area_xy_) and particle diameter distribution. For simplicity, ϕ = 0°. (C) Comparison of simulated probability density of 2D diameters (G(d)) for transparent and semi-transparent particles (ψ = ∞ and 0.25; solid vs. dotted lines) shows little difference for two extreme conditions of lost caps (ϕ = 0° and 70°; purple and blue). In these simulations, projection overlaps did not affect estimates of their size (inset). Circles and error bars denote Keiding-model fit parameters μ_D_ ± σ_D_. T = 1 u.d. and VF = 0.30. (D) Comparison of simulated G(d) for transparent and opaque particles (solid vs. dotted lines). For opaque particles, overlapping projections with -1 < α < 0 were treated as a single projection with larger area (inset), thereby creating a positive skew in G(d). For (A)–(D), average histograms were computed from 20 sections, ∼500 particles per section.

### 2.5 Validation of the Keiding model using 3D reconstructions (vesicle size)

In the previous section, we used Monte Carlo simulations to test the Keiding model’s capacity to estimate F(d) and ϕ from G(d). However, while the simulations included variation (i.e. stochasticity) in particle location and size, they lacked variation normally associated with experimental data such as that due to finite spatial resolution, limited contrast, irregular particle shape and blending with surrounding material (e.g. intracellular/extracellular proteins). Hence, to address this shortcoming, we tested the accuracy of the Keiding model using a volumetric ET z-stack of clusters of MFT vesicles, since this allowed us to directly measure F(d) and ϕ, as well as G(d). First, F(d) was measured by outlining individual vesicles across multiple planes of the z-stack (Figure 6A; ET11). From the outlines of a given vesicle, equivalent radii were computed as a function of z-depth and fit to a circle (Equation 10), resulting in estimates for the vesicle’s 3D diameter (D) and z-axis center point (z_0_) (Figure S7). Aligning the vesicle profiles at their centers revealed the profiles overlaid each other and fit well to a semi-circle (Figure 6B). From the measured 3D diameters, we computed μ_D_ ± σ_D_ = 42.7 ± 3.4 nm and F(d) (Figure 6D; n = 234). To determine the shape of F(d), we fit the distribution to Gaussian, chi and gamma functions (Equations 5, 6, 7) and found closely overlapping fits (Figure S8). Moreover, we found parameter *f* > 70 for the chi and gamma fits indicating Gaussian-like distributions. Repeating the same analysis for F(d) derived from another ET z-stack (ET10) and another study of MFT vesicles^5^ produced similar results. Hence, F(d) of the MFT vesicles was well described by a Gaussian function, which is consistent with the findings of a 3D analysis of vesicles in hippocampal CA1 excitatory synapses^58^.

**Figure 6.**
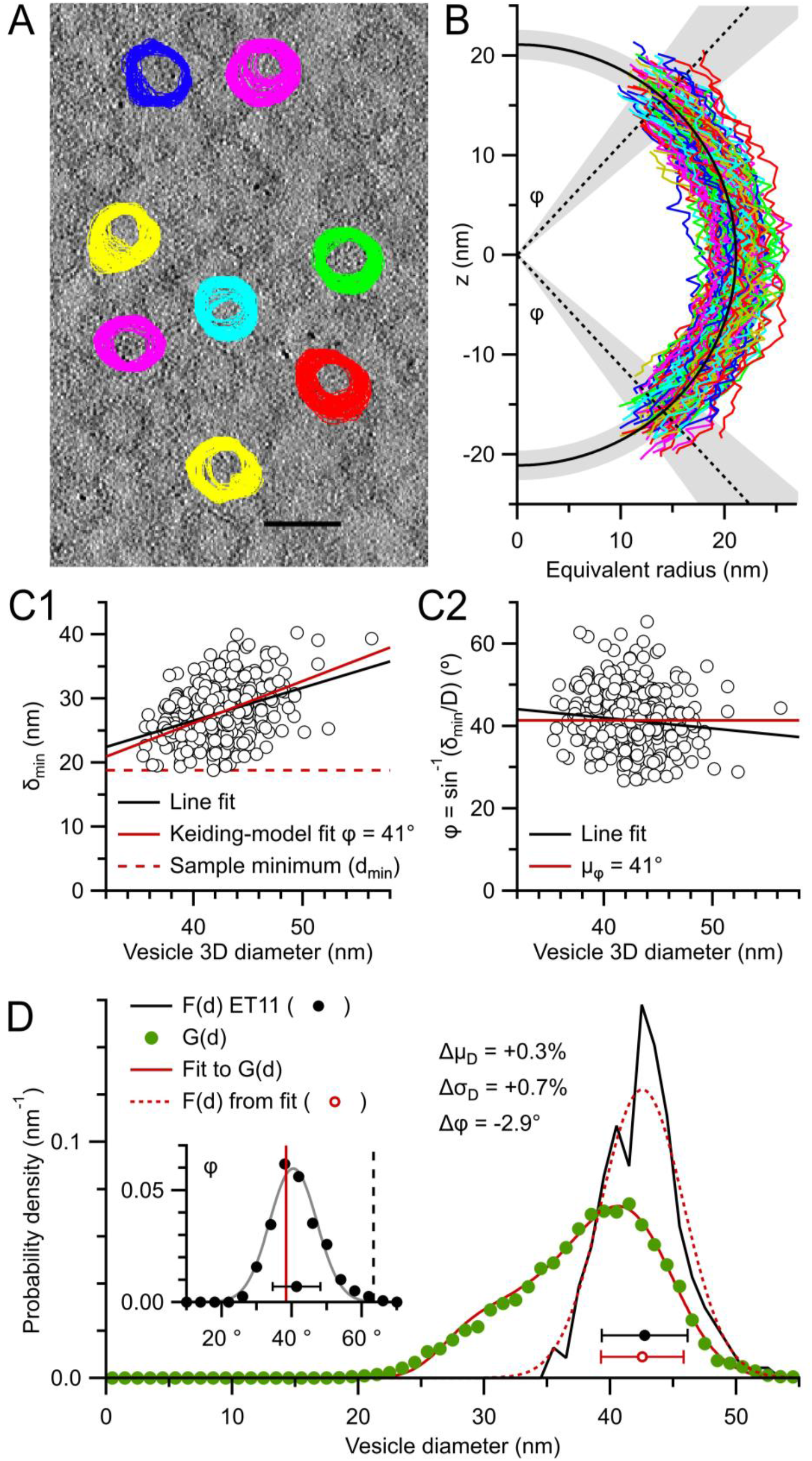
The Keiding model accurately estimates F(d) and ϕ from G(d) for true ϕ < ϕ_cutoff_ (Part III. vesicles ET11). (A) One of 261 serial images of a 3D ET reconstruction (ET11) of a cerebellar MFT section 133 nm thick. 271 vesicles, including 108 caps, were tracked and outlined through multiple z-planes and their d_area_ computed as a function of z-plane number. This image shows outlines for 8 representative vesicles, overlaid with outlines from images above and below. Scale bar 50 nm. (B) Vesicle radius (½d_area_) vs. z-depth (colored lines) with vesicle centers (z_0_) aligned at z = 0; caps are not displayed. Black semi-circle and shading denote μ_D_ ± σ_D_ = 42.7 ± 3.4 nm for all measured 3D diameters (D; n = 234). Black dotted lines and shading denote measured ϕ: μ_ϕ_ ± σ_ϕ_ = 41 ± 7° (n = 397, measures from both poles). Parameters z_0_ and D were estimated via a curve fit to a circle and ϕ = sin^−1^(δ_min_/D), where δ_min_ is the minimum d_area_ at a given pole (Figure S7A). It was not possible to curve fit the smallest caps. (C1) Minimum 2D diameter of a given vesicle (δ_min_) vs. 3D diameter (D, circles; n = 397) with line fit (black line; χ^2^ = 5315, PCC = 0.4, r^2^ = 0.2) and Keiding-model fit (red solid line; δ_min_ = D·sinϕ; fit ϕ = 40.9 ± 0.2°; χ^2^ = 5394, PCC = 0.4, r^2^ = 0.3). The smallest δ_min_ (d_min_; red dashed line) is a poor match to the data. (C2) Same as (C1) but for ϕ = sin^−1^(δ_min_/D) with line fit (black line; PCC = -0.1, r^2^ = 0.01) and μ_ϕ_ = 41.4° (red solid line). (D) Measured F(d) (black line and circle; 1 nm bins) vs. G(d) (green circles; n = 13,914 outlines; 1 nm bins). A curve fit of Equation 1 to G(d) (red solid line; μ_D_ = 42.9 ± 0.1 nm, σ_D_ = 3.4 ± 0.1 nm, ϕ = 38.4 ± 0.3°; T fixed to 0 nm) resulted in estimated F(d) (red dotted line and circle) nearly the same as measured F(d). Inset: probability density of measured ϕ in (C2) (black circles; probability per degree) with Gaussian fit (gray line; Equation 5; μ_ϕ_ ± σ_ϕ_ = 40.4 ± 6.7°), fit ϕ (red line) and ϕ_cutoff_ (black dashed line; ∼63°; Equation 8).

Next, we measured ϕ by computing the axial extent of each vesicle profile, which only partially extended to the north and south poles (Figures 6B and S7A). The missing profiles at the pole regions, i.e. lost caps, are likely due to limited resolution and poor contrast of orthogonally oriented membranes^45^. Plotting the minimum measured diameter (δ_min_) for the vesicle poles interior to the z-stack versus their 3D diameter revealed a linear relation that was well described by the Keiding model, i.e. δ_min_ = D·sinϕ (Figure 6C1). In contrast, the minimum of all measured diameters (d_min_), which has been used as a correction for lost caps (Appendix A), was a poor fit to the δ_min_-D relation. Converting all δ_min_ to ϕ values revealed a distribution that spanned 27–65° (Figure 6C2) with μ_ϕ_ ± σ_ϕ_ = 41.4 ± 6.8° (CV_ϕ_ = 0.2; n = 397) and was well described by a Gaussian distribution (Figure 6D, inset). Hence, this analysis revealed a source of variation not accounted for in the Keiding model, which assumes all particles have the same ϕ. The ramifications of a variation in ϕ are explored in the next section.

Next, we computed G(d) from all measured 2D diameters (Figure 6D). Comparison of G(d) to F(d) revealed a negative skew in G(d) as expected for planar sections (estimated T ≈ 0.1 u.d.; Table 1). Finally, using the measured G(d), F(d) and ϕ, we tested the Keiding model by curve fitting Equation 1 to G(d), assuming a Gaussian F(d) (Equation 5), which our results above confirmed is a good assumption, and comparing the resulting estimated F(d) and ϕ to their measured ‘true’ values. Results showed the estimated F(d) (μ_D_ ± σ_D_ = 42.9 ± 3.4 nm) closely matched the measured F(d) (Δμ_D_ = +0.3% and Δσ_D_ = +0.7%) and the estimated ϕ (38.4°) closely matched μ_ϕ_ (Δϕ = -2.9°). Moreover, both the measured and estimated ϕ were below ϕ_cutoff_ (∼63°; Table 3). Repeating the same 2D versus 3D analysis for another ET z-stack produced similar results (Figure S9; ET10; Δμ_D_ = +1.3% and Δσ_D_ = -4.9%, Δϕ = -0.4°). Hence, this 3D analysis shows that, even with the added variability from experimental data, including variability in ϕ, the Keiding model accurately estimated F(d) and ϕ from G(d) with only small error when true ϕ < ϕ_cutoff_, confirming our previous results from simulations (Figures 4C, S5 and S6).

**Table 3.**
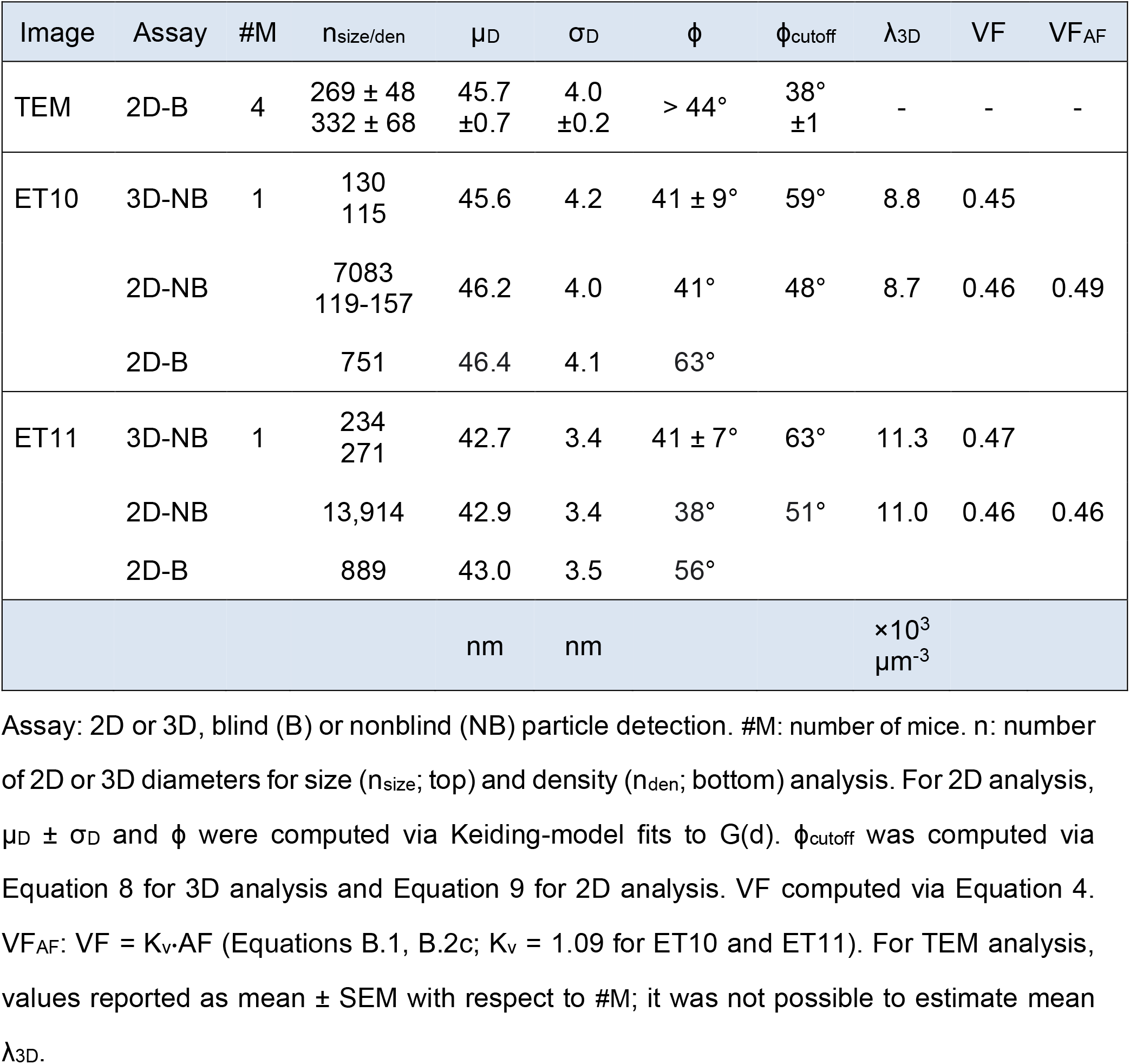
Summary of size and density analysis of cerebellar MFT vesicles.

### 2.6 Exploration of the Keiding model’s fixed-ϕ assumption

Despite the impressive capacity of the Keiding model to estimate F(d) and ϕ of our experimental data, close inspection of the curve fits to G(d) measured from our ET z-stacks showed systematic deviation at the smaller diameters (d = 20–40 nm; Figures 6D, S9D and S10A) and Δμ_D_ and Δϕ were larger than those computed from simulations (Figure 4C). To test whether the discrepancies arose from the variation of ϕ within the sample of vesicles (Figures 6C2 and S9C2), we used Monte Carlo simulations to replicate the z-stack analysis in Figure S9 and compared the resulting G(d) to the experimental G(d) for a fixed-ϕ model where all vesicles have the same ϕ, as in the Keiding model, and a Gaussian-ϕ model where ϕ of each vesicle is randomly drawn from a Gaussian distribution (μ_ϕ_ ± σ_ϕ_) whose shape and variation matched the measured ϕ (CV_ϕ_ = 0.2). Comparison of the two models confirmed that the small discrepancy between the Keiding-model fit and experimental G(d) at the smaller diameters can indeed be explained by the variation in ϕ (Figure S10B1 and B2). Moreover, the simulations allowed us to quantify the errors from assuming a fixed ϕ in our Keiding-model curve-fit routine, giving biases Δμ_D_ = +0.3%, Δσ_D_ = -2.5% and Δϕ = -1° (Figure S10C). Hence, the Keiding-model fixed-ϕ assumption only introduced small biases into the estimates of μ_D_, σ_D_ and ϕ. To further explore the effects of assuming a fixed ϕ in the Keiding model, we computed errors Δμ_D_, Δσ_D_ and Δϕ from Keiding-model fits to simulated G(d), as just described, over a range of ϕ distributions (μ_ϕ_ = 10–50°; CV_ϕ_ = 0.2) for planar and thick sections (T = 0 and 1 u.d.; Figure S11). Results gave negligible estimation errors, except for conditions of a planar section and μ_ϕ_ = 40–50°, in which case errors showed modest biases. Hence, these results show that the Keiding-model fixed-ϕ assumption introduces only small errors when estimating F(d) and ϕ from G(d).

### 2.7 Impact of blind versus nonblind particle detection in 2D images

For the size analysis of MFT vesicles in our 3D reconstructions, G(d) was computed from measurements of vesicles that were tracked through multiple images of the ET z-stack. This ‘nonblind’ approach provided information that made it easier to identify small caps, resulting in a tail of G(d) that descended to 20 nm and ϕ = 41° (Figures 6D and S9D). In contrast, G(d) of MFT vesicles in the first Results section were computed ‘blind’ without knowledge of a vesicle’s 3D position in a z-stack. In this case, G(d) had a symmetrical Gaussian-like appearance with no tail (Figure 2C2). Hence, we wondered if the difference between the two G(d) could simply be explained by one analysis being blind and the other nonblind. To test this, we recomputed G(d) from our ET z-stacks using a blind analysis: images of the z-stack were analysed in random order, and vesicles were not tracked between adjacent images. Results showed G(d) computed blind had a symmetrical Gaussian appearance with no tail, and a similar appearance to F(d) (Figure 7A and B), thus confirming the difference in G(d) can be explained by a greater number of lost caps in the blind analysis. To quantify ϕ for the blind analysis, we simultaneously fit the Keiding model to G(d) computed blind and nonblind, sharing parameters for F(d), i.e. μ_D_ and σ_D_. For ET11, results gave ϕ = 38 ± 1° for the nonblind analysis (similar to the analysis in Figure 6D) and ϕ = 56 ± 1° for the blind analysis. Hence, there was a 17° difference (bias) in ϕ. For ET10, results gave a 22° difference in ϕ. Interestingly, despite a large estimated ϕ for both blind analysis (where ϕ > ϕ_cutoff_), which usually correlates with a large fit error (Figure 4C), the fit errors for this analysis were only ±1°. The small fit errors are due to the sharing of parameters μ_D_ and σ_D_ during the simultaneous fit, in which case good estimates of μ_D_ and σ_D_ were achieved via G(d) computed nonblind, and ϕ was determinable for both blind and nonblind analysis. Besides the difference in ϕ, the simultaneous fit also revealed G(d) ≈ F(d) for the blind analysis, with only a subtle difference between the two curves. These results highlight the difference between blind and nonblind particle detection, and that cap detection can be significantly improved by additional 3D information provided by a z-stack.

**Figure 7.**
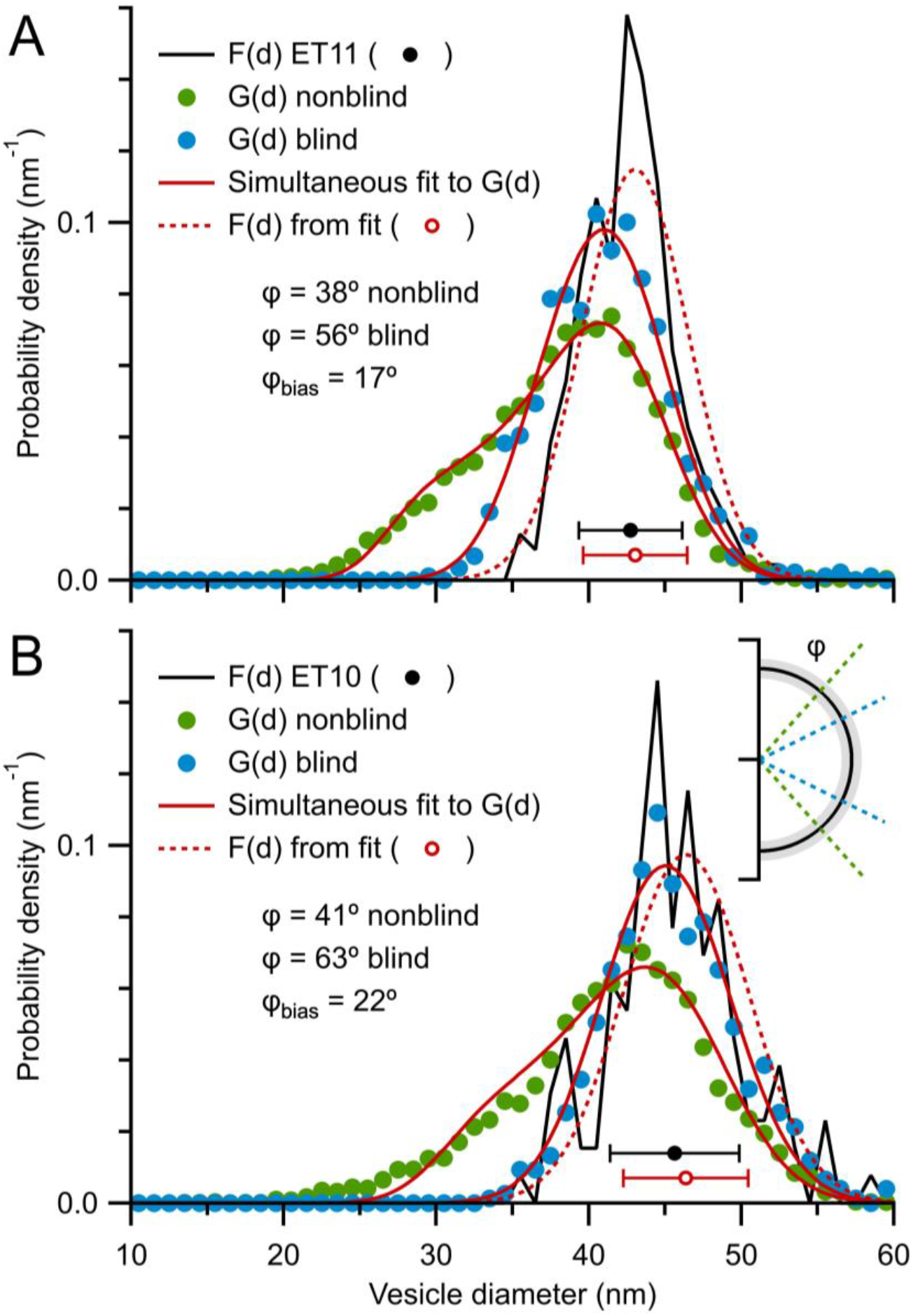
Comparison of G(d) computed blind versus nonblind reveals a large bias in particle cap detection (ϕ_bias_ ≈ 20°). (A) Probability density of 2D diameters (G(d)) computed via a nonblind vesicle detection (green circles; ET11; Figure 6D) versus a blind vesicle detection (blue circles). Both G(d) were simultaneously curve fitted to Equation 1, where parameters μ_D_ and σ_D_ were shared, revealing a 17° bias in ϕ (red solid lines; μ_D_ = 43.0 ± 0.1 nm, σ_D_ = 3.5 ± 0.1 nm, nonblind ϕ = 38.3 ± 0.5°, blind ϕ = 55.5 ± 0.7°). As in Figure 6D, estimated F(d) (red dotted line and circle) is similar to measured F(d) (black solid line and circle). For the blind analysis, there were 889 diameters measured from 18 z-planes (z_#_ = 1–260) spaced 5–15 nm apart, analysed in random order. (B) Same as (A) for ET10 (Figure S9D). The simultaneous curve fit revealed a 22° bias in ϕ (red solid lines; μ_D_ = 46.4 ± 0.1 nm, σ_D_ = 4.1 ± 0.1 nm, nonblind ϕ = 40.9 ± 0.7°, blind ϕ = 63.3 ± 1.3°). For the blind analysis, there were 751 diameters measured from 30 z-planes (z_#_ = 51–236) spaced 2–7 nm apart, analysed in random order. Inset cartoon depicts the bias in ϕ along the axial axis of a vesicle.

### 2.8 Estimation of the 3D size of granule cell somata and nuclei

Having shown the Keiding model accurately estimates F(d) and ϕ from G(d), with the exception that ϕ is indeterminable when true ϕ > ϕ_cutoff_ (Figure 4C), we curve fitted Equation 1 to the 9 G(d) computed from GC somata of rats (Figure 2A2), resulting in estimated μ_D_ = 5.48–6.32 μm, σ_D_ = 0.34–0.49 μm and ϕ = 27–52° with small fit errors of 1–2° (Figures 8A, C and S12). The ϕ-accuracy test computed via Equation 9 showed estimated ϕ < estimated ϕ_cutoff_ (∼44–50°) for all but two fits where estimated ϕ was ∼3–4° above its estimated ϕ_cutoff_. For the two fits that failed the ϕ-accuracy test, we recomputed the fits while fixing their ϕ to that estimated from the other fits of the same preparation (this made little difference in estimates of μ_D_ and σ_D_). Averaging results across the 3 rats gave μ_D_ = 5.78 ± 0.16 μm, σ_D_ = 0.41 ± 0.03 μm and ϕ = 37 ± 2° (±SEM; Table 2). Simulations indicate these estimates have negligible biases and small confidence intervals (Δμ_D_ = 0.0 ± 0.7 %, Δσ_D_ = +1.5 ± 8.4 %, Δϕ = +0.1 ± 1.3° for T = 0.3 u.d., true ϕ = 37°, ∼500 diameters).

**Figure 8.**
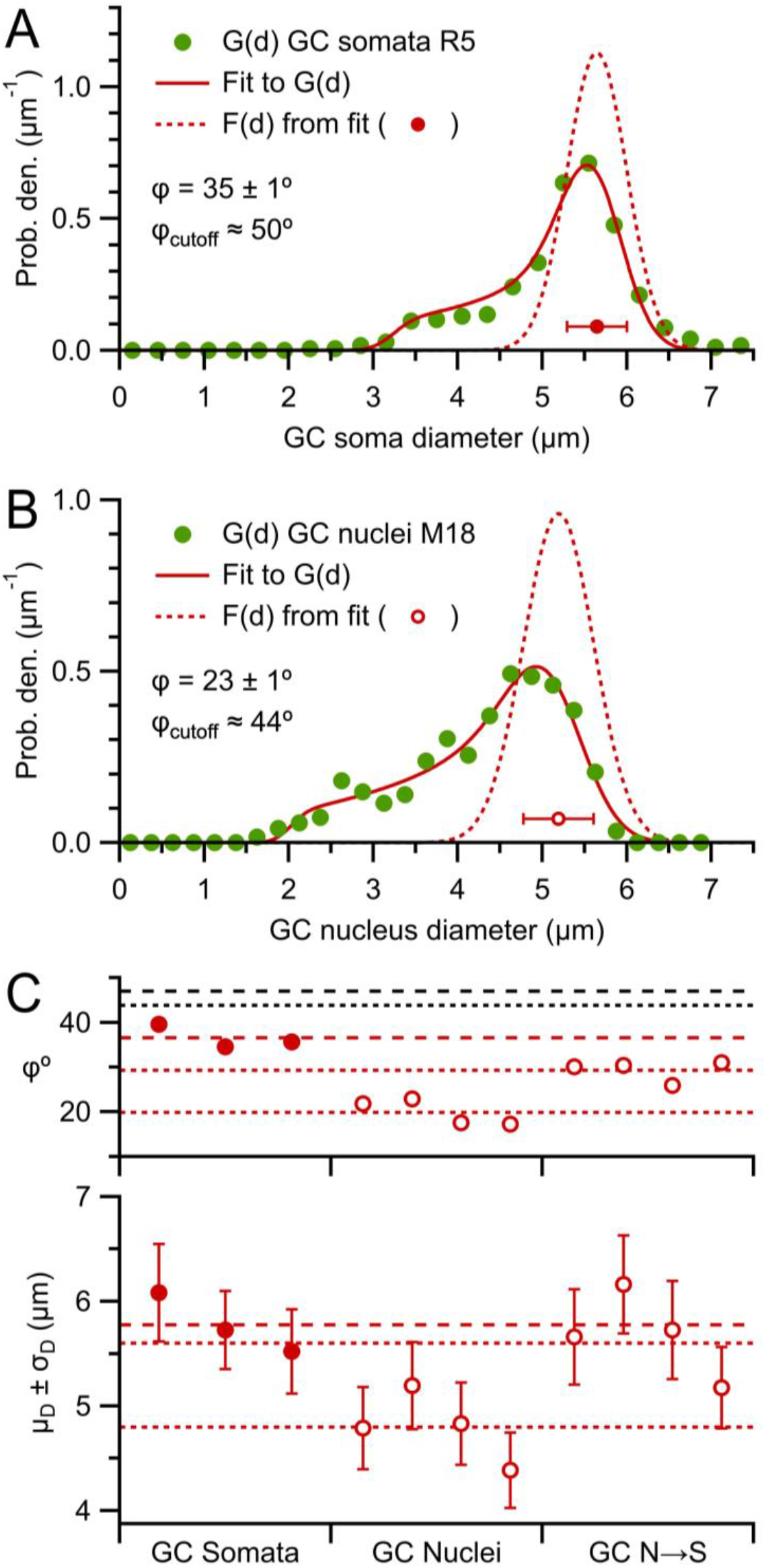
Estimates of F(d) and ϕ from G(d) of cerebellar GC somata and nuclei. (A) Curve fit of Equation 1 (red solid line) to the G(d) of rat GC somata in Figure 2A2 (green circles) resulting in estimates for F(d) (red dotted line and circle), ϕ and ϕ_cutoff_ (Equation 9). (B) Same as (A) for the G(d) of mouse GC nuclei in Figure 2B2. (C) Parameters μ_D_ ± σ_D_ (bottom) and ϕ (top) from Keiding-model fits to G(d) of GC somata for 3 rats (red closed circles; Figure S12; weighted averages with respect to number of tissue sections), G(d) of GC nuclei for 4 mice (red open circles; Figure S13; pooled diameters from 6–7 TEM images per mouse) and the same G(d) of GC nuclei scaled to somata dimensions (N→S). Red dashed and dotted lines denote averages across the 3 rats and 4 mice. Black dashed and dotted lines denote average estimated ϕ_cutoff_ for somata and nuclei (∼47 and 44°).

Next, we curve fitted the Keiding model to the 4 G(d) computed from GC nuclei of mice (Figure 2B2), resulting in μ_D_ = 4.39–5.19 μm, σ_D_ = 0.36–0.42 μm and ϕ = 17–23° with small fit errors of 1–2° (Figures 8B, C and S13) and all ϕ passing the ϕ-accuracy test (i.e. estimated ϕ < estimated ϕ_cutoff_, where estimated ϕ_cutoff_ ≈ 43–44°; Equation 9). The smaller less-variable ϕ for the GC nuclei compared to the GC somata reflects a more complete G(d) with fewer lost caps, achieved via the higher resolution and contrast of the TEM compared to confocal images. Moreover, since the nuclei were much larger than the section thickness, there were no overlaps of the nuclei in the TEM images and therefore less ambiguity of where the borders lay between nuclei. Across the 4 mice, μ_D_ = 4.80 ± 0.17 μm, σ_D_ = 0.39 ± 0.01 μm and ϕ = 20 ± 1° (±SEM), which are similar to estimates computed from an analysis where all G(d) are aligned and pooled into a single G(d) (Figure S14C). Simulations indicate these estimates have negligible biases and small confidence intervals (Figure 4C; Δμ_D_ = -0.1 ± 0.9 %, Δσ_D_ = +1.9 ± 11.9 %, Δϕ = 0.0 ± 1.6° for T = 0 u.d., true ϕ = 20°, ∼500 diameters).

To compare the above estimates for GC size between rats and mice, we scaled G(d) of the GC nuclei of mice via the d_soma_-versus-d_nucleus_ linear relation described above and curve fitted the Keiding model to the new G(d). Results gave μ_D_ = 5.60 ± 0.16 μm and σ_D_ = 0.35 ± 0.01 μm (±SEM), which are similar to that estimated for rats (p = 0.49 and 0.07, respectively; *t*-test). These results are consistent with our simulations that showed overlapping projections of opaque particles (a likely scenario for the rat confocal dataset) create negligible bias in estimates of μ_D_ (Figure 5D). Hence, these results are consistent with F(d) being the same for GCs in rats and mice. The results also indicate the two different tissue preparations of the rat and mouse datasets (chemical versus cryo fixation) had little effect on the size of GCs.

### 2.9 Estimation of the 3D size of vesicles in mossy fiber terminals

To estimate the 3D diameter of MFT vesicles, we curve fitted Equation 1 to the 8 G(d) computed from MFT vesicles of mice (Figure 2C2), resulting in estimated μ_D_ = 42.4–47.2 nm, σ_D_ = 3.5–5.0 nm and ϕ = 45–70° with large fit errors most of which exceeded 80° (Figures 9 and S15). Here, the large estimated ϕ with large fit errors indicate ϕ was indeterminable (Figure 4C). This conclusion was further supported by the finding that all estimated ϕ failed the ϕ-accuracy test (i.e. estimated ϕ > estimated ϕ_cutoff_, where ϕ_cutoff_ ≈ 34–42°). While these estimates of ϕ are larger than that computed from our ET z-stack analysis (Figures 6D and S9D; ϕ = 41°), this is expected for the G(d) analysed here given the greater difficulty in identifying vesicle caps using a blind analysis (Figure 7) and thick sections (T ≈ μ_D_). While the large estimates of ϕ with large errors means there may be small biases in these final estimates of μ_D_ and σ_D_ (absolute values < 1 and 5%, respectively; Figure 4C), the estimates are not significantly different to the 3D measures computed from our two ET z-stacks (p = 0.3 and 0.6, respectively; Student’s *t*-test). Again, the similarity in estimates of F(d) between our datasets indicates differences in tissue preparation (cryo versus chemical fixation) had little effect on the size of synaptic vesicles, which is consistent with other studies^31,59^.

**Figure 9.**
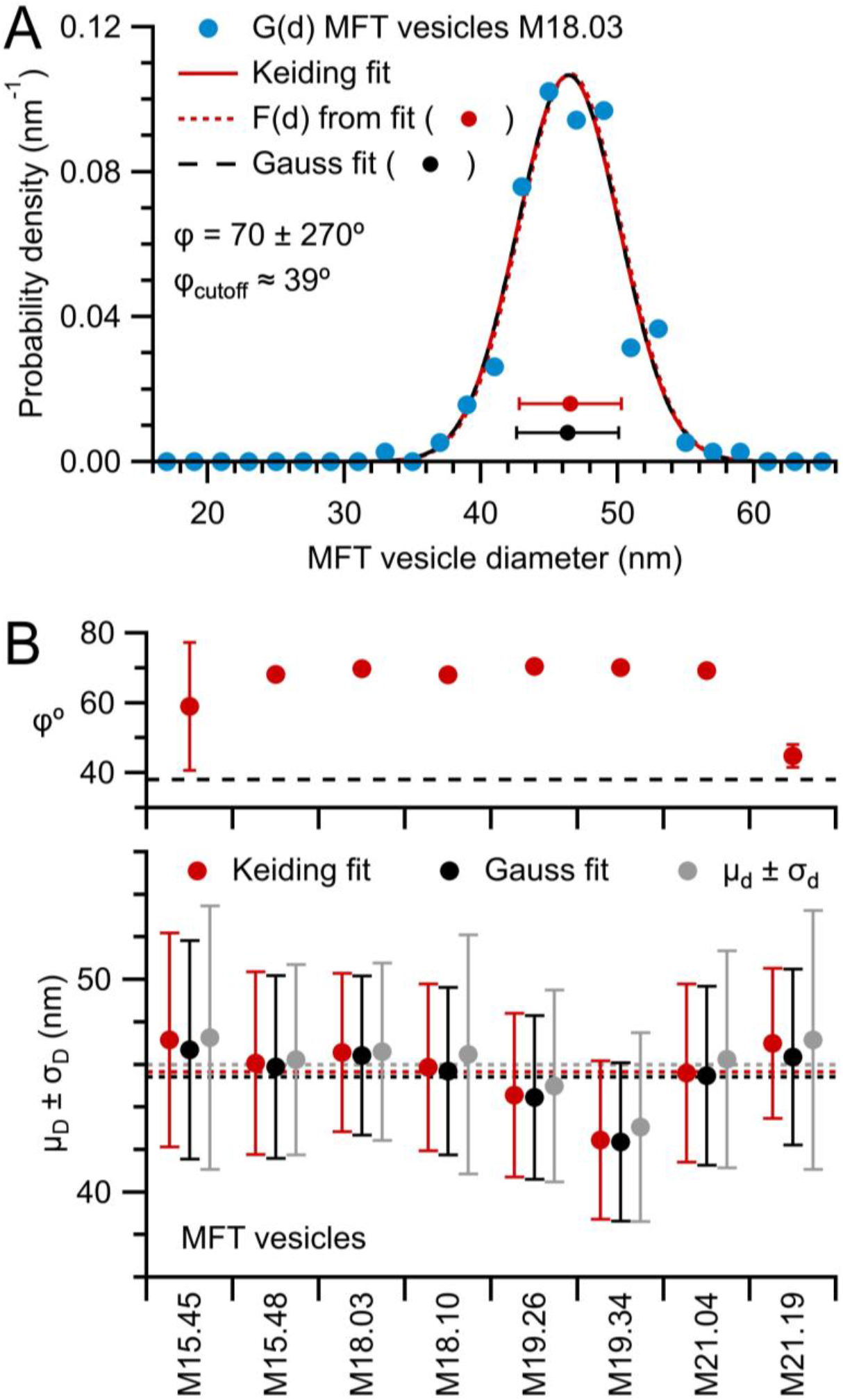
Estimates of F(d) and ϕ from G(d) of MFT vesicles. (A) Curve fit of Equation 1 (red solid line) to the G(d) of mouse MFT vesicles in Figure 2C2 (blue circles) resulting in estimates for F(d) (red dotted line and circle), ϕ and ϕ_cutoff_ (Equation 9). A Gaussian fit to the same G(d) (black dashed line) overlaps the Keiding-model fit and μ_D_ ± σ_D_ of both fits overlap (red and black circles) indicating G(d) ≈ F(d). (B) Parameters μ_D_ ± σ_D_ (bottom) and ϕ (top) from Keiding-model curve fits to G(d) of MFT vesicles for 4 mice, 2 MFTs per mouse (red circles; Figure S15), compared to μ_D_ ± σ_D_ from Gaussian fits (black circles) and μ_d_ ± σ_d_ computed from 2D diameters. Overlapping distributions again indicate G(d) ≈ F(d). Dotted lines (bottom) denote averages. Black dashed line (top) denotes average estimated ϕ_cutoff_ (∼38°); all estimated ϕ are considered inaccurate. Error bars of ϕ denote fit errors, 6 of which are not shown since they are off scale.

### 2.10 Estimating the 3D density of particles from their 2D projection

Having an accurate estimate of the number of particles per unit volume is essential to many fields of science. Historically, estimates of 3D particle density (λ_3D_) were obtained by computing the number of particles per unit area (λ_2D_; Figure S1) observed in a projection divided by the section thickness (T). However, particle caps on the top and bottom of the section inflate the particle count with respect to T, i.e. they create an overprojection^17,40–43^. To correct for overprojection, one can use ϕ to estimate λ_3D_ via Equation 3 (Appendix A).

As a first test of the validity of Equation 3, we used the Monte Carlo simulations in Figure 4C to investigate the relation between the measured λ_2D_ and true λ_3D_ by computing ζ = λ_2D_ / λ_3D_. Comparison of this measured ‘true’ ζ to the expected ζ computed via Equation 2, using true T, μ_D_ and ϕ, showed a close agreement (Figure 10A1). Hence, our simulations of the Keiding model were well described by Equation 3. Next, we tested the ability of the Keiding model to accurately estimate λ_3D_ via Equation 3 by computing Δλ_3D_ for the same simulations using parameters μ_D_ and ϕ estimated via Keiding-model curve fits to G(d) (Figure 4C). Results showed accurate estimates of λ_3D_, except when true ϕ > 45°, in which case the estimation errors of μ_D_ and ϕ translated into estimation errors of λ_3D_ (Figure 10A2). Here, the finding that ϕ_cutoff_ for estimated λ_3D_ is less than that for estimated μ_D_, σ_D_ and ϕ (∼55°) is consistent with an increase in variability from using estimated μ_D_ and ϕ to compute ζ, and therefore consistent with our ϕ-accuracy test, where estimated ϕ_cutoff_ ≈ 43° (Equation 9). Hence, these results demonstrate the ability of Equation 3 to accurately estimate λ_3D_ using Keiding-model estimates of μ_D_ and ϕ as long as estimated ϕ < estimated ϕ_cutoff_. Comparing results for planar and thick sections showed qualitatively similar results, except errors were smaller for thick sections due to smaller errors in μ_D_ (but see next paragraph for caveats of using thick sections). Repeating the error analysis for G(d) computed from ∼2000 diameters gave qualitatively similar results compared to those for ∼500 diameters, but λ_2D_ and λ_3D_ had smaller biases and confidence intervals, where the confidence intervals followed a 1/√n elation for ϕ < 45° (Figure S17) similar to that of estimated μ_D_ and ϕ (Figure S5).

**Figure 10.**
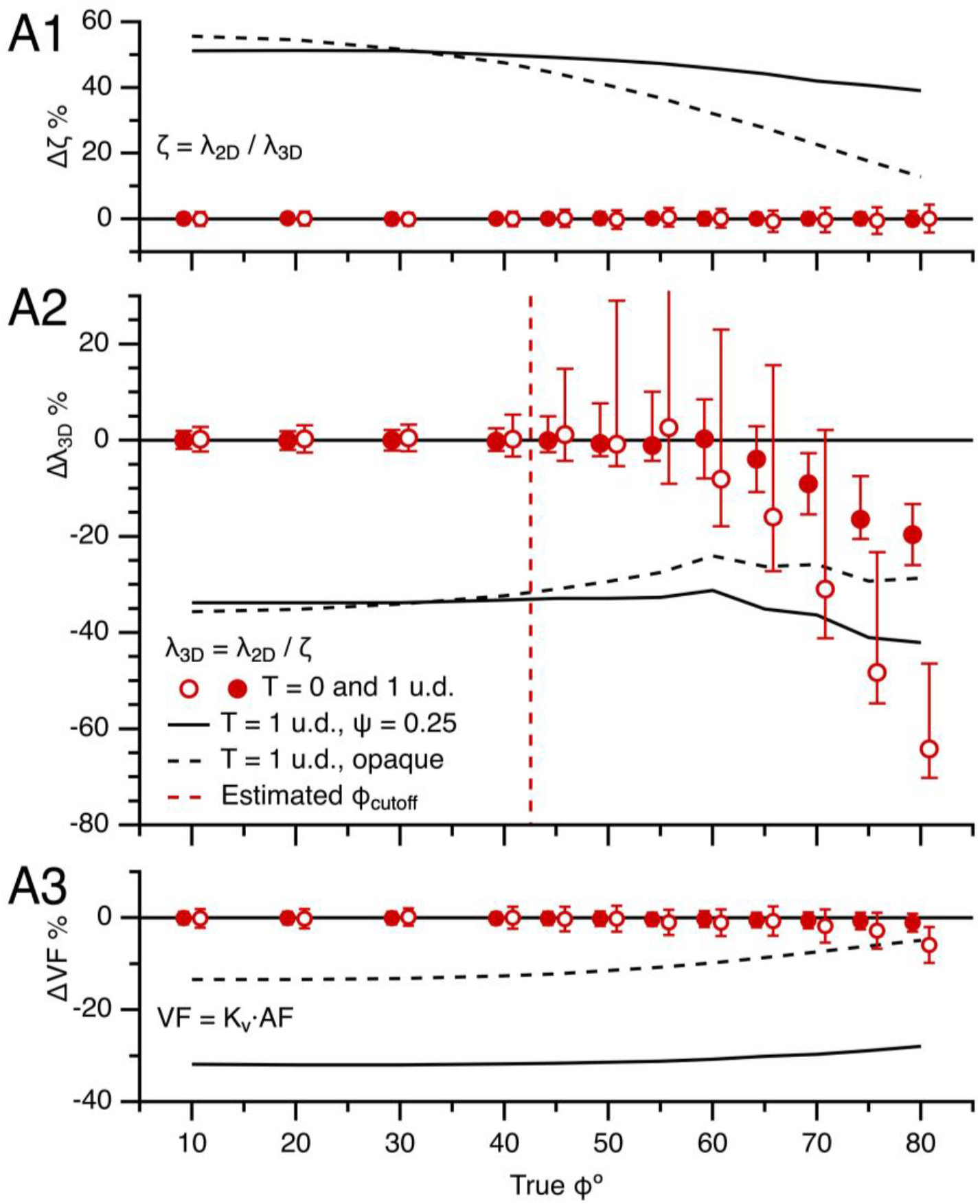
The Keiding model accurately estimates λ_3D_ from λ_2D_ for true ϕ < estimated ϕ_cutoff_ (Part I. simulations). (A1) Average error in section z-depth over which particle center points are sampled (Δζ) vs. true ϕ (top) for simulations in Figure 4C for T = 0 and 1 u.d. (red open and closed circles) where estimated ζ was computed via Equation 2 using true μ_D_, ϕ and T and ‘true’ ζ = λ_2D_/λ_3D_ using measured λ_2D_ and true λ_3D_. Data x-scales were shifted ±0.8° to avoid overlap. (A2) Average error in 3D particle density (Δλ_3D_) vs. true ϕ (middle) for the same simulations, computed via estimated and true λ_3D_, where estimated λ_3D_ was computed via Equation 3 (λ_3D_ = λ_2D_/ζ) using measured λ_2D_ and estimated μ_D_ and ϕ from Keiding-model fits to the simulated G(d) (Figure 4C). Red dashed line denotes estimated ϕ_cutoff_ (∼43°; Equation 9). (A3) Average error in volume fraction (ΔVF) vs. true ϕ (bottom) for the same simulations, computed via estimated and true VF, where estimated VF = K_v_·AF (Equations B.1, B.2c). Results are also shown for transparent particles (black solid line; ψ = 0.25) and opaque particles (black dashed line) in thick sections (T = 1 u.d.; Figure 5), showing estimated ζ was larger than true ζ when projection overlaps hindered particle counting, creating a large underestimation of λ_3D_ and VF. However, for opaque particles, the bias was larger for λ_3D_ than VF since an overlap in two projections reduced the particle count by one, but only partially reduced the AF (overlapping projections coalesce into one).

For thick sections, a major caveat of estimating λ_3D_ from λ_2D_ via Equation 3 is that the calculation assumes one has a good estimate of ζ. As shown in Figure 5B, however, overlapping projections of semi-transparent and opaque particles may preclude counting particles at the bottom of the section, thereby reducing ζ. To demonstrate this, we computed Δζ for simulations similar to those in Figure 5 and found large positive biases for both semi-transparent and opaque particles, i.e. estimated ζ was larger than true ζ (Figure 10A1). As expected, the overestimation of ζ translated into large negative biases in estimates of λ_3D_ (Figure 10A2). Given this caveat, therefore, the best approach to estimate λ_3D_ from λ_2D_ for particles with a high density is to use planar sections, in which case there will be little to no interference in counting particles from overlapping particle projections.

As a second independent method for computing the 3D density, we used the particle area fraction (AF) to VF relation of Weibel and Paumgartner^60^ (VF = K_v_·AF) to compute the VF of our simulations, where AF is the sum of all projection areas divided by the total projection area (Area_xy_) and K_v_ is a proportional scale factor that is a function of T and μ_D_, but modified to be a function of ϕ rather than h_min_ (Appendix B). Results showed excellent agreement between the estimated and true VF of our simulations (Figure 10A3). However, for the simulations of overlapping projections of semi-transparent and opaque particles in thick sections, there was a significant underestimation of the VF, as expected.

### 2.11 Validation of the Keiding model using 3D reconstructions (vesicle density)

To test the accuracy of applying Equation 3 to real data, we used one our ET z-stacks of MFT vesicles (ET11) to measure λ_2D_ and λ_3D_ within a subregion of the z-stack where vesicles were clustered (Figure 11A1 and A2). This analysis gave λ_2D_ = 369.1 μm^−2^ and λ_3D_ = 11,348 μm^−3^ (VF = 0.47) where λ_3D_ was computed as a 3D measurement, i.e. count per volume. Using Equation 3, we then computed λ_3D_ = λ_2D_/ζ = 11,503 μm^−3^ (VF = 0.48; Equation 4) where ζ was computed using the measured μ_D_ and ϕ via Equation 2 (ζ = 32.1 nm; μ_D_ = 42.7 nm, ϕ = μ_ϕ_ = 41.4°). Hence, λ_3D_ computed via Equation 3 was nearly the same as that computed via a direct 3D measurement (Δλ_3D_ = +1%). Repeating the same computation using ζ estimated via Keiding-model curve-fit parameters μ_D_ and ϕ (ζ = 33.6 nm; μ_D_ = 42.9 nm, ϕ = 38.4°), where estimated ϕ < estimated ϕ_cutoff_ (∼51°), gave λ_3D_ = 10,991 μm^−3^ (VF = 0.46; Equation 4), which was also nearly the same to λ_3D_ computed via a 3D measurement (Δλ_3D_ = -3%). In contrast, when we used d_min_ of the z-stack analysis (19 nm) to estimate ζ (using measured μ_D_) rather than ϕ (where d_ϕ_ = 28 nm), we computed λ_3D_ = 9688 μm^−3^. Hence, in this example, using d_min_ to estimate λ_3D_ caused significant error (Δλ_3D_ = -15%). Next, we investigated the range of expected error for the z-stack density analysis, as well as that from assuming a fixed ϕ (Figure S10), by computing λ_2D_ and λ_3D_ for Monte Carlo simulations that mimicked the 3D analysis. Results gave ranges of Δλ_3D_ that were consistent with the above measured Δλ_3D_ (Figure S18) and showed the assumption of a fixed ϕ introduces only small errors for estimating λ_3D_ from λ_2D_ via Equation 3 for ϕ < 50° (Figure S19). Again, using d_min_ to estimate λ_3D_ caused significant error (Δλ_3D_ = -19.2 ± 1.9%; n = 100 simulation repetitions). Finally, using the VF = K_v_·AF relation (Equations B.1, B.2c) we computed a VF via the 2D analysis (0.46) that matched that computed via the 3D analysis (0.47). Repeating the same 2D versus 3D density analysis for our other ET z-stack (ET10) produced similar results (Figure 11B; Table 3). However, unlike ET11, the density analysis of ET10 had to be confined within a subregion of the z-stack due to a nonhomogeneous distribution of vesicles in the axial axis. Overall, our 3D analysis of MFT vesicles shows that Equation 3 accurately estimated λ_3D_ from λ_2D_ with only small error when true ϕ < ϕ_cutoff_, confirming our previous results from simulations (Figure 10).

**Figure 11.**
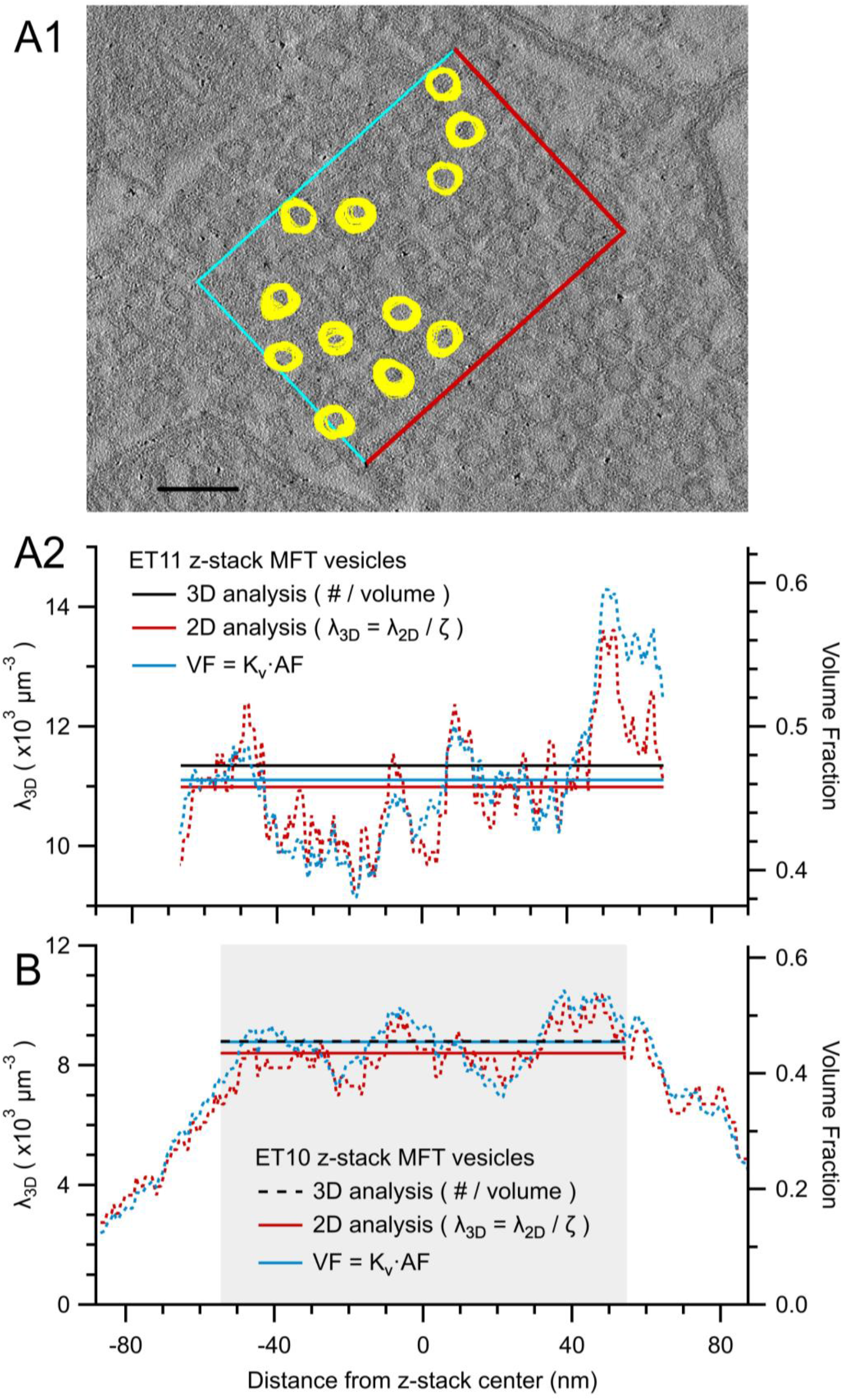
The Keiding model accurately estimates λ_3D_ from λ_2D_ for true ϕ < estimated ϕ_cutoff_ (Part III. vesicles). (A1) One of 261 serial ET images (ET11) of a cerebellar tissue section, where the z-distance between each image (R_z_) is estimated to be 0.51 nm (Figure S7B). A ROI (0.45 × 0.32 μm; Area_xy_ = 0.144 μm^2^) was placed within a large cluster of MFT vesicles and those vesicles that obeyed the inclusive/exclusive borders (blue/red; n = 271) were tracked and outlined through multiple z-planes. Because the analysis includes vesicle caps on the top/bottom of the reconstruction, the vesicle sampling space of the volume of interest (VOI) extends above/below the section such that ζ = 165 nm (Equation 2; 3D measures: T = 133 nm, μ_D_ = 42.7 nm, ϕ = 41°), giving VOI = 0.024 μm^3^. Here, outlines for 12 representative vesicles are overlaid with outlines from images above and below. Scale bar 100 nm. (A2) Vesicle λ_3D_ (left axis) and VF (right axis) computed within the ROI in (A1). For the 3D analysis, λ_3D_ = 11,348 μm^−3^, the number of vesicles within the VOI volume (black solid line). For the first 2D analysis, λ_3D_ = λ_2D_/ζ (Equation 3) computed for each z-plane (red dotted line) and sum of all z-planes (red solid line; 10,991 μm^−3^), where λ_2D_ is the number of outlines per ROI area and ζ = 34 nm computed via Keiding-model estimates (T = 0 nm, μ_D_ = 42.9 nm, ϕ = 38°; Figure 6D). For the second 2D analysis, VF = K_v_·AF (Equations B.1, B.2c) computed for each z-plane (blue dotted line) and sum of all z-planes (blue solid line; 0.46), where K_v_ = 1.07 and AF is the sum of all vesicle outline areas per ROI area. Both true ϕ (41°) and estimated ϕ (38°) are less than estimated ϕ_cutoff_ (51°; Equation 9). Left and right axes are equivalent scales for μ_D_ ± σ_D_ = 42.7 ± 3.4 nm, the measured F(d). (B) Same as (A2) for ET10. Grey shading denotes the axial subregion where 3D analysis and averages were computed. Left and right axes are equivalent scales for μ_D_ ± σ_D_ = 45.6 ± 4.2 nm. See Materials and Methods.

### 2.12 Estimating the 3D particle density via the disector method (a comparison)

The disector method^18,52^ is a popular method for estimating λ_3D_ of particles with arbitrary geometry using two adjacent sections from a z-stack, referred to as the ‘reference’ and ‘lookup’ section. This method counts the number of particles that appear within the reference section but do not appear in the lookup section, i.e. the method counts a particle only if its leading edge appears within the reference section. To compute λ_3D_, one determines λ_2D_ from the leading-edge count and divides λ_2D_ by the distance between the reference and lookup sections (i.e. the section thickness for adjacent pairs). Although theoretically, lost caps should not introduce error into this counting method^52^, underestimation errors due to lost caps have been reported^61^. Given the popularity of the disector method, as well as the potential source of error due to lost caps, we decided to use the disector method to estimate λ_3D_ of our simulated projections where ϕ = 10–80° and compare these results to our analysis in Figure 10, where λ_3D_ was estimated via Equation 3, i.e. the ϕ-correction method. Using T = 0.3 u.d. as recommended^18^, we found no bias due to lost caps for all ϕ tested (Figure 12, black squares; ∼500 particles per projection). This result is consistent with historical thinking that lost caps do not create a bias in counting: while the effect of lost caps on any given reference section is to reduce the particle count, the effect of lost caps on the lookup section is to increase the particle count for that reference section, the two effects thereby cancelling^52^. In essence, lost caps simply shift the reference section to which particles are counted and therefore have no effect on the final particle count. However, these simulations did not take into account a potentially significant bias pointed out by Hedreen^27^: fewer caps are likely to be identified in the reference section compared to the lookup section since the former is done blind and the latter is not (the researcher searches for caps in the lookup section only after identifying a particle in the reference section). In fact, our analysis of ET z-stacks of MFT vesicles revealed a 17–22° bias in ϕ between a blind and nonblind vesicle detection (Figures 7). To investigate the effects of such a bias, we added a bias to our disector simulations by increasing ϕ of the reference section with respect to the lookup section (ϕ_bias_ = 5–20°) thereby reducing cap identification in the reference section (i.e. increasing the number of lost caps) with respect to the lookup section. Interestingly, results of these simulations showed large underestimation errors even for small ϕ_bias_ (Figure 12; - 5% < Δλ_3D_ < -20% for ϕ_bias_ = 10°). Hence, the blind-versus-nonblind bias in particle detection has the potential to create a significant underestimation of λ_3D_. Repeating the disector bias analysis using our experimental ET z-stack data gave similar results, showing a large error (Δλ_3D_ = -41 ± 5%) for the average measured ϕ_bias_ = 20°.

**Figure 12.**
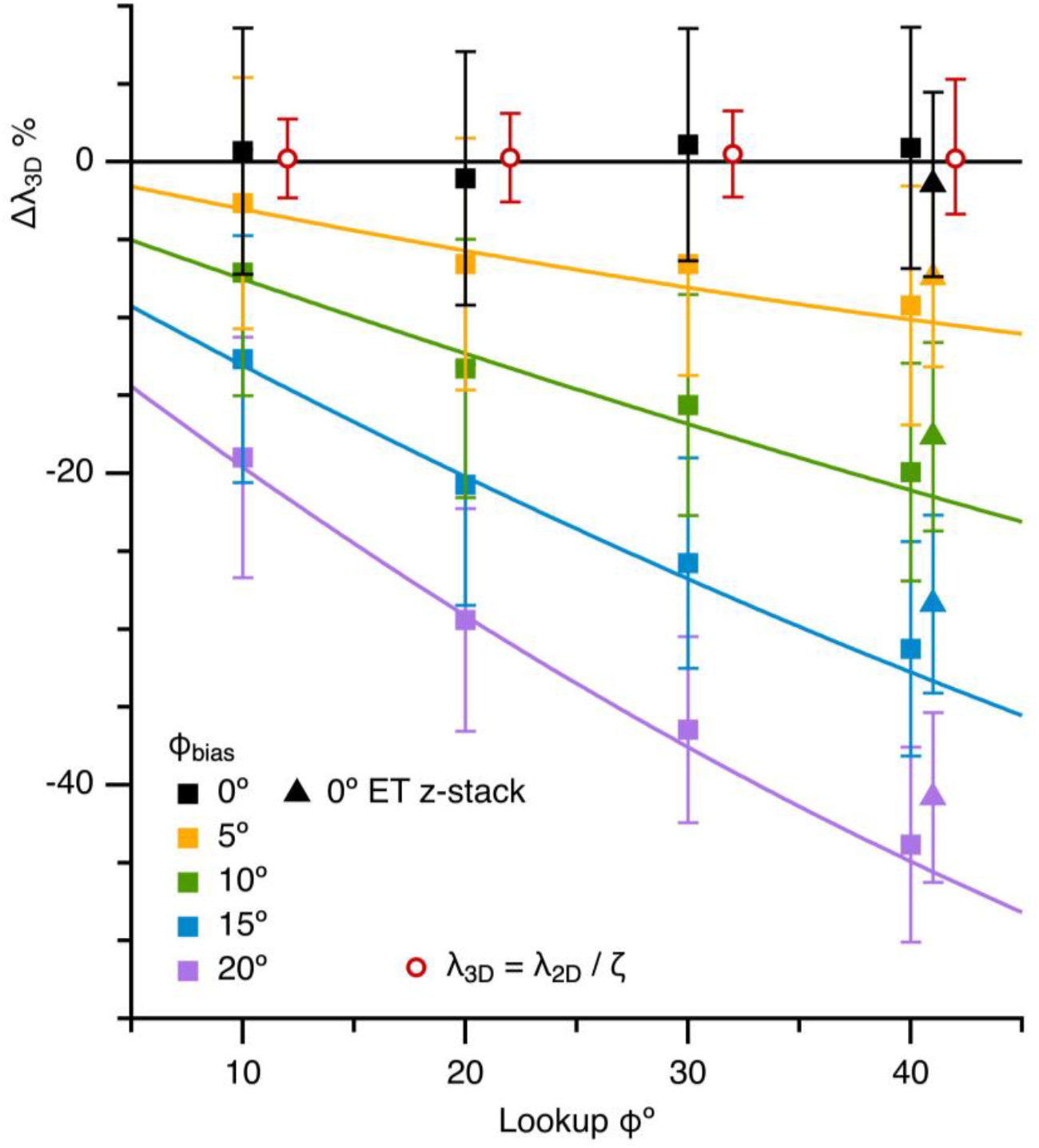
Estimation errors of the 3D density computed via the disector method for different degrees of bias due to blind-versus-nonblind cap detection. Average Δλ_3D_ computed from Monte Carlo simulations of the disector method, where a particle-detection bias was simulated by increasing ϕ of the reference section (a blind cap detection) with respect to the lookup section (a nonblind cap detection) for ϕ_bias_ = 0–20° (colored squares; ϕ_ref_ = ϕ_lookup_ + ϕ_bias_; ∼500 particles per reference section). Thickness of the reference and lookup sections was 0.3 u.d. An equivalent disector analysis using the ET11 z-stack data gave similar results (triangles; lookup ϕ = 41°). Average Δλ_3D_ for the simulations in Figure 10A2 are shown for comparison (red open circles; T = 0 u.d.; λ_3D_ = λ_2D_/ζ; x-scale is true ϕ and is shifted 2° to avoid overlap).

Besides the potential error due to a blind-versus-nonblind bias in particle detection, our simulations show that the disector method is ∼2 to 3-fold less accurate at estimating λ_3D_ (i.e. Δλ_3D_ has a larger ±σ) compared to the ϕ-correction method (Figure S17; ±σ panel) due to a lower particle count per section. Similar results were found using the disector method to compute λ_3D_ of our ET z-stack of MFT vesicles, where Δλ_3D_ = ±6% for the disector method (±σ; ϕ_bias_ = 0°) compared to Δλ_3D_ = ±2% for λ_3D_ computed via Equation 3 (Figure S18). Hence, the inherent smaller count per region of interest (ROI) of the disector method leads to a larger uncertainty in estimates of λ_3D_.

### 2.13 Estimation of the 3D density of granule cell somata and nuclei (confocal vs. TEM)

Having verified Equation 3 using simulations and experimental data, we applied the equation to the estimation of λ_3D_ from λ_2D_ for our sample of GC somata. Using the same confocal images used to compute the 9 G(d) of the GC somata of 3 rats, we computed λ_2D_ = 17.7–22.6 × 10^3^ mm^−2^ (Figure S1A; 8792–29,845 μm^2^ ROI area per image, 196–528 somata per ROI, 3 images per rat). Using estimated μ_D_ and ϕ for each G(d), we then computed ζ = 5.5–7.6 μm via Equation 2 (estimated T = 1.4–2.4 μm, scaled for z-shrinkage; Table 1). Finally, we computed λ_3D_ = λ_2D_/ζ = 2.7–4.1 × 10^6^ mm^−3^ with mean λ_3D_ = 3.2 ± 0.2 × 10^6^ mm^−3^ and VF = 0.32 ± 0.01 (±SEM; Table 2; Equation 4). To verify these results using the relation VF = K_v_·AF, we computed K_v_ = 0.58–0.77, AF = 0.39–0.50 and VF = K_v_·AF = 0.27–0.37, with mean VF = 0.32 ± 0.02 and λ_3D_ = 3.1 ± 0.1 × 10^6^ mm^−3^ (±SEM) which is not significantly different to λ_3D_ computed via Equation 3 (p = 0.4, paired *t*-test).

Next, we estimated λ_2D_ of GC nuclei from the TEM images used to compute G(d) (λ_2D_ = 17.1–34.2 × 10^3^ mm^−2^; Figure S1B; 2046–5747 μm^2^ ROI area per image, 35–158 nuclei per ROI, 6–7 images per mouse). Using estimated μ_D_ and ϕ for each G(d), we then computed ζ = 4.2–4.8 μm via Equation 2 (T = 0.06 μm) and λ_3D_ = λ_2D_/ζ = 3.5–8.1 × 10^6^ mm^−3^, with mean λ_3D_ = 5.9 ± 0.4 × 10^6^ mm^−3^ and VF = 0.34 ± 0.01 (±SEM; Table 2; Equation 4). Simulations indicate λ_3D_ for this dataset should have negligible bias with a small confidence interval (Figure 10; Δλ_3D_ = +0.3 ± 2.8 % for T = 0 u.d. and true ϕ = 20°). To verify these results using the relation VF = K_v_·AF, we computed K_v_ = 0.98–0.99, AF = 0.27–0.39 and VF = K_v_·AF = 0.27–0.39, with mean VF = 0.33 ± 0.01 and λ_3D_ = 5.7 ± 0.5 × 10^6^ mm^−3^ (±SEM; n = 4) which is not significantly different to λ_3D_ computed via Equation 3 (p = 0.1, paired *t*-test). Finally, to scale the VF estimates from nuclei to somata, we estimated the VF for GC somata of mice via Equation 4 using estimated F(d) for GC somata of mice (Figure 8C; N→S) and assuming λ_3D_ was the same for the nuclei and somata. Results gave VF = 0.51–0.57 with mean 0.54 ± 0.01, suggesting GC somata occupied half of the GC layers we analysed.

Unlike our estimates for F(d) of GCs in rats and mice, which show no difference, comparison of the above estimates for λ_3D_ in rats and mice show a 2-fold difference: estimated λ_3D_ of the rat dataset (3.2 × 10^6^ mm^−3^) is significantly smaller than estimated λ_3D_ of the mouse dataset (5.9 × 10^6^ mm^−3^; p = 0.004). While this difference could reflect a true difference between species, a previous comparative study of the cerebellum, which used the same methodology across species, reported similar densities within the GC layer of rats and mice^62^. Hence, it is more likely that the 2-fold difference in estimated λ_3D_ is due to one or more differences in methodology. Although there are perhaps too many differences in methodology to make a decisive conclusion, notwithstanding the confounding problem of the ‘reference trap’^63^, it is still instructive to consider them here. The first difference in methodology is section thickness: thick versus planar. Because the rat dataset was derived from thick sections, overlapping opaque projections could have created an underestimate of λ_3D_ compared to the mouse dataset (Figure 10). The second difference in methodology is imaging technology: confocal versus TEM. Because of the lower contrast and resolution of the confocal images, it was considerably harder to identify and delineate GC somata profiles compared to the GC nuclei profiles. The poor delineation of GCs would create an undercount. The third difference in methodology is tissue preparation: chemical versus cryo fixation. A change in the extracellular volume due to chemical fixation of the rat tissue preparation could have created a lower λ_3D_. However, this explanation is unlikely since chemical fixation tends to reduce the extracellular volume^59^ which would result in a higher rather than lower λ_3D_. The final difference in methodology is the sampling space: ROIs were larger in the rat dataset compared to the mouse dataset. Because GCs are not distributed uniformly, but rather form high-density clumps interspersed by MFTs and blood vessels, there could be a bias towards a larger λ_3D_ in the mouse dataset due to smaller ROIs. However, our analysis of the two datasets using different sized ROIs (not shown) indicates the difference in ROI size cannot account for the 2-fold difference in estimated λ_3D_. Hence, we suspect our estimate of λ_3D_ in the rat dataset is underestimated, and this is most likely due to overlapping projections in thick sections and poor delineation of GC profiles in the confocal images. Because of the tight packing of GC somata, estimates of λ_3D_ via Equation 3 are best achieved by using planar sections, a higher contrast preparation and superior microscope resolution. We therefore believe our most accurate estimate of the GC λ_3D_ is that of our mouse dataset. Although our estimated λ_3D_ of GC somata in rats is likely to be underestimated, it is still 1.7-fold larger than our previous estimate from the same confocal images^6^. Because the latter estimate of λ_3D_ was computed via the disector method, we suspect it is underestimated due to the blind-versus-nonblind bias in particle detection discussed in the previous section. Hence, this result suggests the local MFT-to-GC expansion ratio is larger than previously estimated^6^.

### 2.14 Estimation of the 3D density of clustered vesicles in mossy fiber terminals (TEM vs. ET)

Although ϕ was indeterminable for our analysis of MFT vesicles in TEM images, it was still possible to estimate λ_3D_ over a range of ϕ, i.e. ϕ > ϕ_cutoff_ or ϕ = ϕ_cutoff_–90°. First, we computed λ_2D_ of MFT vesicles (Figure S1C) using the same TEM images used to compute their G(d) (Figure S15; n = 8), giving λ_2D_ = 305–429 μm^−2^. Next, we computed ζ = 99–60 nm via Equation 2 (T = 60 nm) for ϕ = 38–90° (Equation 9). Finally, we computed λ_3D_ = 3100–7200 μm^−3^ via Equation 3. Not surprisingly, this range of λ_3D_ is considerably smaller than that for our 3D ET analysis of MFT vesicles (λ_3D_ = 9,000–11,000 μm^−3^). Again, similar to our analysis of GCs in confocal images, we suspect λ_3D_ is underestimated due to overlapping projections in thick sections (Figure 10). To estimate the effective section depth of those vesicles counted, we assumed λ_3D_ of the TEM dataset was the same as that of our ET dataset and computed ζ = λ_3D_/λ_2D_ = 30–43 nm. This range of ζ is ∼2.0-fold smaller than that estimated via Equation 2, indicating vesicles at the bottom of the tissue sections were not counted (Figure 5B), creating an underestimation of λ_3D_. Hence, this analysis lends support to our conclusion that the optimal method for estimating λ_3D_ from λ_2D_ for particles with a high density is to use planar sections, in which case there will be no interference of counting from overlapping projections. Since the thinnest possible tissue section created via an ultramicrotome is currently on the order of a vesicle, the best option for computing vesicle density is via ET reconstructions (Figure 11).

## 3 Discussion

Stereological methods for estimating the 3D size distribution (F(d)) and density (λ_3D_) of a collection of particles from their 2D projection are essential tools in many fields of science. These methods, however, inevitably contain sources of error, one being the unresolved or nonexistent profiles known as lost caps. Surprisingly, the simple solution for lost caps developed by Keiding et al.^49^, which defines lost caps of spherical particles with respect to a single (i.e. fixed) cap-angle limit (ϕ), has not been widely adopted and has never been validated. Here, we provide the first experimental validation of the Keiding model by quantifying ϕ of unresolved vesicle caps within 3D reconstructions. While this analysis reveals a Gaussian distribution for ϕ rather than a single value, curve fits of the Keiding model to the 2D diameter distribution (G(d)) nevertheless accurately estimate the mean ϕ, as well as F(d). Parameter space evaluation with Monte Carlo simulations revealed that the estimates are most accurate when ϕ falls below a specific value (ϕ_cutoff_). Hence, our experimental and theoretical analyses reveal that, if one only wishes to estimate F(d) from a 2D projection, then the Keiding model can be called to task, whether one is using planar or thick sections. On the other hand, if one wishes to estimate both F(d) and λ_3D_ from a 2D projection, then one will need an accurate estimate of ϕ (Figure 13). As we discuss below, obtaining an accurate estimate of ϕ for some preparations may require optimising experimental conditions.

**Figure 13.**
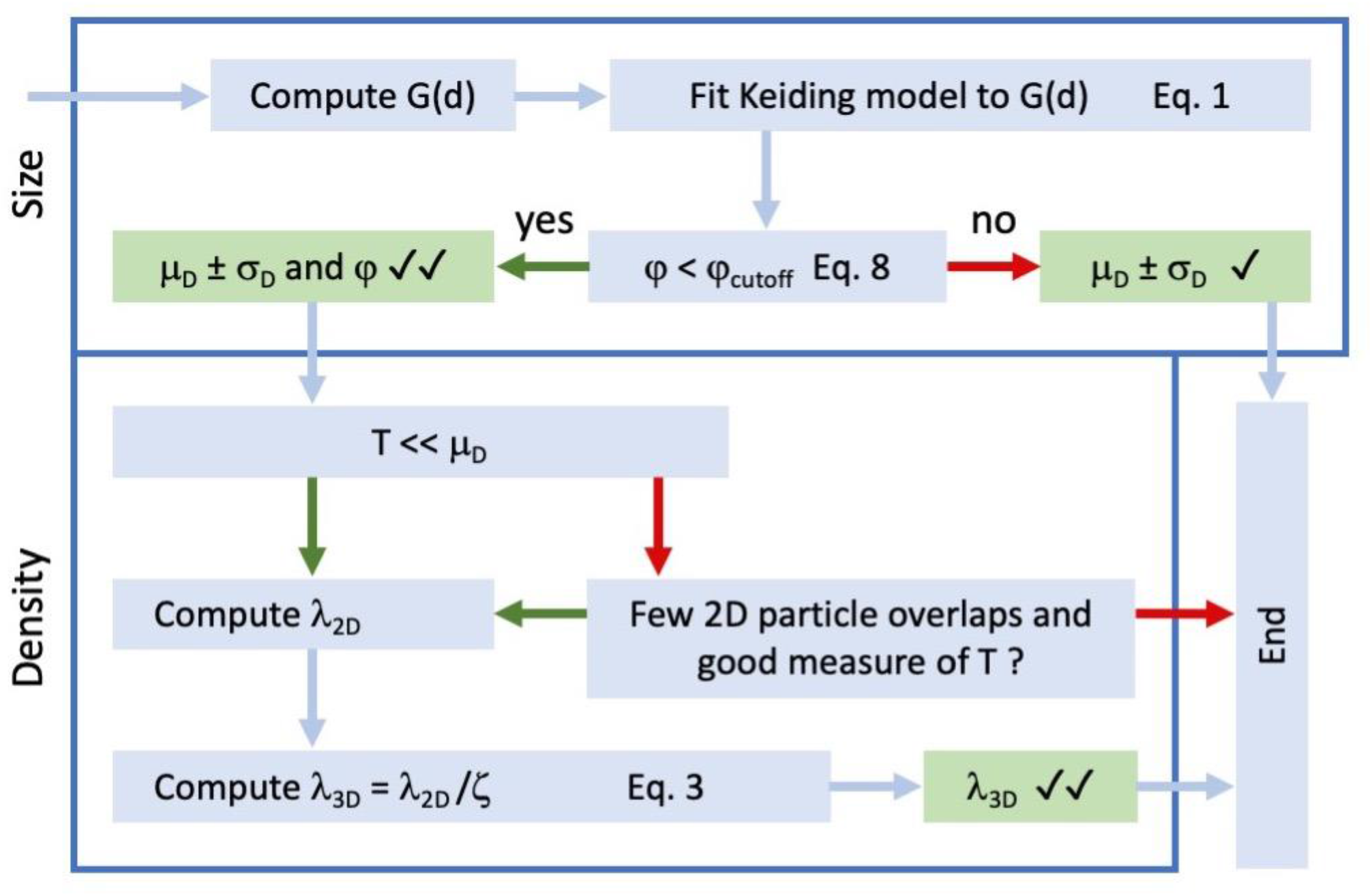
Methods workflow for estimating the 3D size and density of spherical particles from their 2D projection. Workflow diagram describing the sequence of steps for estimating F(d) (i.e. μ_D_ ± σ_D_) and ϕ from G(d), and λ_3D_ from λ_2D_, where G(d) and λ_2D_ are computed from a 2D projection of randomly-distributed particles in a section of thickness T. Two checks indicate a negligible bias and small confidence interval compared to one check. Final estimates of λ_3D_ can be converted to VF using estimated μ_D_ ± σ_D_ (Materials and Methods) and compared to that computed via the relation VF = K_v_·AF (Equations B.1, B.2c). Estimated ϕ is compared to estimated ϕ_cutoff_, computed via fit parameters μ_D_ and σ_D_ (Equation 9).

### 3.1 Basic assumptions of the Keiding model

There are five basic assumptions of the Keiding model^49^ that one must keep in mind when applying it to the estimation of particle size and density via Equations 1–3.

The first assumption is that the particles of interest are approximately spherical, i.e. they are convex with the average shape of a sphere in rotation, which includes elliptical^16^. The assumption of a spherical shape is usually valid for vesicles, vacuoles and nuclei^1,17,44,49,64^, but also for large structures such as follicles^16^ and glomeruli^65^.

The second assumption is that F(d) is well described by a probability density function (PDF), e.g. a Gaussian, chi or gamma distribution. The assumption that F(d) was a simple Gaussian distribution worked well for our analysis of GC somata and nuclei and MFT vesicles, and for the liver cell nuclei of Keiding et al.^49^ (Figure S3). Similarly, the assumption that F(d) was a chi distribution worked well for Wicksell’s corpuscles (Figure S4). If uncertain of the shape of F(d), one can compare fits to distributions obtained from thick sections (T > D) where it is likely that G(d) ≈ F(d) (Figures 2C2 and S3).

The third assumption is that the particles have a random distribution; or if the particles are clustered, then the clusters have a random distribution. It is not necessary that the spatial distribution is perfectly random (i.e. Poisson) but rather there is no order to the distribution. For example, a lattice structure (e.g. hexagonal packing) would be problematic since the particles in a 2D projection would have a discrete size distribution.

The fourth assumption is that ϕ is the same (fixed) for all particles. Because our 3D analysis of MFT vesicles revealed not a single value for ϕ, but a range of ϕ well described by a Gaussian distribution (CV_ϕ_ = 0.2), this assumption could be problematic. However, we found curve fits of the Keiding model to G(d) accurately estimated the mean of this distribution (Figures 6D and S9D). Moreover, by incorporating a Gaussian-ϕ model into our Monte Carlo simulations, we were able to measure the bias introduced by the fixed-ϕ assumption over a range of ϕ distributions (i.e. μ_ϕ_) and found the bias was relatively small for μ_ϕ_ < ϕ_cutoff_ (Figures S11 and S19). Because vesicles are at the lower limit of resolution, we suspect the spread of our measured ϕ distribution, i.e. CV_ϕ_ = 0.2, may be close to a worst-case scenario.

The fifth assumption pertains to thick sections: that they are perfectly transparent so that one observes 2D projections of particles at the bottom of the section as well as those of particles at the top of the section. For opaque particles with high density this assumption is clearly problematic^2,57^. For semi-transparent particles with high density this assumption is less problematic since overlapping projections do not necessarily preclude drawing their outline (Figure 2C1) or counting them (Figure S1C). Yet, one can imagine a section of enough thickness that overlapping projections preclude outlining or counting the particles at the bottom of the section. Using Monte Carlo simulations, we tested the effect of such a scenario by removing particles from simulated projections if they experienced a total sum of overlaps greater than a set limit. Interestingly, results of these simulations showed only a small effect on the shape of G(d) and therefore only small biases for estimated F(d) (Figure 5C). In contrast, our simulations of opaque particles showed a larger effect on the shape of G(d), i.e. a positive skew, since overlapping projections coalesced into larger ones; yet, the end result was only small biases for estimated F(d), mostly a positive bias in σ_D_ (Figure 5D). Estimates of λ_3D_ for the same simulations of semi-transparent and opaque particles, on the other hand, showed large underestimations due to a reduction in the effective ζ (Figure 10). Hence, violation of this fifth assumption is likely to create only small biases in estimated F(d), but large biases in estimated λ_3D_. Our analysis of GC somata in confocal images and MFT vesicles in TEM images, for example, are likely examples of this scenario.

As with other model-based stereological methods, the Keiding model has a number of core assumptions. Nevertheless, our analyses show that for approximately spherical particles these assumptions are either reasonable or can be circumvented with the correct experimental paradigm.

### 3.2 Experimental considerations for optimising the detection of small caps and minimising the cap-angle limit ϕ

Our analysis shows that estimates of F(d) and λ_3D_ are most accurate when true ϕ < ϕ_cutoff_ (Figures 4C and 10). To minimise ϕ, one needs to visually detect a wide spectrum of cap sizes within a 2D projection (i.e. sample as much of G(d) as possible). Here, we suggest experimental conditions/recommendations that could help optimise particle cap detection. First, sections should be thin (T ≈ 0.3 u.d.) to planar (T ≈ 0 u.d.) to avoid overlaps in the particle projections; the use of planar sections is particularly important for particles with a high density. Thin or planar sections can be achieved via ultrathin tissue sections (e.g. an ultramicrotome for TEM) or a high axial resolution of the microscope (ρ_z_; e.g. confocal imaging or ET). Second, the lateral resolution of the microscope (ρ_xy_) and image (R_xy_) should be high with respect to the size of the particle. Third, the efficiency of particle staining and contrast between the particles and their surrounding environment should be high. Fourth, surfaces of the tissue sections should be avoided when creating images, e.g. using guard zones in the axial axis, to avoid lost caps of the nonexistent type, i.e. caps that either fall off the surfaces of tissue sections or never existed, i.e. the microtome failed to transect the particles during sectioning^27,46–48^. Fifth, if the images to analyse exist within a z-stack, cap identification should be performed in a nonblind manner, i.e. by tracking particles through adjacent planes of the z-stack.

Given these considerations, our most suitable preparation for computing G(d) was that of the GC nuclei in high-resolution TEM images, where there were few lost caps and a small ϕ = 20°. For this dataset, the tissue sections were planar (T ≈ 0.01 u.d.) and the lateral resolution of the microscope was high with respect to the nuclei (ρ_xy_ = 1 × 10^−4^ u.d.; Table 1). Moreover, the GC nuclei were easy to identify and delineate due to their dark spotted appearance. In contrast, our analysis of GC somata in confocal images gave a larger number of lost caps and larger ϕ (37°). For this dataset, the optical sections were not planar (T ≈ 0.3 u.d), the lateral resolution of the microscope was comparatively low (ρ_xy_ = 0.06 u.d.) and, due to the opaque immunolabeling and dense packing of the somata, the task of delineating between adjacent somata was more difficult compared to that of the nuclei in TEM images.

Our least successful analysis for computing a complete G(d) was that of MFT vesicles. For example, our blind analysis of MFT vesicles in ET images resulted in a large number of lost caps and a large ϕ = 56–63° (Figures 7). This result is surprising given the virtual ET sections were thin in comparison to the vesicles (estimated ρ_z_ ≈ 0.1 u.d.) and the relative lateral resolution (ρ_x_ ≈ 0.09 u.d.) was comparable to that of the confocal images of GC somata. Even when the analysis was repeated in a nonblind manner, i.e. by tracking the vesicles through multiple planes of the ET z-stack, and dramatically reducing the number of lost caps, the resulting ϕ (∼40°) was only on par with that of the GC somata dataset. These results suggest the small size of synaptic vesicles places them at the lower limits of cap detection with respect to microscope resolution and image contrast. Similar to our blind analysis of MFT vesicles in ET images, our analysis of MFT vesicles in high-resolution TEM images resulted in a large number of lost caps and an estimated ϕ > ϕ_cutoff_. For this dataset, the tissue sections were thick in comparison to the vesicles (T ≈ 1.3 u.d.) but the lateral resolution was comparatively high (ρ_xy_ = 0.01 u.d.). The large estimated ϕ for this dataset is consistent with a blind cap detection and may also reflect the difficulty of identifying caps in thick sections with a high vesicle density. The absence of vesicle caps observed in thick sections has previously been noted^54,56^. In general, the results from our ET and TEM image analysis highlight the difficulty in computing a complete G(d) of MFT vesicles due to their small size.

### 3.3 Section thickness, axial distortions and the advantages of using planar sections

To convert measures of λ_2D_ to estimates of λ_3D_, both 2D model-based and 3D design-based stereological methods divide λ_2D_ by an estimate of the particle sampling space along the axial axis. For 2D methods, the axial sampling space is ζ in Equation 3, which is a function of μ_D_ and section thickness (T). For 3D disector methods, the axial sampling space is the distance between the reference and lookup sections (H). For both methods, sections are either physical tissue sections, in which case T and H are measures of the thickness of the tissue, or optical, in which case T is a measure of the axial resolution of the microscope (ρ_z_) and H is the distance between optical sections. Hence, to obtain an accurate estimate of λ_3D_, one must consider obtaining an accurate measure of T or H^2,25,26,45,66^. However, distortions of particle density along the axial axis of tissue sections must also be considered^2,28,30,48,67,68^. These include uniform shrinkage and differential deformations along the axial axis, and lost caps at the surfaces of the sections (i.e. nonexistent caps).

The challenges of estimating T and avoiding axial distortions of particle density also pertain to serial 3D reconstructions. Hence, the reason 3D reconstructions should not automatically be assumed to be the ‘gold standard’. For the 3D reconstructions used in this study, we avoided lost caps of the nonexistent type by avoiding the section surfaces. Moreover, the spherical shape of the vesicles allowed us to estimate the axial tissue shrinkage (Figure S7) which for EM can be considerable depending on the amount of electron beam exposure^69^. Finally, plots of vesicle density as a function of z-depth allowed us to verify that λ_3D_ was computed within a homogeneous vesicle distribution (Figure 11A2 and B). Hence, the effort to estimate λ_3D_ from a 3D reconstruction in this study was nontrivial.

With these axial-axes difficulties in mind, it becomes clear that the 2D methods have a distinct advantage over the 3D methods in that they can effectively remove T from the estimation of λ_3D_ by using planar sections. Under this condition, any bias due to an inaccurate estimate of T or uniform tissue shrinkage along the axial axis would be small. Hence, the 2D ϕ-correction method for estimating λ_3D_, when applied to data derived from planar sections, is potentially the most accurate method of estimating λ_3D_. However, planar sections come with their own challenges. First, planar sections can be technically challenging and costly to achieve, perhaps requiring TEM or ET. Second, due to the use of high-resolution microscopy, planar sections are likely to have a significantly smaller field of view, potentially creating bias if particles have a nonhomogeneous distribution. A smaller field of view, however, is not strictly prohibitive since it can be counteracted by analysing more images that have been acquired using a design-based random sampling strategy ^70,71^.

Our recommendation for the use of planar sections with 2D methods is counter to previous recommendations of using thick sections^25,26,47^. The reasoning put forth for using thick sections is that any bias introduced by lost caps (which affects the magnitude of μ_D_ in Equation 3) will be relatively small in comparison to T. However, this approach will only be valid if particles have a low density, one can reliably count particles at the bottom of the sections, one has a good measure of T and one can correct for any distortions of particle density along the axial axis, as discussed above. Moreover, our analysis indicates that when sections are thick, most caps are likely to be lost, in which case ϕ will be close to 90° and indeterminable (Figure 9 and Table 4). In the case of an indeterminable ϕ, one can use a range of ϕ (e.g. ϕ_cutoff_–90°) to estimate λ_3D_. This would be an improvement to using the Abercrombie correction^17^ that assumes no caps are lost (ϕ = 0°) or d_min_ correction^47^ (or equivalent h_min_ correction^42^) since d_min_ is not a good measure of the lost-cap distribution when ϕ > 20° (Appendix A).

**Table 4.**
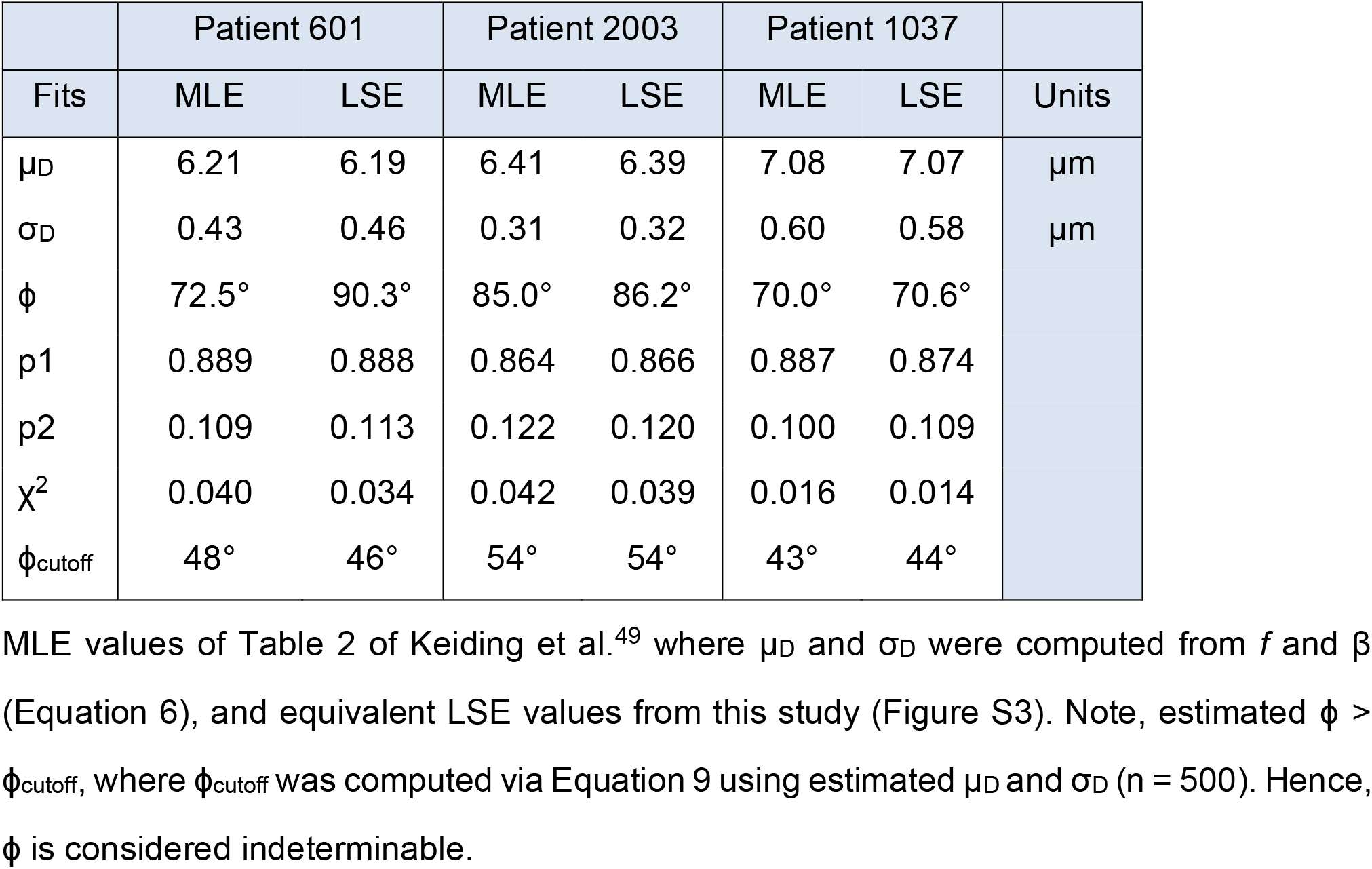
Replication of the original Keiding-model fits to G(d) of human liver cell nuclei.

### 3.4 The cap-angle limit ϕ in previous studies

Although ϕ was never reported for Wicksell’s spleen corpuscles^16^, our curve-fit analysis of Wicksell’s G(d), which produced an estimated F(d) that matched that of Wicksell, resulted in ϕ = 25° with a small fit error of 3° (Figure S4). In the Wicksell study, the corpuscles were large (μ_D_ = 323 μm) in comparison to the tissue section (18 μm), in which case T ≈ 0.06 u.d. Hence, the small estimated ϕ for Wicksell’s dataset is consistent with that of our GC nuclei in planar sections where ϕ = 20°.

Another study of GC nuclei^64^ computed a lost-caps correction factor *f* = 0.978. If one expresses *f* as a cap-angle limit, then ϕ = cos^−1^(*f*) = 12°. This ϕ is smaller than that computed for the GC nuclei in this study (20°), which is unexpected given Harvey and Napper used thin sections (T ≈ 0.3) and light microscopy compared to planar sections and TEM. It seems likely that the smaller ϕ of Harvey and Napper is due to their use of d_min_ to estimate *f*, in which case *f* is likely to be underestimated since d_min_ < d_ϕ_ with high probability (Appendix A; Table 6; Figure 3B–D).

A previous study of vesicles within mineralized cartilage matrix estimated ϕ = 60° using an unfolding method that included the Keiding model^44^. In the Hunziker study, matrix vesicles had an average diameter of 70 nm and the average section thickness was 28.5 nm^72^, in which case T ≈ 0.4 u.d. These results are consistent with those of our blind analysis of MFT vesicles in TEM images where estimated ϕ > 50°. Since ϕ was indeterminable for this blind analysis (Figure 9B), the estimate of ϕ for the matrix vesicles is likely to suffer the same problem. To improve our estimate for MFT vesicles, we performed a nonblind cap detection in planar sections of ET z-stacks to obtain ϕ < ϕ_cutoff_ (Figures 6 and S9).

The analysis of liver cell nuclei in thick sections (T ≈ 1 u.d.) by Keiding et al.^49^ resulted in estimated ϕ = 70–85° (Table 4) with a large estimation error (their Table 3). Using their estimated μ_D_, σ_D_ and N, we computed ϕ_cutoff_ ≈ 43–54° via Equation 9, indicating their estimated ϕ > ϕ_cutoff_, i.e. ϕ was indeterminable. Hence, the results of Keiding et al. also parallels that of our blind analysis of vesicles in thick sections where estimated ϕ > ϕ_cutoff_ with a large estimation error (Figure 9).

In summary, the results from this and previous studies reveal a wide range of ϕ, highlighting the fact that ϕ is highly dependent on the experimental conditions. However, thin or planar sections are best for obtaining ϕ < ϕ_cutoff_. Hence, if one wishes to accurately estimate ϕ, which is necessary for estimating λ_3D_ via Equation 3, then one should use thin or planar sections and, if possible, adopt the other recommendations for optimising cap detection discussed above.

### 3.5 Distribution-based versus distribution-free methods and the assumption F(d) is Gaussian

Both the LSE method used in this study and the MLE method of Keiding et al.^49^ are distribution-based (i.e. parametric) methods for estimating F(d) and ϕ from G(d) since both models assume a statistical model for F(d), e.g. a Gaussian, chi or gamma distribution. By contrast, distribution-free methods, also known as non-parametric methods or unfolding algorithms, have been extensively used in various scientific fields^2^. There are advantages and disadvantages to both methods and we refer the reader elsewhere for discussions^49,50,73,74^. However, the advantages of using a distribution-based rather than distribution-free method are that it is more exact and stable and does not create implausible negative probabilities for F(d). Moreover, as we have demonstrated here, a distribution-based method allows one to accurately estimate ϕ, which can subsequently be used to estimate λ_3D_ from λ_2D_ via Equation 3.

There are multiple pieces of evidence that suggest a Gaussian F(d) is valid for our samples of GC somata and nuclei and MFT vesicles. First, curve fits of the Keiding model to G(d), where F(d) is assumed to be a Gaussian distribution, showed excellent agreement for the GC somata and nuclei and MFT vesicles (Figures S12–S16). Moreover, repeating the same curve fits, but assuming a chi or gamma distribution for F(d), resulted in the same Gaussian solutions for F(d) (i.e. large *f*; Figures S14C and S16C). Finally, our curve fits of Gaussian, chi and gamma distributions to F(d) computed from ET z-stacks of MFT vesicles converged to the same Gaussian solution (Figure S8). Interestingly, the MLE fits of Keiding et al.^49^ to G(d) of liver cell nuclei, which assume a chi distribution for F(d), also converge to a Gaussian F(d) (i.e. large *f*). Our replication of the MLE fits shows that assuming either a chi or Gaussian distribution for F(d) makes little difference in the shape of the curve fits or final estimates of μ_D_, σ_D_ and ϕ (Figure S3; Table 4).

While our study focused on F(d) described by a Gaussian model, our findings should be applicable to F(d) described by other statistical models. Moreover, our numerical solutions of the Keiding model, which have been incorporated into the latest version of the analysis package NeuroMatic^53^, can be readily used as templates for creating new models that assume other distributions for F(d). They can also be adapted to fit G(d) with multiple peaks (Figure S3) as previously described^49^.

### 3.6 Comparison of the Keiding model to the disector method

While our simulations of the disector method for estimating λ_3D_ show no biases due to lost caps, as previously hypothesised^52^, this was only true if the identification of caps on the reference and lookup sections were equally probable, since any bias due to lost caps on the reference section cancelled that on the lookup section. However, nearly all disector analyses are performed sequentially, with a blind particle detection on the reference section followed by nonblind particle detection on the lookup section, in which case there is an increased probability of identifying caps on the lookup section (a scenario proposed by Hedreen^27^). When such an asymmetrical bias was added to our simulations, there was an underestimation error of λ_3D_ that was large even for small degrees of bias. Using the blind-versus-nonblind bias in particle detection measured from our analysis of MFT vesicles (ϕ_bias_ = 20°), for example, we found a large underestimation error (Δλ_3D_ = -40%; Figure 12). Interestingly, an underestimation counting error of -15% for adjacent reference and lookup sections has been previously reported and attributed to lost caps^61^. To remove the blind-versus-nonblind bias in the disector method, Hedreen^27^ suggests using a third section immediately below the reference section to guide the identification of caps in the reference section, i.e. a nonblind-nonblind particle detection.

Besides the potential underestimation error due to the blind-versus-nonblind bias in identifying caps, the low particle count of the disector method makes it inherently less accurate in estimating λ_3D_ compared to the ϕ-correction method. Our simulations indicate that, given the same number of particles per projection, the disector method is ∼2 to 3-fold less accurate than the ϕ-correction method due to the smaller counts per section. To get the same level of accuracy one would need to increase the cross-sectional area of the section (or ROI) by ∼4-fold. Hence, the ϕ-correction method for estimating λ_3D_ is potentially more efficient and accurate than the disector method.

Finally, for particles with a high density, especially those that are touching one another (e.g. cerebellar GCs and MFT vesicles), the disector method is not recommended^18^. In this case, the ϕ-correction method can be used in conjunction with planar sections. That said, it is important to keep in mind that the ϕ-correction method requires an accurate estimate of ϕ, which is not always possible for a given particle and imaging technique, and is designed for particles with a spherical geometry.

### 3.7 Comparison of the Keiding model to alternative model-based and design-based stereological methods

Since the onset of design-based stereological methods in the 1980s, there has been considerable debate about the merits of these methods in comparison to the older model-based stereological methods^20,22–30,51,75^. Much of the debate has stemmed from the large variability in estimates of 3D particle counts (i.e. λ_3D_) within studies using model-based methods, i.e. the Abercrombie^17^ or Floderus^42^ correction, or between studies using either model-based or design-based methods. Our analysis of the two methods hopefully sheds light on the source of those variabilities: there are potential biases in both model-based and design-based counting methods. The Abercrombie correction^17^ (ζ = T + μ_D_), for example, is likely to underestimate λ_3D_ since it assumes no caps are lost (ϕ = 0°), and the Floderus h_min_ correction^42^, or equivalent Konigsmark d_min_ correction^47^, is likely to underestimate λ_3D_ since it assumes all particles are the same size (Appendix A). The use of d_min_ (or h_min_) to correct for lost caps is also problematic since this measure is susceptible to being an outlier (e.g. a single false-positive measurement). Moreover, those using the Abercrombie and Floderus corrections typically use approximate measures of μ_D_, potentially adding an additional bias. As discussed above, the design-based disector method has been deemed unbiased at its conception^52^; however, our analysis of this method has confirmed that a blind-versus-nonblind bias in particle detection, as proposed by Hedreen^27^, leads to an underestimation of λ_3D_. Moreover, biases due to inaccurate measures of section thickness (T) and z-axis tissue distortions can lead to significant biases in estimated λ_3D_ for both design-based and model-based methods, as discussed above. Hence, it is not surprising that comparisons of estimated λ_3D_ between model-based and design-based methods show large discrepancies. Our analysis supports the conclusion that neither model-based or design-based methods should automatically be assumed unbiased^25,26^ and both methods need to be verified/calibrated, preferably via 3D reconstructions^24,28,29,51^. Yet, 3D reconstructions should not automatically be assumed to be the ‘gold standard’ due to potential z-axis distortions of tissue sections, as discussed above.

Here, we validated a model-based ϕ-correction method for estimating λ_3D_ that has been long overlooked since the 1980s^50^, most likely since design-based methods have become the defacto tools in modern stereology. Results of the validation show the ϕ-correction method can estimate λ_3D_ with high accuracy. A high accuracy is achieved via a superior model of the lost-cap distribution, i.e. the Keiding model, that represents the mean cap-angle limit of a population of spherical particles. Moreover, use of an LSE routine (or MLE routine) allows one to use all measured 2D diameters (i.e. G(d)) to estimate ϕ, which is a significant improvement to using d_min_, a single measure that is likely to be an outlier, and also gives an accurate estimate of μ_D_. Comparison of estimated λ_3D_ computed via the ϕ-correction method to that computed via the Abercrombie and Floderus correction methods show biases (underestimations) in the latter corrections by as much as ∼20% for our MFT vesicle dataset (Table 6).

To estimate particle size, we used outlines to compute the cross-sectional area of a particle’s projection in 2D images, which is equivalent to a high-resolution design-based point-grid method, since pixels define a grid^70^. Because particles are never perfect spheres, cross-sectional area is a better measure of 2D size than the commonly used diameter line-segment measures d_long_ and d_short_ (Table 5), and is consistent with methods for creating 3D reconstructions. Moreover, the cross-sectional areas can be used to estimate particle density via the VF = K_v_·AF relation (Equations B.1, B.2c). To estimate F(d), we computed G(d) from the equivalent diameters of the cross-sectional areas (d_area_) and curve fitted Equation 1 to G(d) using an LSE algorithm, as discussed above. This method provides accurate measures of both μ_D_ and σ_D_, even when true ϕ > ϕ_cutoff_. Hence, the Keiding model offers a simple and efficient means of accurately estimating F(d). The design-based nucleator and rotator methods, on the other hand, which are used in conjunction with the disector method^18,21^, only provide estimates of mean particle volume, are more time consuming since they require multiple line-segment measurements per particle, and are potentially less accurate due to the low particle count of the disector method.

**Table 5.** 2D diameter measures compared to d_area_.

| Particle | Image | $d_{\text{long}}$ | $d_{\text{short}}$ | $d_{\text{geometric}}$ | $d_{\text{avg}}$ |
| --- | --- | --- | --- | --- | --- |
| GC soma | Confo | +15% *** | -13% *** | -0% | +1% * |
| GC nucleus | TEM | +17% *** | -21% *** | -5% *** | -2% *** |
| MFT vesicle | TEM | +2% ** | -12% *** | -5% *** | -5% *** |
| MFT vesicle | ET11 | +0% | -20% *** | -11% *** | -10% *** |

Confo: confocal. Percent difference ( $\Delta$ ) measured with respect to $d_{\text{area}}$ . Significance measured via paired $t$ -tests in comparison to $d_{\text{area}}$ (\* $p < 0.05$ , \*\* $p < 0.01$ , \*\*\* $p < 0.001$ ). There were 50 particles per sample.

Hence, our analysis of the Keiding model demonstrates that a model-based approach to estimating the size and density of spherical particles can offer high levels of accuracy. This approach can be used in conjunction with the design-based random sampling strategies to avoid sampling biases^20,67,70,71^.

## 4 Materials and Methods

### 4.1 Transmission electron microscopy of cerebellar sections

Acute sagittal sections of the cerebellar vermis (∼200 μm thick) were prepared from 2 male and 2 female C57B6/J WT mice (P26–31; Charles River Germany, from the Jackson Laboratory; line #000664; RRID:IMSR_JAX:000664) in ice-cold high-sucrose artificial cerebrospinal fluid (ACSF; 87 mM NaCl, 25 mM NaHCO_3_, 2.5 mM KCl, 1.25 mM NaH_2_PO_4_, 10 mM glucose, 75 mM sucrose, 0.5 mM CaCl2, and 7 mM MgCl_2_, equilibrated with 95% O_2_ and 5% CO_2_, 325 mOsm) using a Leica microsystems vibratome (VT1200S) as previously described_3_. Sections were allowed to recover in high-sucrose ACSF at 35°C for 30–45 min, then in normal ACSF (125 mM NaCl, 25 mM NaHCO_3_, 25 mM D-glucose, 2.5 mM KCl, 1.25 mM NaH_2_PO_4_, 2 mM CaCl_2_, and 1 mM MgCl_2_, equilibrated with 5% CO_2_ and 95% O_2_) at room temperature (∼23°C).

To prepare the cerebellar sections for high-pressure freezing, sections were heated to 37°C for 5–10 minutes in ACSF, then mounted into a sample ‘sandwich’ on a table maintained at 37°C. The sample sandwich was assembled by placing a 6 mm sapphire disk on the middle plate of a transparent cartridge system, followed by a spacer ring, the section, a drop of ACSF containing 15% of polyvinylpyrrolidone for cryoprotection and adhesion, another sapphire disk and finally a spacer ring. The sample sandwich was frozen via a Leica EM ICE high-pressure freezing machine.

Freeze substitution of the frozen samples was performed using an AFS1 or AFS4 Leica system equipped with an agitation module^76^. While in liquid nitrogen, frozen samples were transferred from storage vials to freeze-substitution vials containing 0.1% tannic acid and acetone, previously frozen in liquid nitrogen. Vials were transferred to the AFS1/AFS4 system and shaken for 22–24 hours at -90°C. Inside the AFS1/AFS4 system, samples were washed for 10 minutes in pre-chilled acetone at -90°C for 3–4 repetitions. Next, a contrasting cocktail with 2% osmium and 0.2% uranyl acetate in acetone was chilled to -90°C and added to each vial. The temperature of the vials was kept at -90°C for 7–10 hours, raised to -60°C within 2 hours (15°C/hour), kept at -60°C for 3.5 hours, raised to -30°C within 4 hours (7.5°C/hour), kept at -30°C for 3.5 hours, raised to 0°C within 3 hours (10°C/hour), kept at 0°C for ∼10 min, then transferred to ice where samples were washed with acetone (3 × 10 min). Samples were transferred from the vials to glass dishes containing acetone at room temperature and inspected for intactness and proper infiltration. Samples were washed with propylene oxide (2 × 10 min) and infiltrated with Durcupan resin at 2:1, 1:1 and 1:2 propylene oxide/Durcupan resin mixtures (1 hour at room temperature). Samples were left in pure resin overnight at room temperature, embedded in BEEM capsules (Electron Microscopy Sciences, Hatfield, PA, USA) and allowed to polymerize over a second night at 100°C. Samples were trimmed with glass knives and cut into ultrathin (∼60 nm) sections via a Leica EM UC7 Ultramicrotome with Diatome Histo diamond knife (6 mm, 45°). Sections were placed in formvar-coated slot grids and post-stained in 2% uranyl acetate for 10 minutes, then lead citrate for 2 minutes. Sections were imaged via a transmission electron microscope (FEI Tecnai 10, 80 kV accelerating voltage) with an OSIS Megaview III camera and Radius acquisition software.

Mice were bred in a colony maintained in the preclinical animal facility at IST Austria. All procedures strictly complied with IST Austria, Austrian, and European ethical regulations for animal experiments, and were approved by the Bundesministerium für Wissenschaft, Forschung und Wirtschaft of Austria (BMWFW-66.018/0010-WF/V/3b/2015 and BMWFW-66.018/0008-V/3b/2018).

### 4.2 Electron tomography of cerebellar sections

One male C57Bl6 WT mouse (P30) was anaesthetized with ketamine and transcardially perfused with 2% paraformaldehyde and 1% glutaraldehyde in 0.1 M Na-acetate buffer for 2 min, then 2% paraformaldehyde and 1% glutaraldehyde in 0.1 M Na-borate buffer for one hour. After perfusion, the mouse’s brain was dissected and 60 μm sections were cut from the cerebellar vermis. Sections were treated with 1% OsO_4_, stained in 1% uranyl acetate, dehydrated in a graded series of ethanol and embedded in epoxy resin (Durcupan). From the embedded sections, serial sections ∼200 nm thick were cut with a Leica Ultramicrotome EM UCT and collected onto copper slot grids, where fiducial markers were introduced at both sides of the grids. Single-axis tilt series were acquired via an FEI Tecnai G2 Spirit BioTWIN transmission EM (0.34 nm line resolution) operating at 120 kV and equipped with an Eagle 4K HS digital camera (FEI, Eindhoven, The Netherlands). Tilt series were recorded between ±65° (with 2° increments between ±45°, then 1° increments) at 30,000× magnification (R_xy_ = R_z_ = 0.38 nm/voxel) using FEI Xplore3D. Tomographic subvolumes were reconstructed using IMOD (RRID:SCR_003297) and exported as z-stack images (R_xy_ = 1.14 nm/pixel). Two different z-stacks of MFT vesicles were analysed in this study, denoted ET10 and ET11.

The mouse was housed in the vivarium of the Institute of Experimental Medicine in a normal 12 hour/12 hour light/dark cycle and had access to water and food ad libitum. The experiment was carried out in accordance with the Hungarian Act of Animal Care and Experimentation 40/2013 (II.14) and with the ethical guidelines of the Institute of Experimental Medicine Protection of Research Subjects Committee.

### 4.3 Analysis of 2D projections

From 2D images, outlines of particles (i.e. somata, nuclei and vesicles) were drawn using Fiji’s freehand tool^77^ (RRID:SCR_002285; https://imagej.net/Fiji) and an equivalent diameter (d) was computed from the area of each outline (d_area_ = 2(area/π)^½^)^56,58^. To avoid introducing bias by pooling data from multiple researchers^54^, outlines were drawn by a single author (JSR). To avoid selection bias, e.g. outlining only the largest particles, an attempt was made to outline all visually identifiable particles within each selected ROI. Histograms of equivalent-area diameters, G(d), were computed as counts per bin, then normalised to give a probability density by dividing the count within each bin by the total number of diameters times the bin size. Images and associated analyses are denoted with identification (ID) tags for the rat confocal images (R1, R5, R6) and mouse TEM images (M15, M18, M19, M21).

A numerical approximation for G(d) as defined in Equation 1 was computed via Igor Pro (RRID:SCR_000325; WaveMetrics, Portland, Oregon) where the integral in this equation was solved via an adaptive Gaussian quadrature integration routine (Integrate1D), avoiding the singularity in the denominator by setting the denominator to 1 × 10^−7^ when d = δ. The same numerical approximation was used to curve fit Equation 1 to our simulated and experimental G(d) via Igor Pro’s CurveFit operation, using the Levenberg-Marquardt LSE algorithm. F(d) in Equation 1 was assumed to be a Gaussian function (Equation 5) unless specified. The estimated error of each fit parameter is reported as ±1 standard deviation (±σ). During the fit routine, parameter T was fixed at its estimated value. The initial guess for ϕ was set to θ_min_, where θ_min_ = sin^−1^(d_min_/μ_D_) and d_min_ is the smallest non-zero diameter bin of G(d). Initial guesses for μ_D_ and σ_D_ were set to μ and σ of G(d), where μ of G(d) was computed as the sum of d·G(d)·dx over all bins and σ^2^ was computed as the sum of (d - μ)^2^·G(d)·dx. For a small number of fits to the simulated G(d), usually for conditions of true ϕ > ϕ_cutoff_, initial guesses had to be adjusted to get a successful fit. No parameters were constrained during the fits (e.g. 0 ≤ ϕ ≤ 90°) since testing of the LSE routine using a variety of datasets showed such constraints were never active or violated during the test fits. To validate the LSE routine, the MLE fits of Keiding et al.^49^ were replicated, showing nearly identical results (Figure S3; Table 4). Likewise, an LSE fit to Wicksell’s G(d) of spleen corpuscles^16^ resulted in an estimated F(d) that was nearly the same to that of Wicksell’s unfolding solution (Figure S4).

The distribution of lost caps, L(d), was computed from G(d) via Equation 1 for T = 0 u.d. as follows: L(d, ϕ) = G(d, ϕ = 0°) - G(d, ϕ), where G(d, ϕ = 0°) and G(d, ϕ) were computed over the range d = 0–3 u.d. and G(d, ϕ) was normalised so that its last data point at d = 3 u.d. equaled that of G(d, ϕ = 0°).

To compute λ_2D_ from a 2D image using Fiji, a rectangular ROI was defined within a distribution of the particles of interest and two adjacent borders were designated as inclusive and the other two as exclusive (Figure S1). Particles were counted if they touched the inclusive borders or were completely contained within the ROI, and not counted if they touched the exclusive border^78^. λ_2D_ was computed as the particle count (N_2D_) divided by the ROI area. Using λ_2D_, μ_D_ and ϕ, λ_3D_ was estimated via Equation 3, in which case it was important that λ_2D_ was computed from the same image (or z-stack) from which μ_D_ and ϕ were estimated, since there was variation in μ_D_ and ϕ between sections. To compute the particle VF for a given λ_3D_ and F(d), the following expression was used:

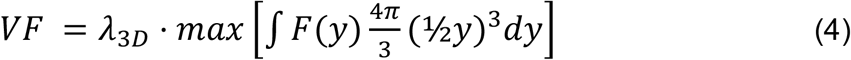

where F(d) is a PDF (Equation 5), max is the maximum value of the set of values and *y* is the variable of integration.

Measurements of diameters and density were analysed using NeuroMatic^53^ (RRID:SCR_004186; Key Resources), an acquisition, analysis and simulation tool that runs within the Igor Pro environment. Functions for Equation 1 have been incorporated into the latest version of NeuroMatic which can be accessed via NeuroMatic’s analysis Fit tab, or Igor Pro’s analysis Curve Fitting graphical user interface or Global Fit package. These functions (NMKeidingGauss, NMKeidingChi and NMKeidingGamma) assume either a Gaussian, chi or gamma PDF for F(d) (Equations 5, 6, 7) and can be readily used as templates for creating new Keiding models that assume other PDFs.

### 4.4 Probability Density Functions (PDFs)

PDFs (e.g. F(d)) were described by either a Gaussian, chi or gamma distribution. The Gaussian distribution was as follows:

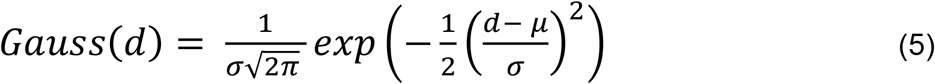

where d is the independent variable (e.g. diameter), and μ and σ are the mean and standard deviation of the distribution. The chi distribution was the same as that used by Keding et al.^49^:

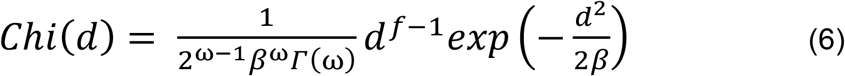

where *f* denotes the number of degrees of freedom, ω = ½f, β the scale parameter and Γ the gamma function. Given *f* and β, one can compute the distribution’s μ = γ(2β)^½^ and σ^2^ = β(*f* - 2γ^2^) where γ = Γ[½(*f* + 1)]/Γ(½*f*) (Equation 5.2 of Keiding et al.). The gamma distribution was as follows:

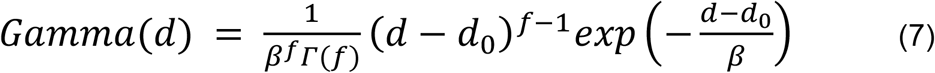

where d_0_ is an x-axis offset parameter added for flexibility. Given *f* and β, one can compute the distribution’s μ = d_0_ + *f*·β and σ^2^ = *f*·β^2^. Note, both the chi and gamma distribution converge to a Gaussian distribution as *f* → ∞. Hence, a large *f* indicates a Gaussian-like distribution.

### 4.5 Monte Carlo simulations

2D projections of spherical particles were simulated using D3D, a reaction-diffusion simulation package that includes a Monte Carlo algorithm for distributing non-overlapping hard spheres in arbitrary 3D geometries^5^ (Key Resources). Spherical particles were randomly distributed in a rectangular cuboid using periodic boundary conditions (Figure S2). The xy-square dimensions of the cuboid were adjusted to accommodate the required number of particles per projection, and the z-dimension was adjusted to accommodate the required number of projections. The particle VF = 0.40 unless specified. 3D particle diameters (D) were randomly drawn from a Gaussian distribution for a given F(d). Projections in the xy-plane were computed by identifying those particles with their center point located within a given section (interior particles), and those with their center point located above or below the section (caps) at a distance dz, where dz < D/2. The 2D projected diameter (d) was computed as d = D for an interior particle and d = (D^2^ - 4dz^2^)^½^ for a cap, derived from the trigonometric relation: (½D)^2^ = (½d)^2^ + dz^2^. A particle’s cap angle was computed as θ = sin^−1^(d/D). Other measures computed for each particle were the particle’s distance from the section surface and the sum of overlaps within the xy-projection (Ω; circle-circle overlaps expressed as a fraction) between the particle and particles higher in the section. To simulate lost caps, particles were excluded from the projection if θ was less than a fixed lower limit (ϕ), as in the Keiding model^49^. For a few simulations, however, ϕ was not fixed but variable, in which case particles were assigned a ϕ randomly drawn from a Gaussian distribution (μ_ϕ_ ± σ_ϕ_) for a given CV_ϕ_. To simulate the inability to observe particles deep in the section due to overlapping projections, particles were excluded from the projection if their Ω was greater than a fixed upper limit (ψ). To simulate the merging of circular projections for opaque particles, the xy-distance between two projections was computed according to the α-parameter of Hilliard^57^: α = (½d_1_·d_2_)(4d_12_^2^ - d_12_ - d_22_), where d_1_ and d_2_ are the projection diameters and d_12_ is the distance between the projection center points; if -1 < α < 0, then the projections were merged into one, resulting in a decrease in projection count and increase in projection size. The procedure for merging projections began with the particle closest to the section surface, which was then merged with other particles if -1 < α < 0. The merging procedure continued with the next particle closest to the section surface, and so on. To compute λ_2D_, the number of particles in a projection was divided by the geometry Area_xy_. Because periodic boundary conditions were used in the simulations (Figure S2), this λ_2D_ is equivalent to one computed using inclusive/exclusive rectangular borders for counting (Figures S1).

To simulate the disector method of computing density^18,52^, particles with a Gaussian F(d) were randomly distributed within a cuboid geometry whose xy-square dimensions were adjusted to accommodate ∼500 particles per projection, and the z-dimension was adjusted to accommodate 100 sections with T = 0.3 u.d. To compute the density of a given section, a projection was computed for that section (the reference projection) as well as an adjacent section of equal thickness (the lookup projection). Particles were counted if they appeared within the reference projection but not the lookup projection. λ_2D_ was computed as particle count per Area_xy_ and λ_3D_ = λ_2D_/T. Simulations included parameter ϕ, i.e. particles were excluded from a given projection if their θ < ϕ. To simulate a blind-versus-nonblind bias in vesicle detection, ϕ was defined separately for the reference section (ϕ_ref_) and lookup section (ϕ_lookup_) such that ϕ_ref_ ≥ ϕ_lookup_, with their difference (bias) defined as ϕ_bias_ = ϕ_lookup_ - ϕ_ref_.

To quantify ϕ_cutoff_, the estimation error Δϕ (estimated ϕ - true ϕ) was computed from curve fits of Equation 1 to simulated G(d) for planar sections (T = 0 u.d.) and true ϕ = 10–80° (5° steps) over a range of CV_D_ (0.04–0.17) and number of diameters (n = 200–2000; Figures 4C and S5). ϕ_cutoff_ for a given CV_D_ and n was defined as the upper limit of true ϕ for when |Δϕ| ≤ 5° occurs with at least 0.68 probability (i.e. at least 68 out of 100 simulation repetitions). To derive an expression relating ϕ_cutoff_ to CV_D_ and 1/√n, a 2D matrix was constructed for the equivalent unit diameters of ϕ_cutoff_ (d_cutoff_ = sinϕ_cutoff_), with the row and column dimensions defining CV_D_ and 1/√n, and a bivariate polynomial with 4 dependent variables was curve fitted to the d_cutoff_ matrix in Igor Pro. The inverse sine of the curve-fit solution was as follows:

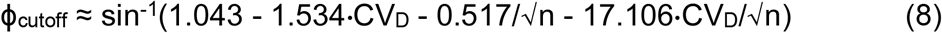

where CV_D_ is computed using true μ_D_ and σ_D_. To investigate whether this expression can be used to test the accuracy of estimated ϕ, ϕ_cutoff_ was computed using estimated μ_D_ and σ_D_ of each simulation (rather than true μ_D_ and σ_D_) and this ‘estimated’ ϕ_cutoff_ was compared to the corresponding estimated ϕ. Results showed that the use of estimated μ_D_ and σ_D_ to compute CV_D_ translated into negative offsets in ϕ_cutoff_, ranging from -17° to -10° for n = 200 to 2000 diameters, respectively. To account for these offsets, diameters in the d_cutoff_ matrix were adjusted to remove the offsets and the matrix was refit to the bivariate polynomial, resulting in:

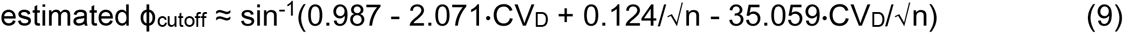

where CV_D_ is computed using estimated μ_D_ and σ_D_. In conjunction with the fit error of ϕ, this expression was used as an accuracy test of estimated ϕ, i.e. estimated ϕ was considered accurate if it was less than estimated ϕ_cutoff_. For the analysis of ET z-stacks (ET10 and ET11; Figures 6, 11, S9) and simulated z-stacks (Figures S10, S18), n in Equations 8 and 9 was reduced 3-fold to account for the reduction in sampling of F(d), a factor determined via simulations.

### 4.6 3D analysis of electron tomography reconstructions

For the size and density analysis of MFT vesicles using ET z-stacks (ET10 and ET11), vesicles were tracked and outlined through multiple planes of the z-stacks, and the equivalent xy-radii of each outline (r = ½d_area_) was computed as a function of the z-stack

image number (z_#_). The 3D diameter (D) and z-axis center point (z_0_) of each vesicle was then estimated by curve fitting the vesicle’s r-z_#_ relation to the following expression for a circle (which defines spherical dimensions):

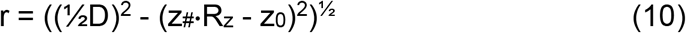

where R_z_ is the axial resolution, i.e. distance between z-stack images (Figure S7A). Because tissue shrinkage in the axial axis can be significant, R_z_ was considered unknown and estimated via a simultaneous curve fit of all vesicle r-z_#_ relations to Equation 10 while sharing parameter R_z_ (Figure S7B). To quantify lost caps, ϕ = sin^−1^(δ_min_/D) was computed for the positive and negative pole of each vesicle (if the pole was interior to the z-stack) where δ_min_ was the smallest d_area_ measurement near a given pole. For ET10, one giant vesicle ∼69 nm in diameter was excluded from the analysis. For ET11, 7 vesicles were excluded from the 3D analysis since they had a non-spherical shape.

To estimate the resolution of the ET z-stacks, the resolution formula of Crowther et al.^79^ was used to estimate ρ_x_ as follows:

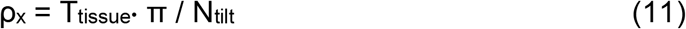

where T_tissue_ is the tissue thickness and N_tilt_ is the number of scan tilts. For ET10, T_tissue_ = 176 nm and N_tilt_ = 87. Results gave ρ_x_ = 6.3 nm. However, the Crowther formula assumes a total scan angle of 180°, and the total scan angle was 130° (±65°) for the ET scans used in this study. Hence, the Crowther formula was expressed with respect to the tilt increment (Δ_tilt_) as follows:

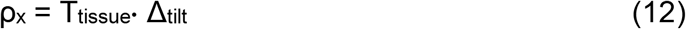

where Δ_tilt_ is the total scan angle divided by N_tilt_ (IMOD Tomography Guide). This modified formula gave ρ_x_ = 4.6 nm. Due to the ‘missing wedge effect’, the resolution in the axial axis (ρ_z_) is expected to be longer than ρ_x_ by the following scale factor:

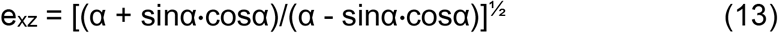

where α is the maximum scan angle^80^. For this study, α = 65° in which case e_xz_ = 1.4. Hence, ρ_z_ = ρ_x_·e_xz_ = 6.5 nm. For ET11, where T_tissue_ = 133 nm, ρ_x_ = 3.5 nm and ρ_z_ = 4.9 nm. Next, ρ_z_ was estimated from experimental data by curve fitting Equation 1 to G(d) computed from the MFT vesicle analysis (Figures 6D and S9D) while fixing μ_D_, σ_D_ and ϕ to their ‘true’ values measured from the 3D analysis and leaving T as the one free parameter. Results gave estimated T = 5.3 ± 0.8 nm for ET10 and 0.2 ± 0.4 nm for ET11. Hence, estimates of T (i.e. ρ_z_) of the ET z-stacks spanned 0–6 nm. Given the small range of estimated T, and comparatively large dimensions of the MFT vesicles, the ET analysis was simplified by assuming T = 0 nm. To test what effect this assumption might have estimates of μ_D_, σ_D_ and ϕ, the curve fits in Figures 6D and S9D were recomputed assuming T = 6 nm for ET10 and T = 5 nm for ET11 and found Δμ_D_, Δσ_D_ and Δϕ were similar to those for assuming T = 0 nm: ET10: Δμ_D_ = -0.4 vs. +1.3%, Δσ_D_ = -1.2 vs. -4.9% and Δϕ = -1.1 vs -0.4°; ET11: Δμ_D_ = -1.3 vs. +0.3%, Δσ_D_ = +8.0 vs. +0.7% and Δϕ = -3.0 vs. -2.9°.

To estimate vesicle density within a cluster for ET10, it was necessary to confine the density analysis to a subregion of the original z-stack along the axial axis since the vesicle density as a function of z-depth was nonhomogeneous, being smaller at the top and bottom of the stack (Figure 11B). Within this subregion, we estimated λ_2D_ by counting the number of outlines that fell within a ROI (Area_xy_ = 0.039 μm^2^) using inclusive/exclusive borders (Figure S1C) at the center of the vesicle cluster for 10 z-stack images spaced 10–15 nm apart along the axial axis, giving λ_2D_ = 304.2 ± 15.5 μm^−2^ (±SEM). There was an average of 12 vesicles per ROI, which is 7-fold larger than the theoretical optimal number of particles for the disector method ^30^. Using the same ROI and vesicle outlines, we computed AF = 0.45 ± 0.02 (±SEM) and VF = 0.49 ± 0.03 (VF = K_v_·AF, where K_v_ = 1.09; Equations B.1, B.2c). To estimate λ_3D_, we divided the number of vesicles counted within the z-stack subregion (115 vesicles) by the sampling space of the volume of interest (Figures 1B and 11A): VOI = Area_xy_·ζ = 0.013 μm^3^, where Area_xy_ = 0.091 μm^2^ and ζ = 143.0 nm (Equation 2; 3D measures: T = 109 nm, μ_D_ = 45.6 nm, ϕ = 41°). Here, Area_xy_ was the ROI area scaled to the equivalent xy-dimensions of the vesicle cluster, i.e. scale factor = (count per image) / (count per ROI) = 28/12 = 2.33). Results gave λ_3D_ = 8804 μm^−3^ with equivalent VF = 0.45 (Equation 4). Hence, the VF estimated via the 2D analysis is similar to that estimated via the 3D analysis.

To estimate vesicle λ_3D_ for ET11 via the ‘physical’ disector method, a reference section with T = 12.8 nm (0.3 u.d.) was randomly located within the center of the z-stack and a corresponding adjacent lookup section with the same T was defined. Vesicles that appeared in the reference section (i.e. vesicles that had one or more of their 2D outlines from the nonblind analysis in Figure 6 appear in the reference section) but not the adjacent lookup section were counted and used to compute λ_3D_ = count/(Area_xy_·T), where Area_xy_ = 0.144 μm^2^. This analysis resulted in ∼20 vesicles per section, or λ_3D_ ≈ 11,000 μm^−3^. To simulate a bias between a blind reference vesicle detection and nonblind lookup vesicle detection (ϕ_bias_), vesicle outlines from the nonblind analysis were used for the lookup vesicle detection (mean ϕ_lookup_ = 42°) and a copy of the same outlines for the reference vesicle detection, but modified to have a larger ϕ (ϕ_ref_ = ϕ_lookup_ + ϕ_bias_) by deleting the necessary number of extreme outlines from the negative and positive pole regions to achieve the desired ϕ_ref_. Setting ϕ_bias_ = 17° (Figure 7A) resulted in ∼14 counts per section, or λ_3D_ ≈ 7000 μm^−3^, with estimation error Δλ_3D_ = -32%. To compare these results to those of the Monte Carlo disector simulations, results of 7–9 reference sections as just described were combined to give a total of ∼500 vesicles per reference section for a given ϕ_bias_, and an average Δλ_3D_ was computed from 100 such reference sections.

### 4.7 Statistics

Comparisons between diameter distributions were computed via a Kolmogorov-Smirnov (KS) test (significant p < 0.05). Other comparisons were computed via a Student’s *t*-test where noted (unpaired two-tailed equal-variance, unless specified differently; significant p < 0.05; F-test used to verify equal variance). Errors reported in the text and graphs (bars/shading) indicate the standard deviation (±σ), except in a few instances they indicate the standard error of the mean (±SEM) which is noted.

The estimation error (Δ) of parameters μ_D_, σ_D_, λ_2D_ or λ_3D_ was computed as the percent difference between a parameter’s estimated (ε) and true (t) value [Δ = 100(ε - t)/t], except for ϕ, which was computed as a difference (Δ = ε - t) since division by ϕ caused distortion at small ϕ. For Monte Carlo simulations with multiple repetitions, the mean and standard deviation of a parameter’s estimation error (μ_Δ_ ± σ_Δ_) is referred to as the bias and (68%) confidence interval, respectively. For simulations with true ϕ < ϕ_cutoff_, the Δ-distributions were typically normal. However, for simulations with true ϕ ≥ ϕ_cutoff_, the Δ-distributions were often skewed (i.e. absolute skew > 0.5); in this case, μ_Δ_ was computed as the median of the Δ- distribution and +σ_Δ_ and -σ_Δ_ were computed separately above and below μ_Δ_.

## Appendix A. Derivation of Equation 3 and why d_min_ is not a good measure of lost caps

Abercrombie^17^ derived the following relation between the expected true particle count (N_true_) and actual crude particle count (N_measured_) of a section of thickness T, also known as the Abercrombie correction formula:

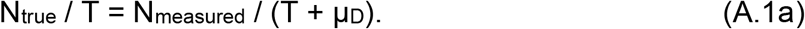

Dividing both sides of this equation by the cross-sectional area over which the particles are counted (Area_xy_) gives:

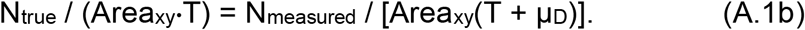

Substituting in terms λ_3D_ and λ_2D_ used in this study gives:

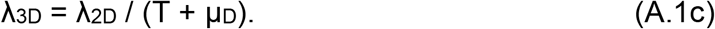

However, this relation is only correct if there are no lost caps. If there are lost caps, then μ_D_ must be scaled smaller. Using the Keiding fixed-ϕ model, scaling of μ_D_ can be achieved via cosϕ (Figure 1) and λ_3D_ can be estimated via the following^50^:

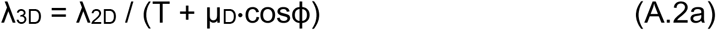

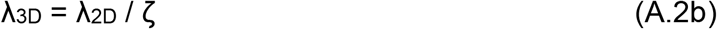

which is Equation 3. Floderus^42^ derived a similar expression with respect to the section penetration depth of the smallest observable cap (h_min_):

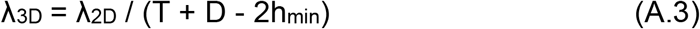

and this expression was later recast by Konigsmark^47^ as a function of d_min_, the diameter of the smallest observable cap:

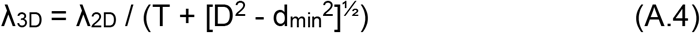

using the trigonometric relation:

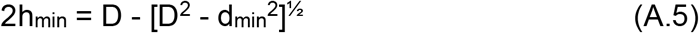

For analysing images, Equation A.4 has proved more useful than the original Floderus equation since d_min_ can be directly measured from the sample of measured 2D diameters. However, an underlying assumption of the Floderus/Konigsmark correction is that all particles are the same size. If there is a distribution of particle size, then each particle will have its own minimum observable diameter (δ_min_) and d_min_ will be the minimum of all δ_min_ (Figures 6C1 and S9C1). Depending on the spread of the δ_min_ distribution, the difference between d_min_ and the mean δ_min_ can be large, in which case the d_min_ correction will create a large underestimation of λ_3D_. Moreover, there is a high probability that d_min_ is an outlier, i.e. an unusually small value; this could happen when conditions for identifying small caps are better than average, or when there are false positive measurements.

Our 3D analysis of MFT vesicles showed the Keiding model accurately describes the δ_min_-D relation of the vesicles (Figure 6C1 and S9C1) and therefore gives an accurate estimate of λ_3D_ via Equation 3 (Figure 11). This is in contrast to the d_min_ correction which underestimated λ_3D_ for the same datasets by 13–20% and the Abercrombie correction which underestimated λ_3D_ by 21–23% (Table 6; ET10 and ET11). However, under ideal conditions where there are few lost caps, differences between λ_3D_ estimated via the Keiding model and the d_min_ and Abercrombie corrections are expected to be small. For example, our analysis of GC nuclei in planar sections, where ϕ = 20°, showed only a 2–6% difference between estimated λ_3D_.

**Table 6.** Comparison of λ_3D_ estimated via the Keiding model, Abercrombie correction or Konigsmark (d_min_) correction.

| | | | $\zeta = T + \mu_D \cdot \cos\theta_{\text{min}}$ | | | $\zeta = T + \mu_D$ | |
| --- | --- | --- | --- | --- | --- | --- | --- |
| Particle | Image | $\phi$ | $\theta_{\text{min}}$ | $\Delta\zeta$ | $\Delta\lambda_{3D}$ | $\Delta\zeta$ | $\Delta\lambda_{3D}$ |
| GC soma | Confo | 37° | 27° | +9% | -8% | +19% | -12% |
| GC nucleus | TEM | 20° | 17° | +2% | -2% | +6% | -6% |
| MFT vesicle | ET10 | 41° | 18° | +27% | -20% | +33% | -23% |
| MFT vesicle | ET11 | 41° | 27° | +19% | -15% | +33% | -21% |

Confo: confocal. Mean $\phi$ from Tables 2 and 3. $\theta_{\text{min}} = \sin^{-1}(d_{\text{min}}/\mu_D)$ , used in Equations 2 and 3 as a substitute for $\phi$ . Right column: Abercrombie correction<sup>17</sup>, which assumes no lost caps ( $\theta_{\text{min}} = 0^\circ$ ). $\Delta = 100[X(\theta_{\text{min}}) - X(\phi)]/X(\phi)$ where $X = \zeta$ and $\lambda_{3D}$ . For vesicles, comparisons are with respect to 'true' $\zeta$ and $\lambda_{3D}$ computed via 3D reconstructions. See Appendix A.

## Appendix B. Estimation of the volume fraction of spherical particles from the area fraction of their 2D projections

The relation between the volume fraction (VF) of spherical particles and their observed area fraction (AF) in a 2D projection was derived by Weibel and Paumgartner^60^ (their Equations 13 and 37) and is as follows:

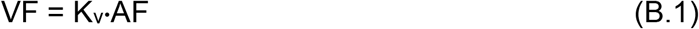

where

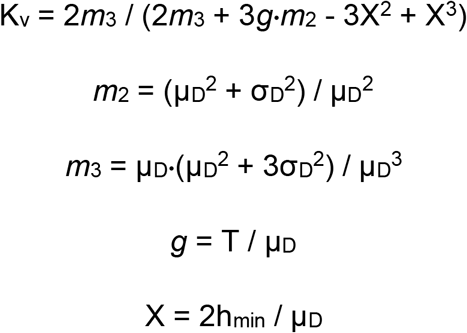

Here, *m*_^2^_ and *m*_^3^_ are dimensionless moments for a Gaussian distribution and μ_D_ = 2μ_R_. Because ϕ is a better descriptor of the lost-cap distribution than h_min_ or its equivalent d_min_ (Appendix A; Figures 6C1 and S9C1), we express X as a function of ϕ rather than h_min_ by first defining X with respect to d_min_ via Equation A.5:

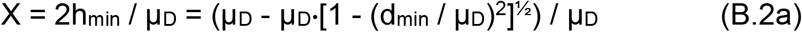

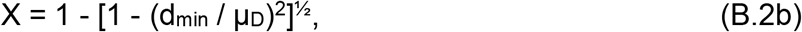

and then substitute d_ϕ_ = μ_D_·cosϕ for d_min_:

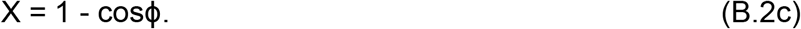

To test this relation, we can consider the ideal scenario of a planar section with no lost caps (ϕ = 0°), in which case *g* = 0, X = 0, K_v_ = 1 and VF = AF, which is expected^60^. We can also consider the scenario where all particles have the same diameter (D), in which case σ_D_ = 0, *m*_2_ = *m*_3_ = 1 and:

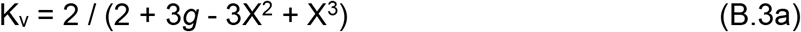

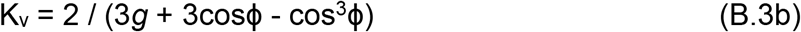

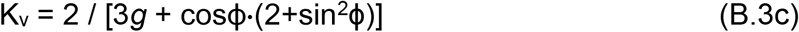

which is equivalent to Equation 51 of Weibel and Paumgartner if one substitutes d_ϕ_ for d_min_ (i.e. d_min_/D → d_ϕ_/D = sinϕ). Finally, we computed the VF of the Monte Carlo simulations in Figure 4C using Equations B.1, B.2c and found close agreement to the true VF (Figure 10A3). Likewise, the VF computed via the same equations for our ET z-stack analysis showed close agreement to the VF computed via our 3D analysis (Figure 11; Table 3).

## Key Resources

D3D^5^ https://github.com/SilverLabUCL/D3D

NeuroMatic^53^ http://NeuroMatic.ThinkRandom.com

https://github.com/SilverLabUCL/NeuroMatic

Igor Pro https://www.wavemetrics.com/

IMOD Tomography Guide https://bio3d.colorado.edu/imod/doc/tomoguide.html

## Definition of Key Terms

T: thickness of tissue section (transmission microscopy) or focal plane (ρ_z_)
D: 3D diameter of a particle
μ_D_ ± σ_D_: mean and standard deviation of 3D particle diameters
CV_D_: σ_D_ / μ_D_
u.d.: unit diameter, length normalised to μ_D_ (e.g. T/μ_D_)
planar: T < 0.1 u.d.
thin: T ≈ 0.3 u.d.
thick: T ≥ 1 u.d.
d: observed 2D diameter of a particle
d_min_: minimum 2D diameter of a sample of particles^47^
h_min_: minimum penetration depth of a sample of particles^42^
δ_min_: minimum 2D diameter of a given particle (z-stack analysis)
d_area_: equivalent-area 2D diameter: d_area_ = 2(area/π)^½^
d_short_, d_long_: short and long-axis 2D diameter
d_geometric_: (d_short_·d_long_)^½^
d_avg_: ½(d_short_ + d_long_)
μ_d_ ± σ_d_: mean and standard deviation of 2D diameters
F(d): probability density of 3D diameters
G(d): probability density of 2D diameters L(d) probability density of lost caps
θ: particle cap angle from section surface: sinθ = d/D where 0 ≤ θ ≤ 90
θ_min_: equivalent cap angle of d_min_: sinθ_min_ = d_min_/D
ϕ: lower limit of θ where 0 ≤ ϕ ≤ 90
ϕ_cutoff_: upper cutoff limit of when ϕ is determinable
μ_ϕ_ ± σ_ϕ_: mean and standard deviation of ϕ
CV_ϕ_: σ_ϕ_ / μ_ϕ_
d_ϕ:_: equivalent 2D diameter of ϕ d_ϕ_ = μ_D_·sinϕ
ζ: section z-depth over which particle center points are sampled
Area_xy_: ROI xy-area over which particles are counted
VF: Particle volume fraction within a volume of interest
AF: Particle area fraction within a ROI
N_3D_: Particle count within a volume of interest
N_2D_: Particle count within a ROI
λ_3D_: 3D particle density, λ_3D_ = N_3D_ / Volume_xyz_
λ_2D_: 2D particle density, λ_2D_ = N_2D_ / Area_xy_
Ω: sum of projection overlaps for a given particle where Ω ≥ 0
ψ: upper limit of Ω, i.e. 0 ≤ Ω ≤ ψ
χ^2^: sum of squared differences between data and fits (or simulations)
Δ: Parameter estimation error: % difference or difference from true value
μ_Δ_ ± σ_Δ_: bias and (68%) confidence interval of a parameter’s estimation error
ρ_xyz_: Microscope resolution
R_xyz_: Image/z-stack resolution

## Acknowledgements

We thank the IST Austria Electron Microscopy Facility for technical support, and Diccon Coyle, Andrea Lőrincz and Zoltan Nusser for their helpful comments and discussions.

## TABLES AND FIGURES

**Figure S1.**
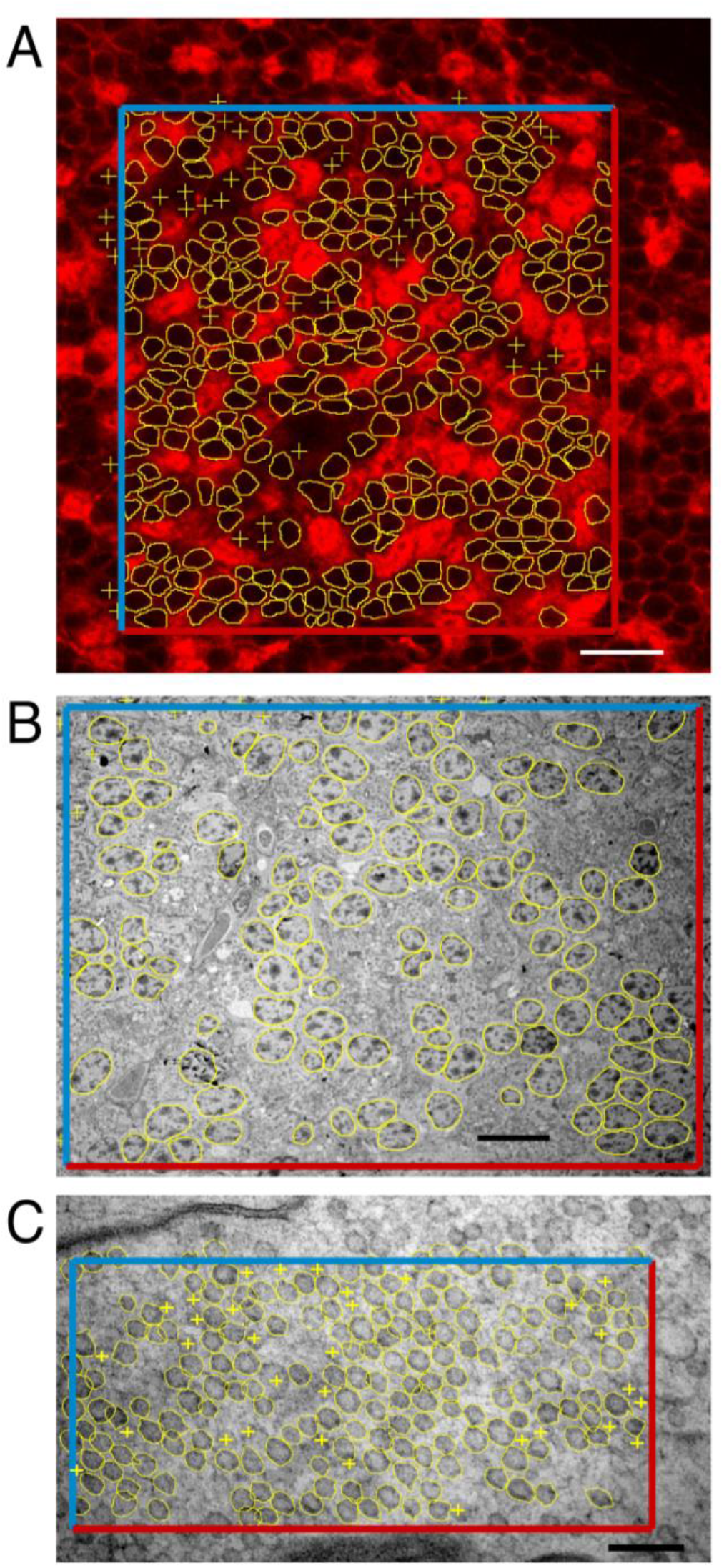
Estimating the 2D density of GC somata and nuclei and MFT vesicles from 2D images. (A) To compute λ_2D_ of GC somata of rats in a confocal image, a rectangular ROI (119.0 × 126.3 μm) was drawn within the GC layer and two adjacent borders were designated as inclusive and the other two as exclusive_78_ (blue and red). Somata were counted if they touched the inclusive borders or were completely contained within the ROI; they were not counted if they touched the exclusive borders. λ_2D_ = 21,354 mm^−2^, computed as count (321) per ROI area (15032 μm^2^). Somata were outlined if they were well delineated, otherwise they were marked via crosses (yellow). Scale bar 20 μm. Image ID R5.SL2.1. (B) Same as (A) for GC nuclei of mice in a TEM image. ROI = 88.0 × 65.3 μm. n = 130 nuclei. λ_2D_ = 22,619 mm^−2^. Scale bar 10 μm. Image ID M18.N2.51. (C) Same as (A) for MFT vesicles of mice in a TEM image. ROI = 1147 × 561 nm. n = 209 vesicles. λ_2D_ = 324.5 μm^−2^. Scale bar 150 nm. Image ID M15.L1.48.

**Figure S2.**
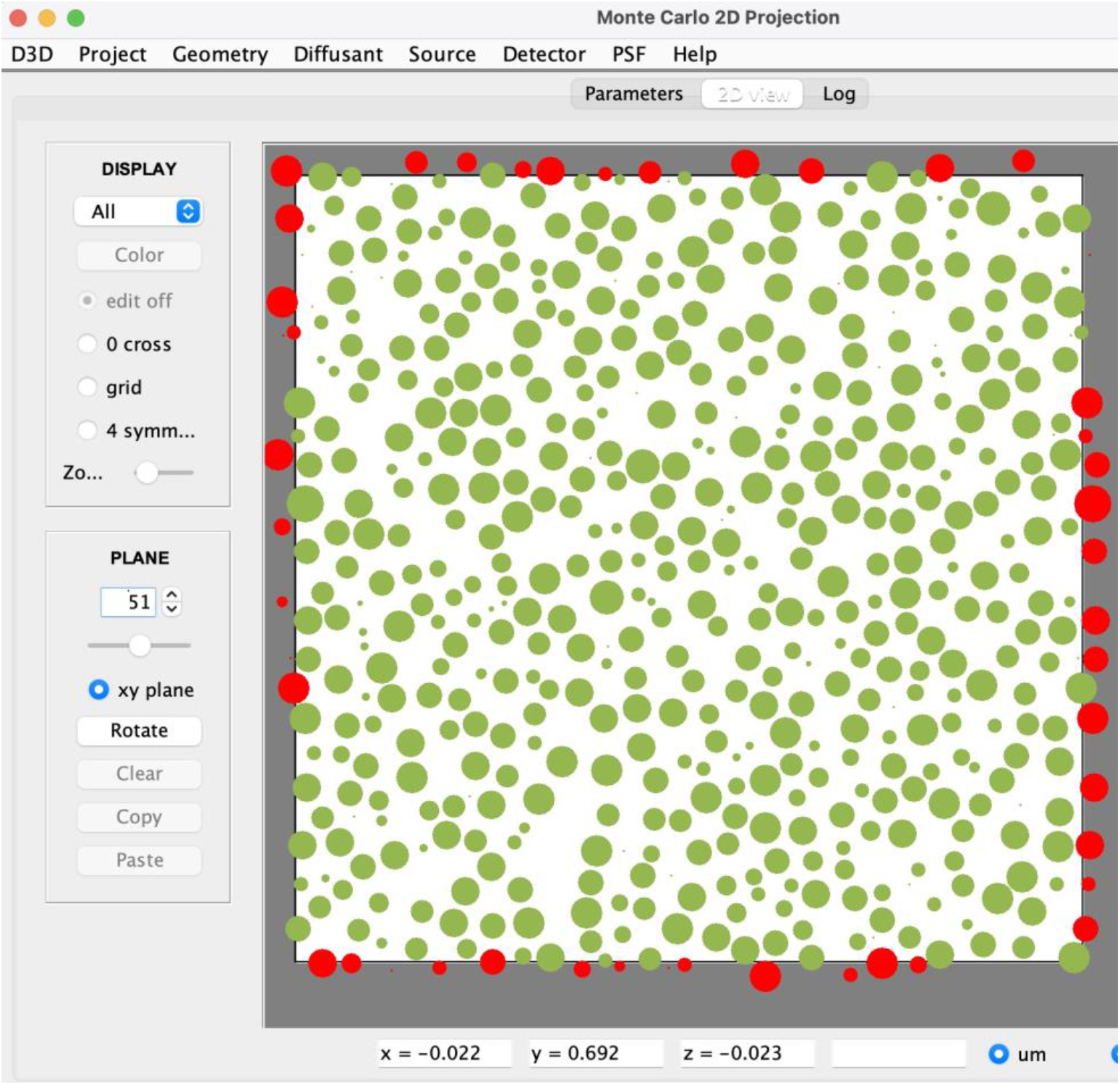
Monte Carlo simulation of a 2D projection of spherical particles. 2D projection of a planar section (T = 0, ϕ = 40°) through the middle of a Monte Carlo simulation (Methods and Materials) in which spherical particles were randomly distributed in a cuboid geometry (26 × 26 × 104 u.d.) where VF = 0.40. F(d) was a Gaussian distribution with normalised mean (Equation 5; μ_D_ ± σ_D_ = 1.00 ± 0.09 u.d.). Red circles denote particles reflected at the x and y borders, i.e. periodic boundary conditions, shown only for display purposes (i.e. they do not contribute to any size or density analysis). The geometry z-dimension was deep enough to accommodate 100 such projections. The simulation was computed via D3D (https://github.com/SilverLabUCL/D3D).

**Figure S3.**
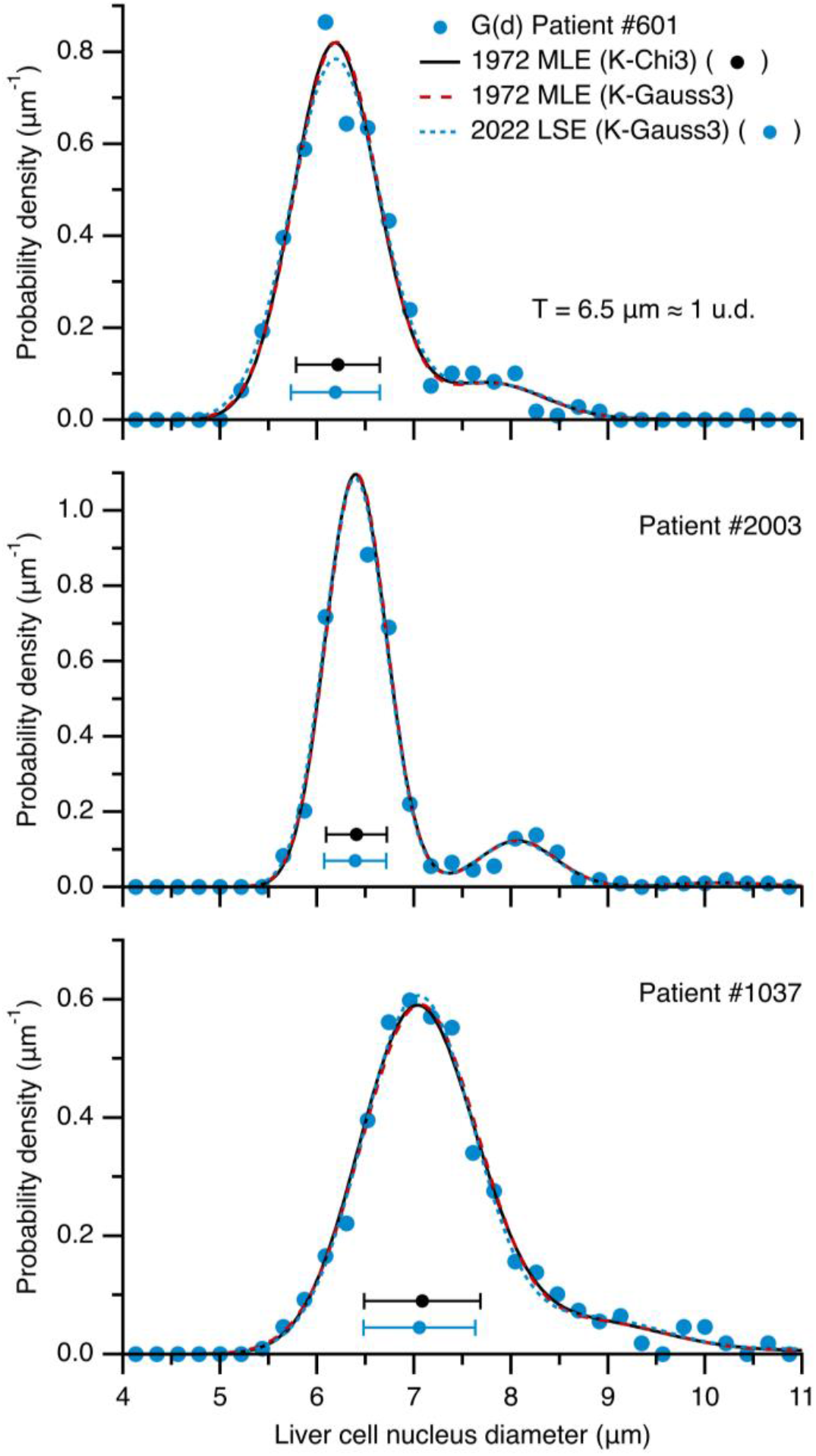
Replication of the original Keiding-model fits to G(d) of liver cell nuclei. G(d) of liver cell nuclei (blue circles) computed from Table 2 of Keiding et al.^49^ for patients 601, 2003 and 1037. Distribution x-axes radii (units of mm) were converted to diameters (units of μm) via the following conversion factor: Θ = 2×1000/2300 μm/mm, where 2300 corrects for magnification. The maximum likelihood estimation (MLE) fits of Keiding et al. (black lines) were computed by plugging the parameters from their Table 2 (*f*, β, ϕ, p1, p2) into a modified version of Equation 1 (NMKeidingChi3) that includes the weighted sum of 3 G(d) as described by Keiding et al. (their Equation 6.1) using a chi distribution for F(d) (K-Chi3; Equation 6) and converting β from units of square radii (mm^2^) to square diameters (μm^2^) by multiplying by Θ^2^. For comparison, MLE fits were recomputed using a Gaussian distribution for F(d) (red lines; K-Gauss3; Equation 5; NMKeidingGauss3) where μ_D_ and σ_D_ were computed from *f* and β. The overlap of these two curves (K-Chi3 vs. K-Gauss3) demonstrates the estimated chi distributions from the MLE fits are approximately Gaussian, which is expected since *f* is large for these fits (104, 208 and 70). As a last comparison, the same modified version of Equation 1 (NMKeidingGauss3) was curve fitted to the 3 G(d) using the LSE routine used in this study (blue dashed lines) resulting in nearly equivalent curves and parameters (Table 4).

**Figure S4.**
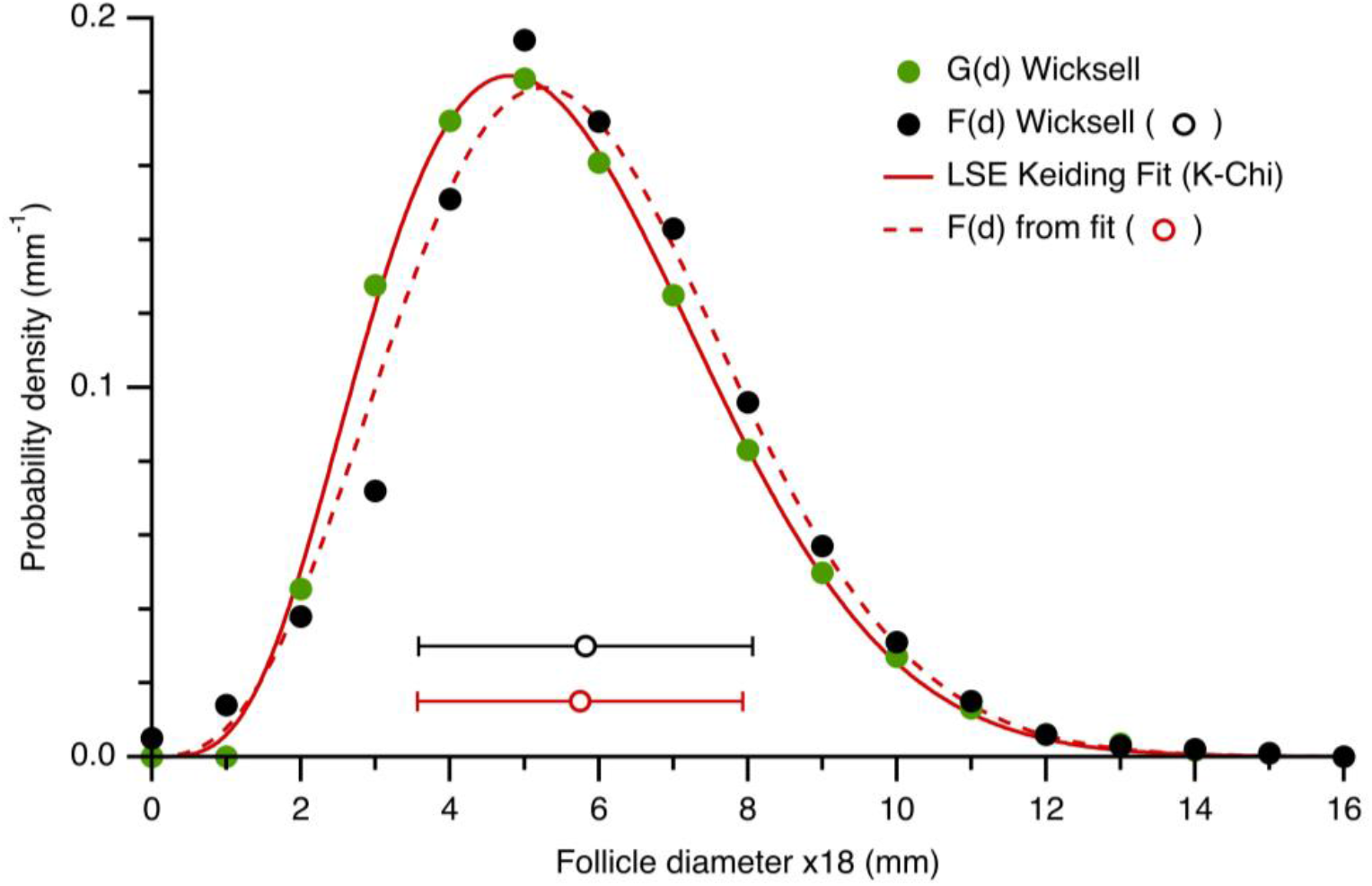
Keiding-model curve fit to G(d) of Wicksell’s corpuscle problem. Curve fit of Equation 1 (red solid line; χ^2^ = 0.0001) to G(d) of Wicksell’s corpuscle problem^16^ (green circles) where F(d) was defined by the chi distribution (K-Chi; Equation 6). F(d) derived from the fit (red dashed line and circle; μ_D_ ± σ_D_ = 5.75 ± 2.18 mm) matches Wicksell’s F(d) computed via a finite-difference unfolding algorithm (black circles; μ_D_ ± σ_D_ = 5.82 ± 2.24 mm). Keiding-model fit *f* = 3.67 ± 0.33, β = 10.31 ± 0.61 mm^2^, ϕ = 25 ± 3°. X-scale of Wicksell includes 18-fold magnification. T = 0.018 mm. Note, assuming a Gaussian function for F(d) (K-Gauss; Equation 5) resulted in a poor fit (χ^2^ = 0.002).

**Figure S5.**
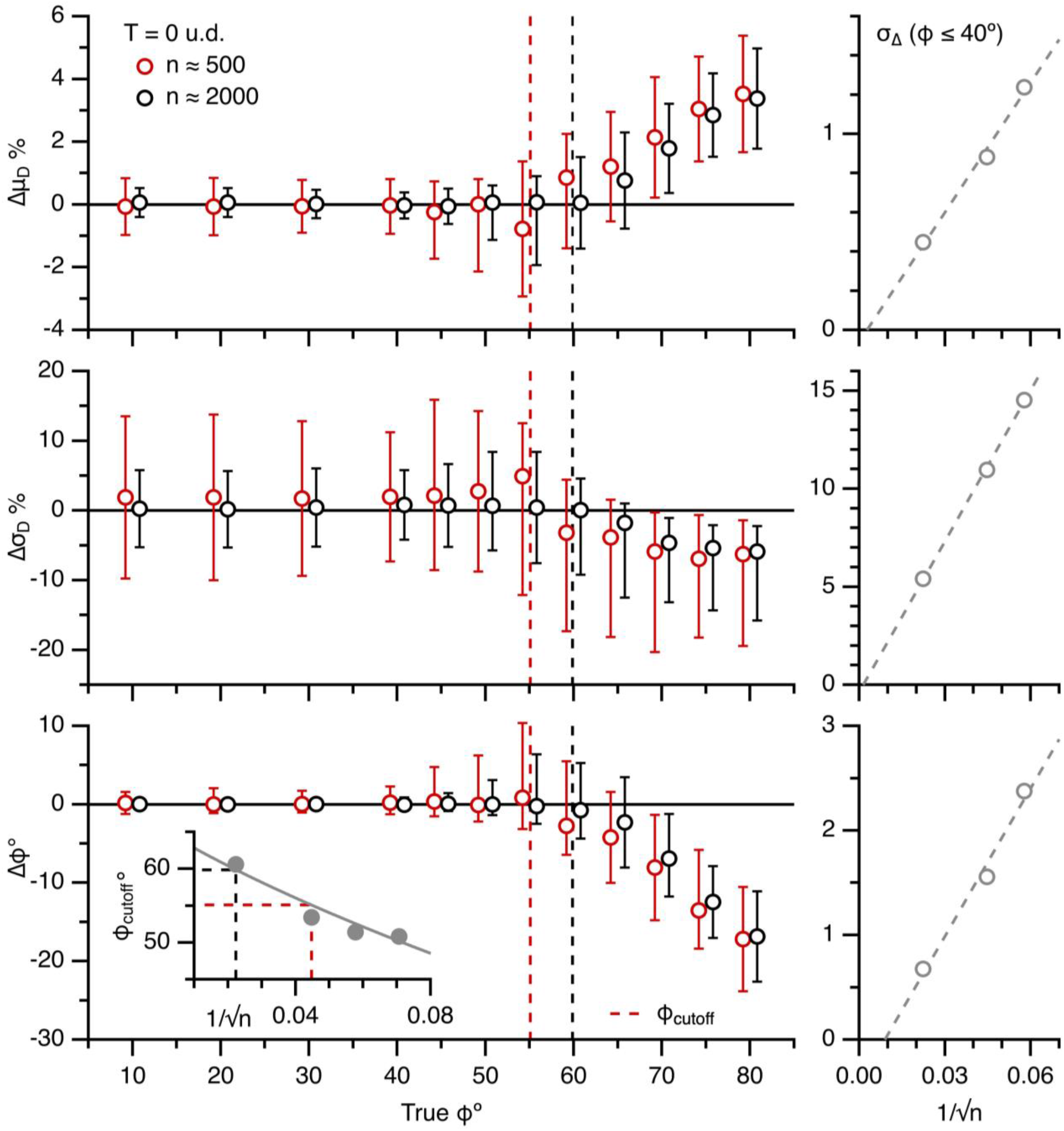
The Keiding model accurately estimates F(d) and ϕ from G(d) for true ϕ < ϕ_cutoff_ (Part II. 500 vs. 2000 simulated diameters). Left column: average Δμ_D_, Δσ_D_ and Δϕ of Keiding-model fits to simulated G(d) computed from ∼500 diameters (red circles; data from Figure 4C) and ∼2000 diameters (black circles) for true ϕ = 10–80°, T = 0 u.d., CV_D_ = 0.09. Red and black dashed lines denote respective ϕ_cutoff_ (∼55 and 60°; Equation 8). Data x-scales shifted ±0.8° to avoid overlap. Bottom inset: ϕ_cutoff_ vs. 1/√n for simulations (gray circles) and Equation 8 (gray line; CV_D_ = 0.09). Right column: 68% confidence interval (σ_Δ_) of Δμ_D_, Δσ_D_ and Δϕ (averaged across true ϕ = 10–40°) vs. 1/√n (n = 300, 500 and 2000 diameters) curve fitted to a linear function (dashed lines).

**Figure S6.**
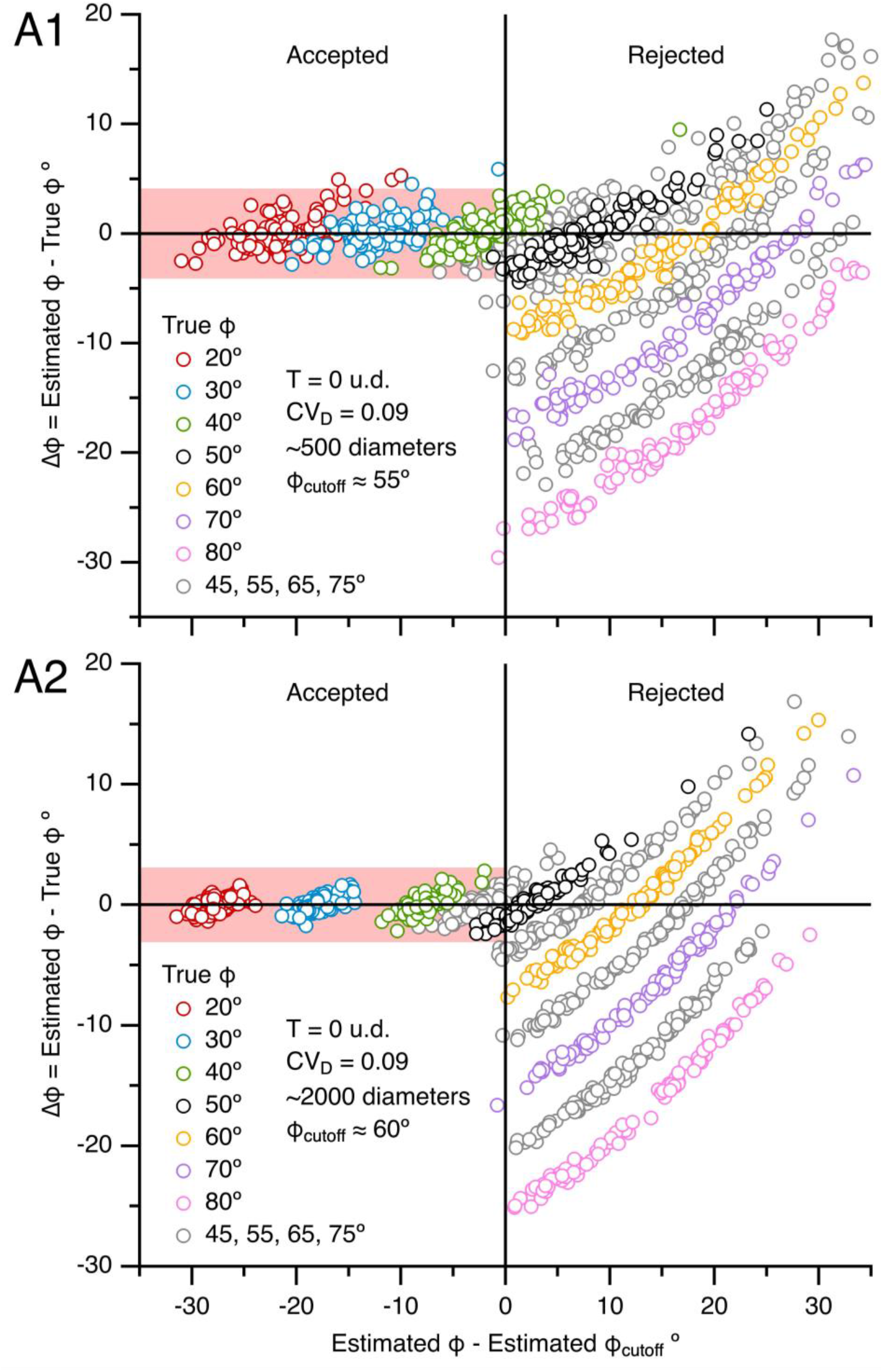
The ϕ-accuracy test using estimated ϕ_cutoff_. Δϕ of Keiding-model fits to simulated G(d) versus the difference between estimated ϕ and estimated ϕ_cutoff_, where estimated ϕ_cutoff_ was computed via Equation 9 using estimated μ_D_ and σ_D_. These plots show that if estimated ϕ < estimated ϕ_cutoff_ (left side of graphs) there is a high probability |Δϕ| < 4° for G(d) computed from ∼500 diameters (A1; red shading; data from Figure 4C), |Δϕ| < 3° for G(d) computed from ∼2000 diameters (A2; data from Figure S5) and |Δϕ| < 5° for G(d) computed from ∼300 diameters (not shown). For the simulations, true ϕ = 20–80°, T = 0 u.d., CV_D_ = 0.09, with 100 simulated G(d) per true ϕ. A few data points are off scale.

**Figure S7.**
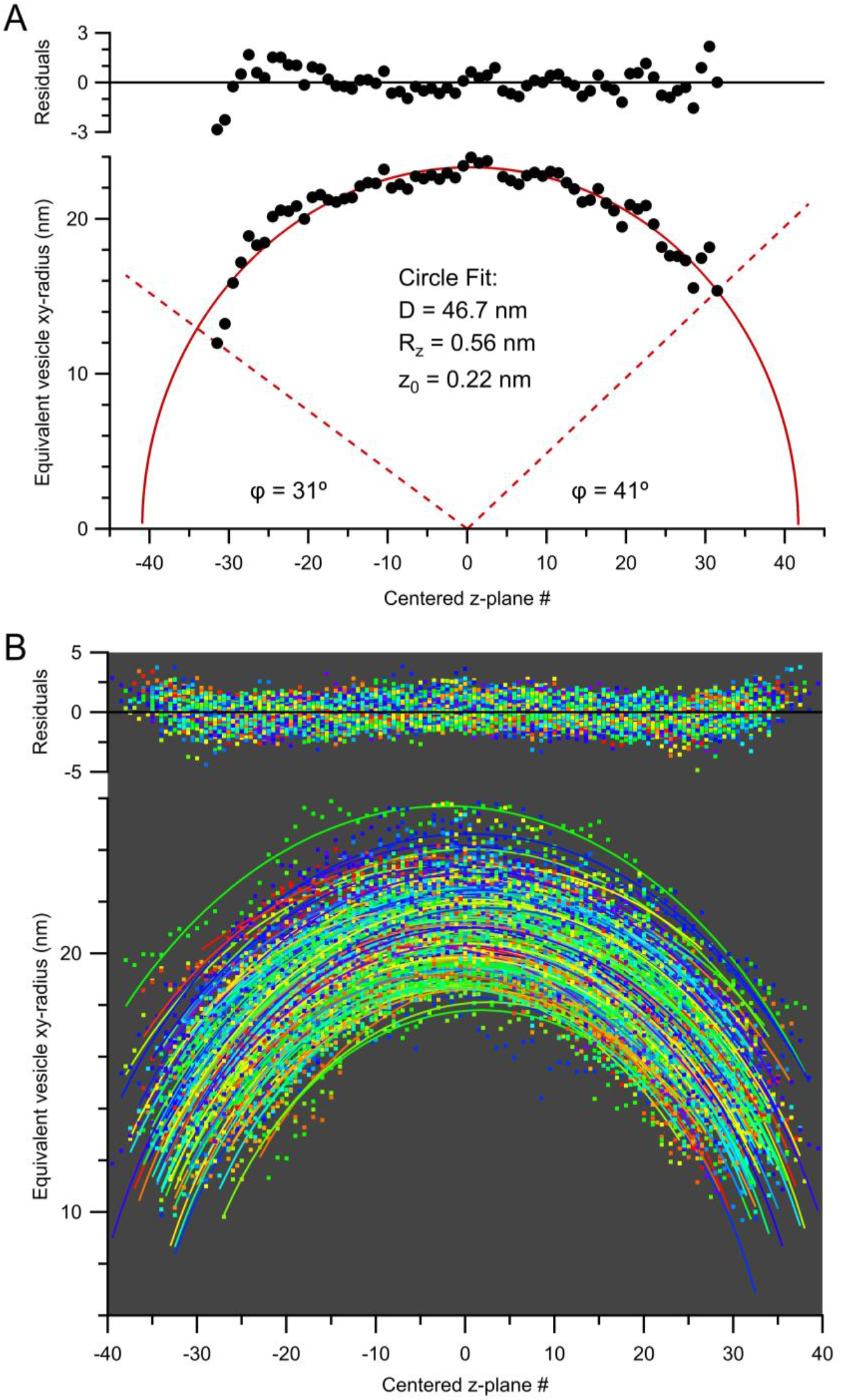
3D analysis of MFT vesicles in an ET z-stack. (A) Example curve fit of a circle (Equation 10; red solid line) to the equivalent xy-radius (½d_area_) vs. z-stack image number (z_#_) relation of a MFT vesicle (black circles) from an ET z-stack where the z-axis was centered at z_#_ = 0. Top graph shows residuals between data and fit. Red dashed lines denote measured ϕ for the vesicle’s north and south poles computed as ϕ = sin^−1^(δ_min_/D), where δ_min_ is the minimum diameter measured at the given pole. (B) Results of a simultaneous curve fit of Equation 10 to the ½d_area_-z_#_ relations of those vesicles whose north and south poles were interior to the z-stack (ET11; Figure 6; n = 163). Fit R_z_ = 0.513 ± 0.001 nm. This R_z_ is 1.4-fold larger than the acquisition R_z_ (0.379 nm) indicating the section thickness after imaging was 74% of the original thickness, which is consistent with previous findings^69^. Repeating the same analyses for the ET10 z-stack gave R_z_ = 0.605 ± 0.002 nm, indicating 63% tissue shrinkage. The fit was computed using Igor Pro’s Global Fit package.

**Figure S8.**
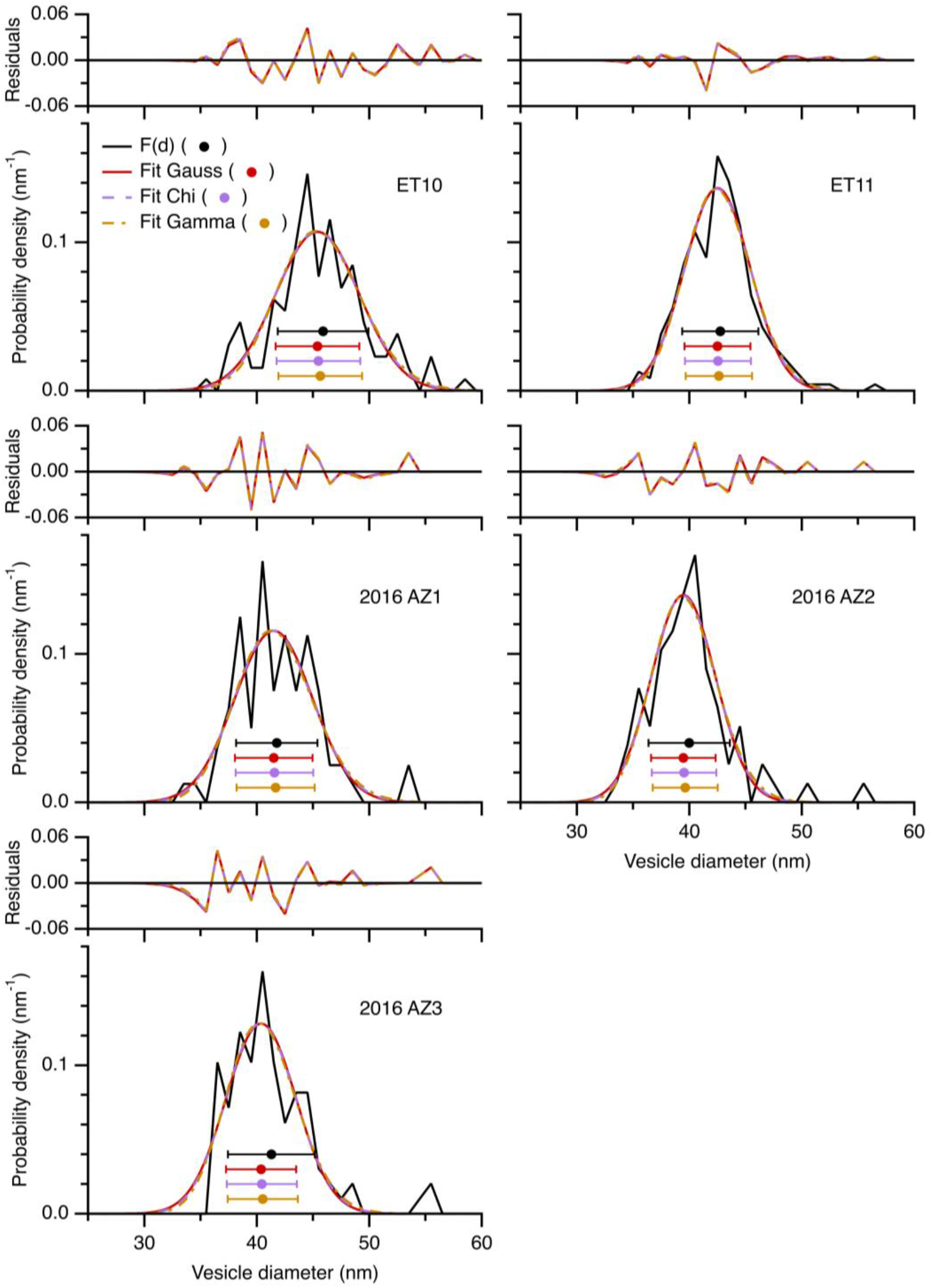
F(d) of cerebellar MFT vesicles is well described by a Gaussian distribution. Comparison of LSE curve fits of a Gaussian distribution (Equation 5; red line), chi distribution (Equation 6; purple line) and gamma distribution (Equation 7; yellow line) to F(d) (black line) in Figures 6D (ET10; n = 130 diameters) and S9D (ET11; n = 234 diameters) and 3 F(d) from a previous study^5^ (AZ1–3; n = 80, 78, 98 diameters) all computed from ET z-stacks of MFT vesicles. The fits overlap and show little difference in residuals, χ^2^ and μ_D_ ± σ_D_ (circles ± error bars). For fits to the gamma distribution, d_0_ was fixed at 10 nm, a value determined by an initial round of fits where d_0_ was allowed to vary.

**Figure S9.**
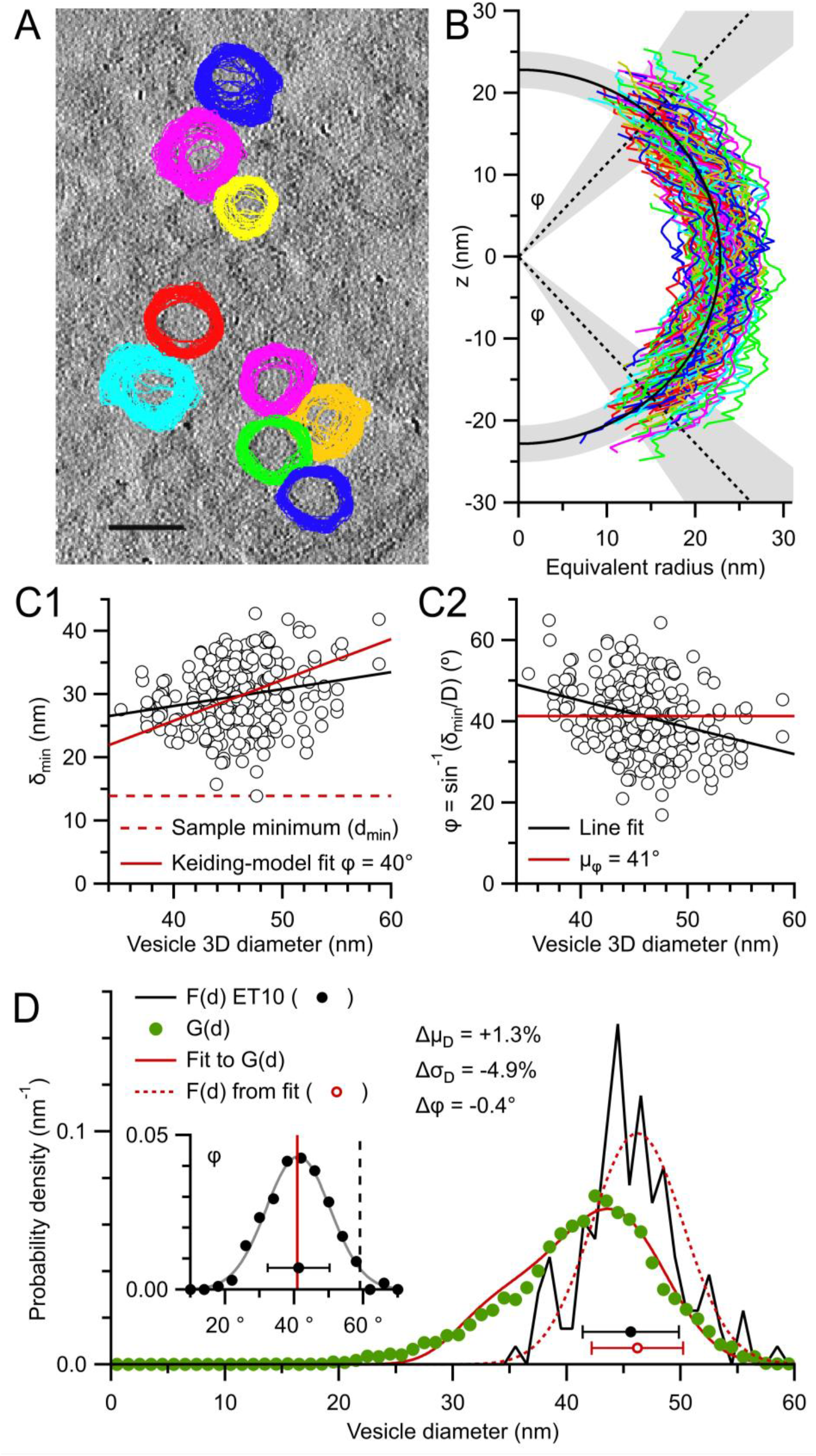
The Keiding model accurately estimates F(d) and ϕ from G(d) for true ϕ < ϕ_cutoff_ (Part III. vesicles ET10). Same as Figure 6 for ET10. (A) One of 291 serial images of a 3D ET reconstruction (ET10) of a cerebellar MFT section 176 nm thick. 141 vesicles, including 24 caps, were tracked and outlined through multiple z-planes and their d_area_ computed as a function of z-plane number. This image shows outlines for 9 representative vesicles, overlaid with outlines from images above and below. Scale bar 50 nm. (B) Black semi-circle and shading denote F(d) = μ_D_ ± σ_D_ = 45.6 ± 4.2 nm for all measured 3D diameters (D; n = 130). Black dotted lines and shading denote measured ϕ: μ_ϕ_ ± σ_ϕ_ = 41.3 ± 8.9° (n = 247, measures from both poles). (C1) δ_min_ vs. D (circles; n = 247) with line fit (black line; χ^2^ = 6398, PCC = 0.2, r^2^ = 0.05) and Keiding-model fit (red solid line; fit ϕ = 40.2 ± 0.4°; χ^2^ = 7019, PCC = 0.2, r^2^ = 0.3). (C2) ϕ vs. D for data in (C1) with line fit (black line; PCC = -0.3, r^2^ = 0.1) and μ_ϕ_ = 41.3° (red solid line). (D) Measured F(d) (black line and circle) vs. G(d) (green circles; n = 7083 outlines). A curve fit of Equation 1 to G(d) (red solid line; μ_D_ = 46.2 ± 0.1 nm, σ_D_ = 4.0 ± 0.1 nm, ϕ = 40.9 ± 0.6°; T fixed to 0 nm) resulted in estimated F(d) (red dotted line and circle) nearly the same as measured F(d). Inset: probability density of measured ϕ in (C2) with Gaussian fit (gray line; Equation 5; μ_ϕ_ ± σ_ϕ_ = 41.2 ± 9.3°), fit ϕ (red line) and ϕ_cutoff_ (black dashed line; ∼59°; Equation 8).

**Figure S10.**
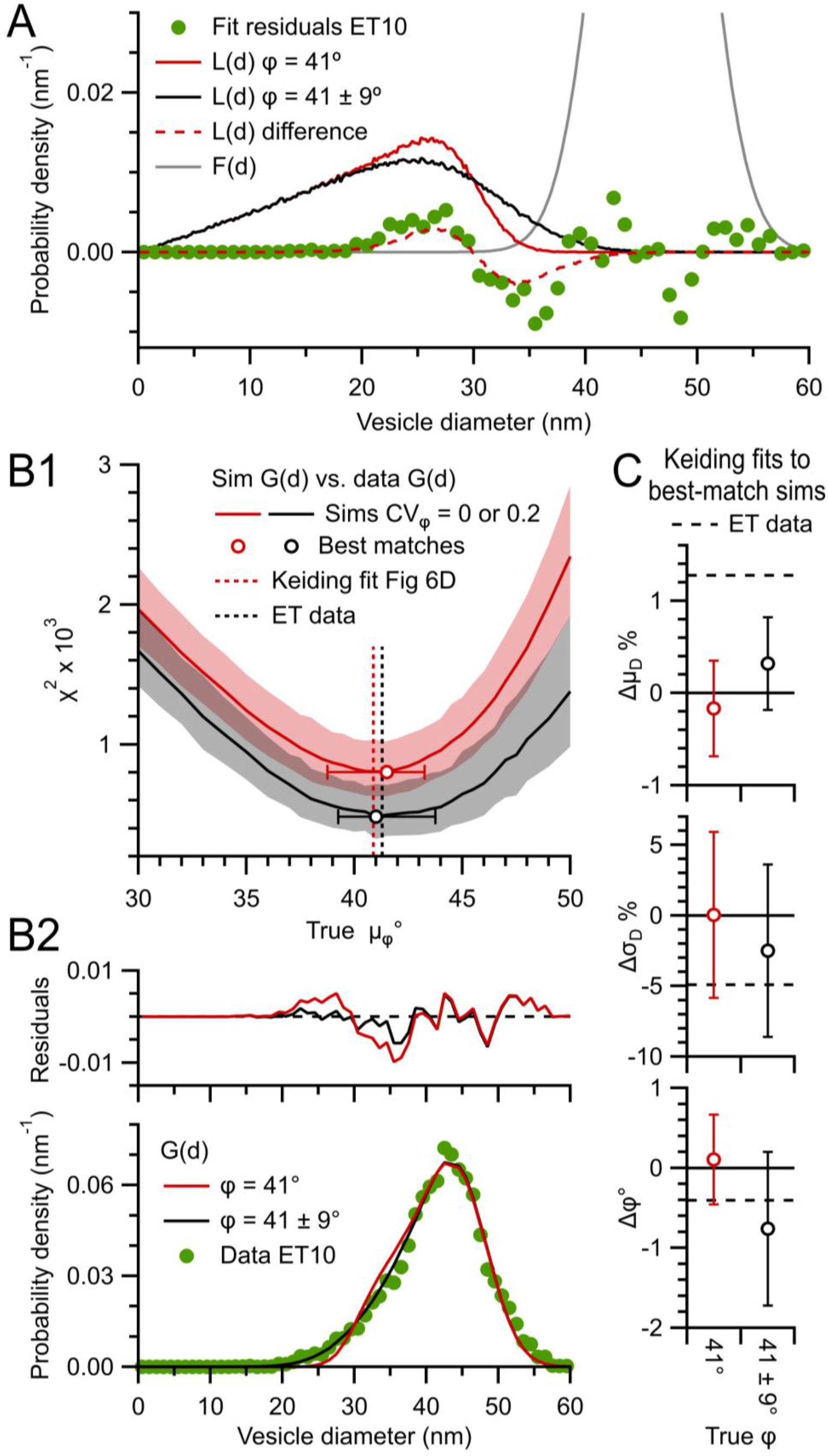
The Keiding-model fixed-ϕ assumption introduces only small errors when estimating F(d) and ϕ from G(d) (Part I: z-stack simulations). (A) Distribution of lost caps, L(d), for the Keiding-model fit in Figure S9D where ϕ = 41° (red solid line; ET10; see Figure 3F) and Gaussian distribution ϕ = 41 ± 9° (black solid line). The difference between these two distributions (red dashed line) matches the residuals of the Keiding-model fit to G(d) in Figure S9D. Hence, the deviation of the Keiding-model fit from G(d) at 20–40 nm can be explained by the difference between a model that assumes a fixed ϕ (the Keiding model) and a Gaussian ϕ. L(d) was computed via simulations. F(d) is shown for comparison (gray line) and goes off scale. (B1) χ^2^ comparison of experimental G(d) in Figure S9D to G(d) computed from a Monte Carlo model that assumes either a fixed ϕ as in the Keiding model (CV_ϕ_ = 0; red solid line denotes 50% probability and shading denotes 16–84% probability computed from 100 repetitions per μ_ϕ_) or a Gaussian ϕ that matches the experimental data in Figure S9D (CV_ϕ_ = 0.2; black solid line and shading). The Gaussian-ϕ model had significantly smaller χ^2^ than the fixed-ϕ model. To simulate the z-stack data in Figure 6, F(d) of the simulated particles was matched to the measured F(d) (μ_D_ ± σ_D_ = 46 ± 4 nm) and 2D diameters were computed from particle projections within an xy-plane (0.28 × 0.28 μm; T = 0 nm) that was z-shifted 290 times in 0.6 nm steps. The number of analysed particles (∼140) and 2D diameters (∼6600) was similar that of the experimental data for μ_ϕ_ = 41°. μ_ϕ_ was the only free parameter and was varied between 37–48° in steps of 0.5°, or 1° outside this region. Circles denote best-match μ_ϕ_ of the fixed-ϕ model (41.5°) and Gaussian-ϕ model (41.0°), as described below. Particle VF = 0.45. Red vertical dashed line denotes ϕ of the Keiding-model fit to experimental G(d). Black vertical dashed line denotes measured μ_ϕ_. ϕ_cutoff_ ≈ 60° (Equation 8). (B2) Experimental G(d) (bottom; green circles) compared to best-match simulated G(d) in (A1), where simulated G(d) are the average for 100 repetitions for μ_ϕ_ = 41° (red and black lines). Differences between experimental and simulated G(d) (top) shows the Gaussian-ϕ model is a better match to experimental G(d). (C) Average Δμ_D_, Δσ_D_ and Δϕ of curve fits of Equation 1 to the 100 G(d) of the best-match simulations in (A1), computed with respect to ‘measured’ μ_D_, σ_D_ and μ_ϕ_ of the particles in the projection. Black dashed lines denote Δμ_D_, Δσ_D_ and Δϕ of the experimental data in Figure S9D. Errors of the Gaussian-ϕ model match the experimental errors better than those of the fix-ϕ model. To find the best-match μ_ϕ_ for the given experimental G(d), the sum of squared differences (χ^2^) was computed between the experimental G(d) and simulated G(d). Cumulative distribution functions (CDFs) of χ^2^ were computed from the 100 simulated G(d) at a given μ_ϕ_. From the CDFs, χ^2^ values with 50% probability (χ^2^-50%) were computed as a function of μ_ϕ_, as well as χ^2^ values with 16 and 84% probabilities representing ±σ. The μ_ϕ_ with smallest χ^2^-50% was deemed the best match. Confidence intervals above and below the best-match μ_ϕ_ denote the range of μ_ϕ_ whose χ^2^ are not significantly different to that of the best-match μ_ϕ_, computed via a KS test (p > 0.05).

**Figure S11.**
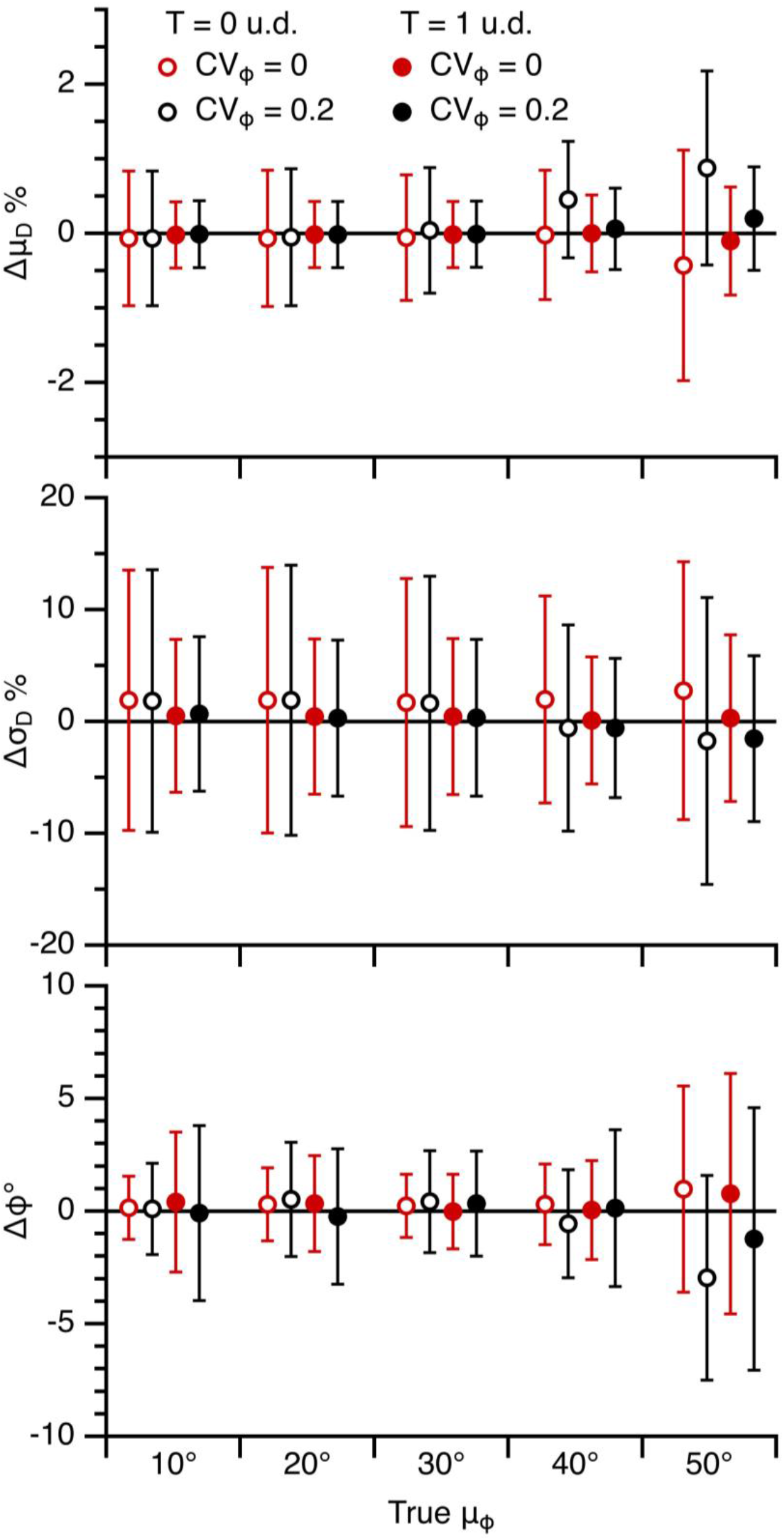
The Keiding-model fixed-ϕ assumption introduces only small errors when estimating F(d) and ϕ from G(d) (Part II: 2D projection simulations). Average Δμ_D_, Δσ_D_ and Δϕ of curve fits of Equation 1 to 100 G(d) computed from simulations as in Figures 3 and 4 for ∼500 particles with a fixed or Gaussian ϕ for μ_ϕ_ = 10–50° (CV_ϕ_ = 0 or 0.2; red vs. black) and T = 0 and 1 u.d. (open and closed circles). Fixed-ϕ simulation data is from Figure 4C. ϕ_cutoff_ ≈ 55° (Equation 8).

**Figure S12.**
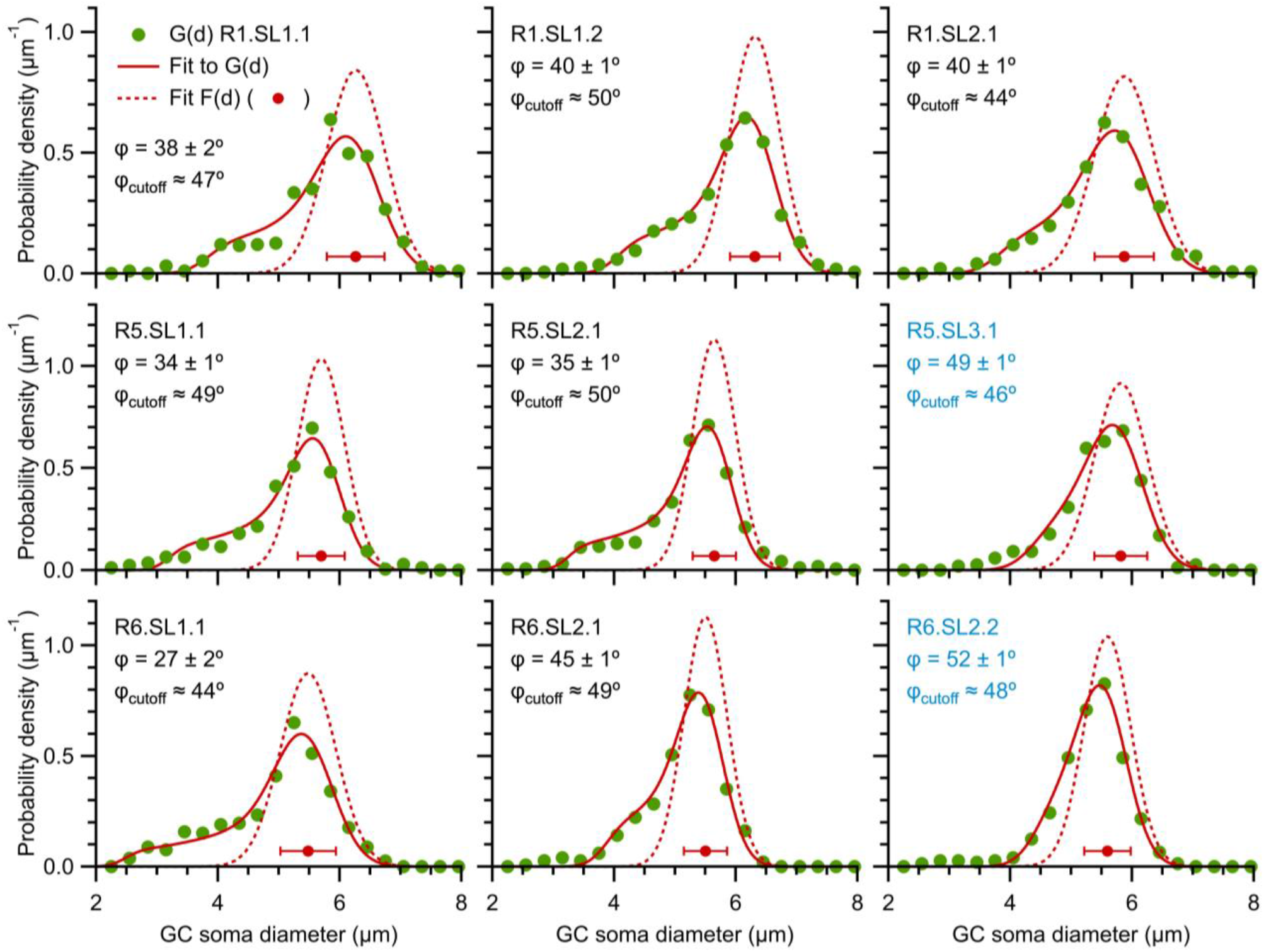
Keiding-model fits to G(d) of cerebellar GC somata. G(d) (green circles) computed from outlines of GC somata (n = 494–638; 0.3 μm bins) measured from confocal images of the GC layer as in Figure 2A1 and A2, where each row is for a different rat (R1, R5, R6) and each G(d) is computed from 1–3 images of a confocal z-stack. There were 2–3 tissue sections per rat (SL1, SL2, SL3). Each G(d) was curve fitted to Equation 1 (red solid lines) resulting in estimates for F(d) (red dotted lines and circles) and ϕ. Estimated ϕ_cutoff_ was computed via Equation 9. For all but two fits, estimated ϕ < estimated ϕ_cutoff_. Comparisons of the diameter distributions within rats showed significant differences (KS test; p < 0.05), even after alignment on their mean μ_D_.

**Figure S13.**
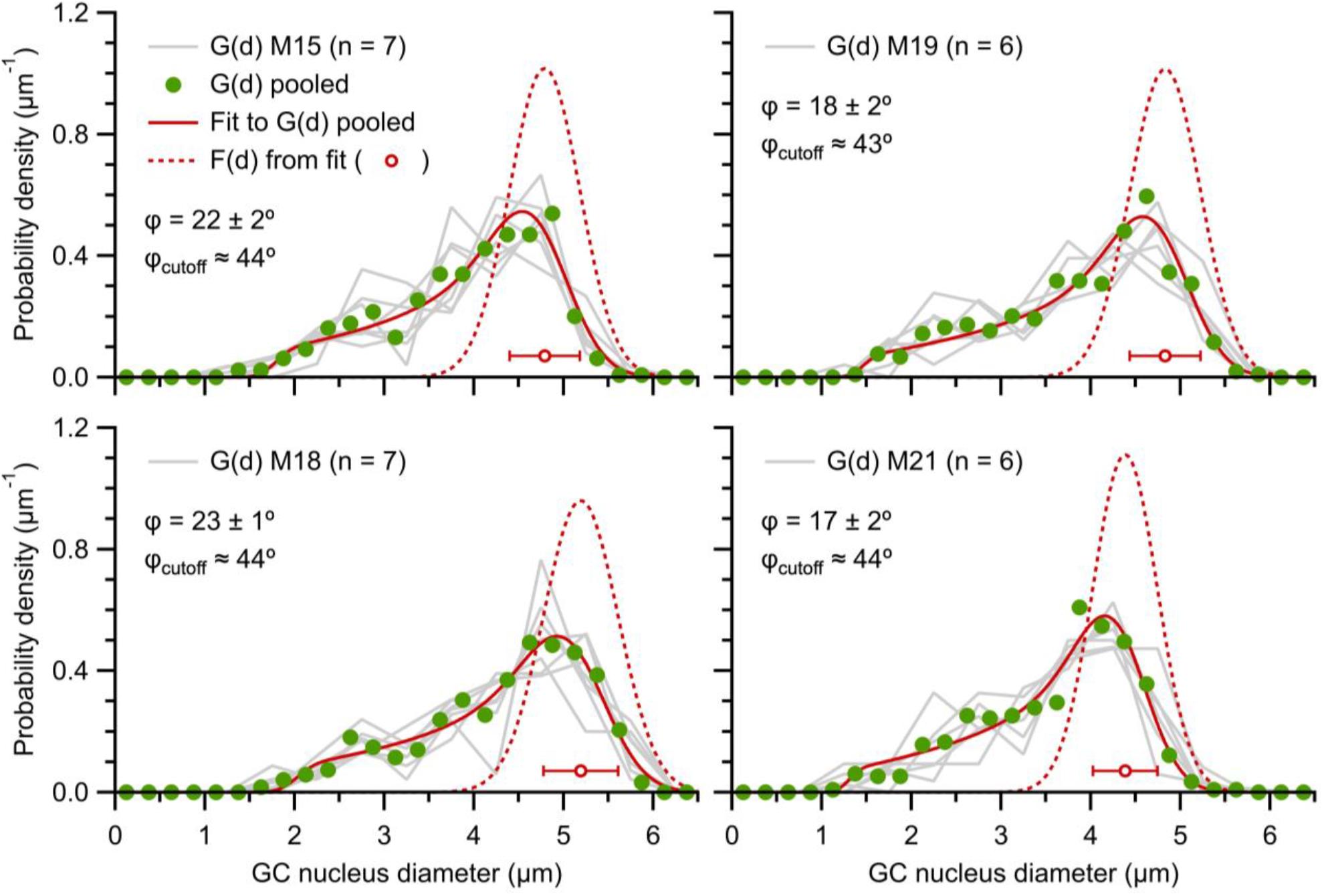
Keiding-model fits to G(d) of cerebellar GC nuclei. G(d) (gray lines) computed from outlines of GC nuclei (n = 35–158 diameters; 0.5 μm bins) measured from TEM images of the GC layer as in Figure 2B1 and B2, where each plot is for a different mouse (M15, M18, M19, M21). G(d) within mice (n = 6 or 7) were not significantly different and therefore pooled (green circles; n = 416–519 diameters; 0.25 μm bins). The resulting 4 pooled G(d) were curve fitted to Equation 1 (red solid lines) resulting in estimates for F(d) (red dotted lines and circles) and ϕ. Estimated ϕ_cutoff_ was computed via Equation 9. Image IDs: M15 = M15.L3.11, 12, 15, 22, 25, M15.L4.04, 08 (n=7); M18 = M18.N2.02, 05, 08, 51, M18.N3.17, M18.N4.10, 15 (n = 7); M19| = M19.O2.06, 12, 13, 20, 38, 44 (n = 6); M21 = M21.P5.16, 19, 33, 48, 52, 59 (n = 6).

**Figure S14.**
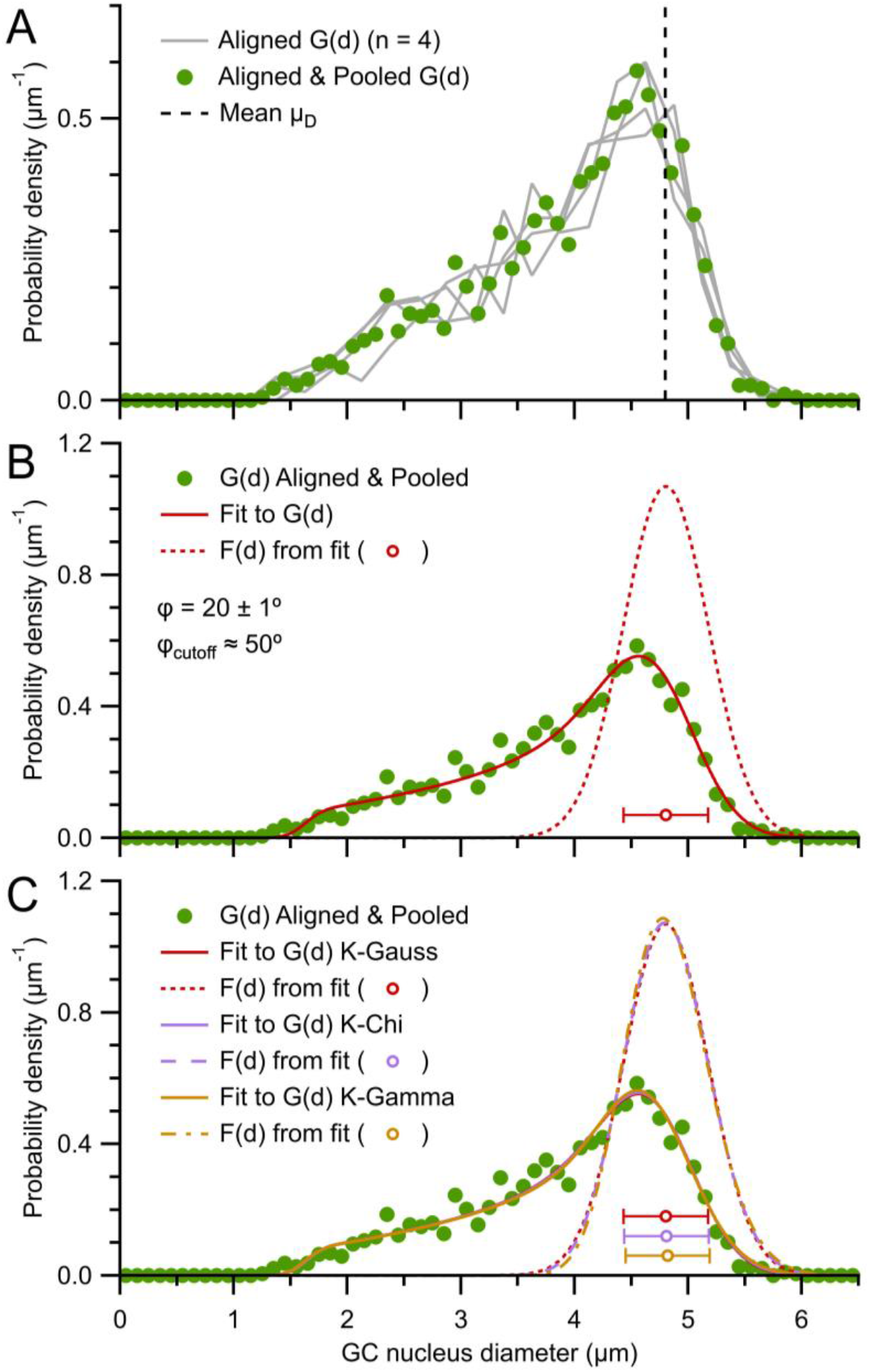
Analysis of aligned G(d) of cerebellar GC nuclei. (A) Pooled G(d) for each mouse in Figure S13 (gray lines) aligned at their mean μ_D_ (4.80 μm; black dashed line). After alignment, none of the 4 distributions were significantly different (KS test; p > 0.16) and were therefore pooled into a single G(d) (green circles; n = 1882 diameters; 0.1 μm bins). (B) Aligned and pooled G(d) from (A) curve fitted to Equation 1 (red solid line; μ_D_ = 4.80 ± 0.01 μm, σ_D_ = 0.37 ± 0.01 μm, ϕ = 20 ± 1°; estimated ϕ_cutoff_ = 50°) with estimated F(d) (red dotted line and circle). (C) The Keiding-model fit in (B), which assumed a Gaussian function for F(d) (K-Gauss; Equation 5), compared to fits that assumed a chi distribution for F(d) (K-Chi; Equation 6; purple line; *f* = 84.0 ± 6.3, β = 0.28 ± 0.02 μm^2^, ϕ = 20 ± 1°; μ_D_ ± σ_D_ = 4.81 ± 0.37 μm) and gamma distribution for F(d) (K-Gamma; Equation 7; orange line; *f* = 80.9 ± 6.2, β = 0.041 ± 0.003 μm, ϕ = 20 ± 1°, d_0_ fixed at 1.5 μm; μ_D_ ± σ_D_ = 4.82 ± 0.37 μm). Comparison of F(d) from the fits show overlapping distributions (circles and error bars).

**Figure S15.**
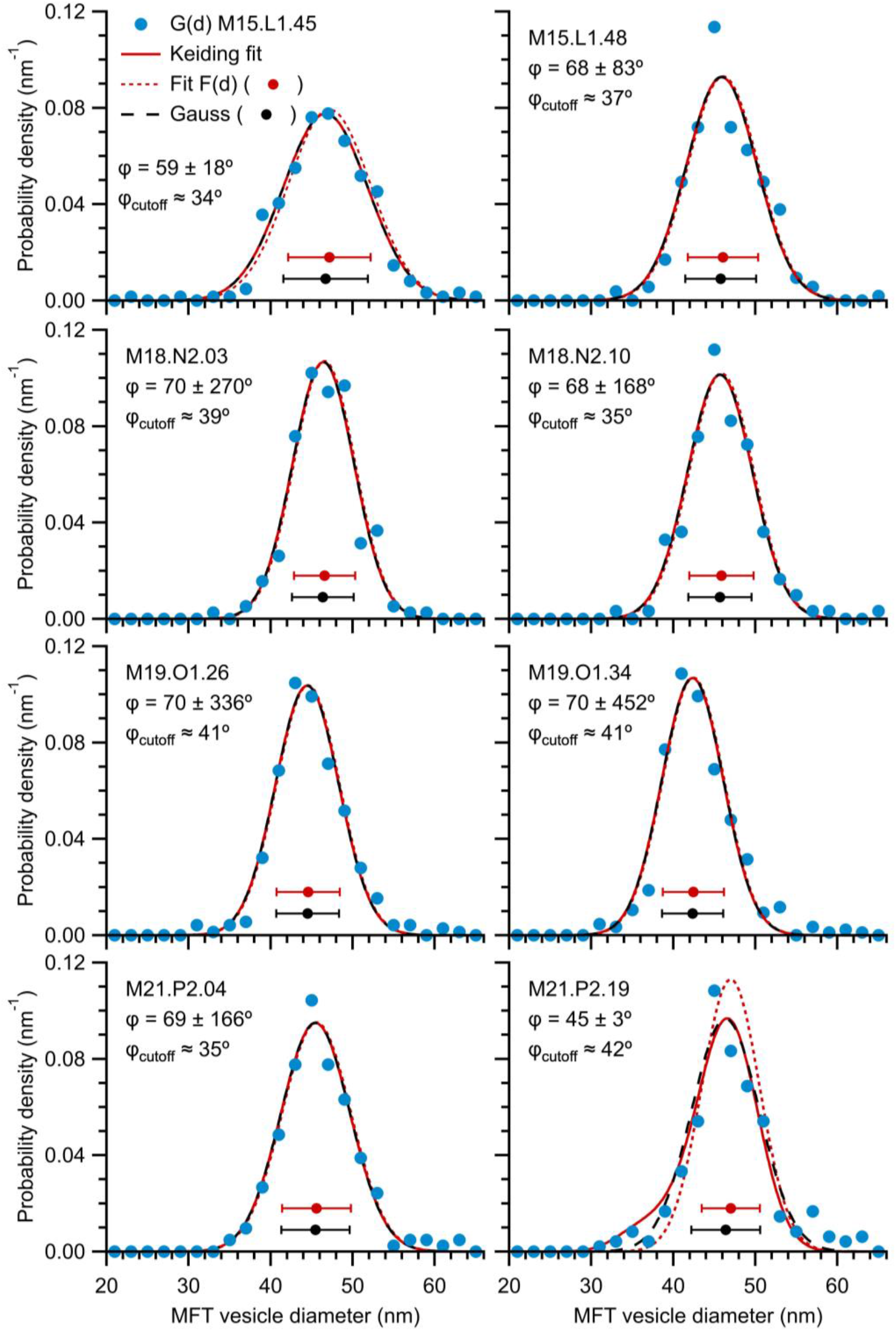
Keiding-model fits to G(d) of cerebellar MFT vesicles. G(d) (blue circles) computed from outlines of MFT vesicles (n = 152–428; 2 nm bins) measured from TEM images of the GC layer as in Figure 2C1 and C2, where each row is for a different mouse and each column is for a different MFT. Each G(d) was curve fitted to Equation 1 (red solid lines) resulting in estimates of F(d) (red dotted lines and circles). For all fits, ϕ has a large error and is greater than estimated ϕ_cutoff_ (Equation 9) indicating ϕ is indeterminable and G(d) ≈ F(d) (Figure 4C). Gaussian fits to the same G(d) (Equation 5; black dashed lines and circles) overlap the Keiding-model fits and estimated F(d).

**Figure S16.**
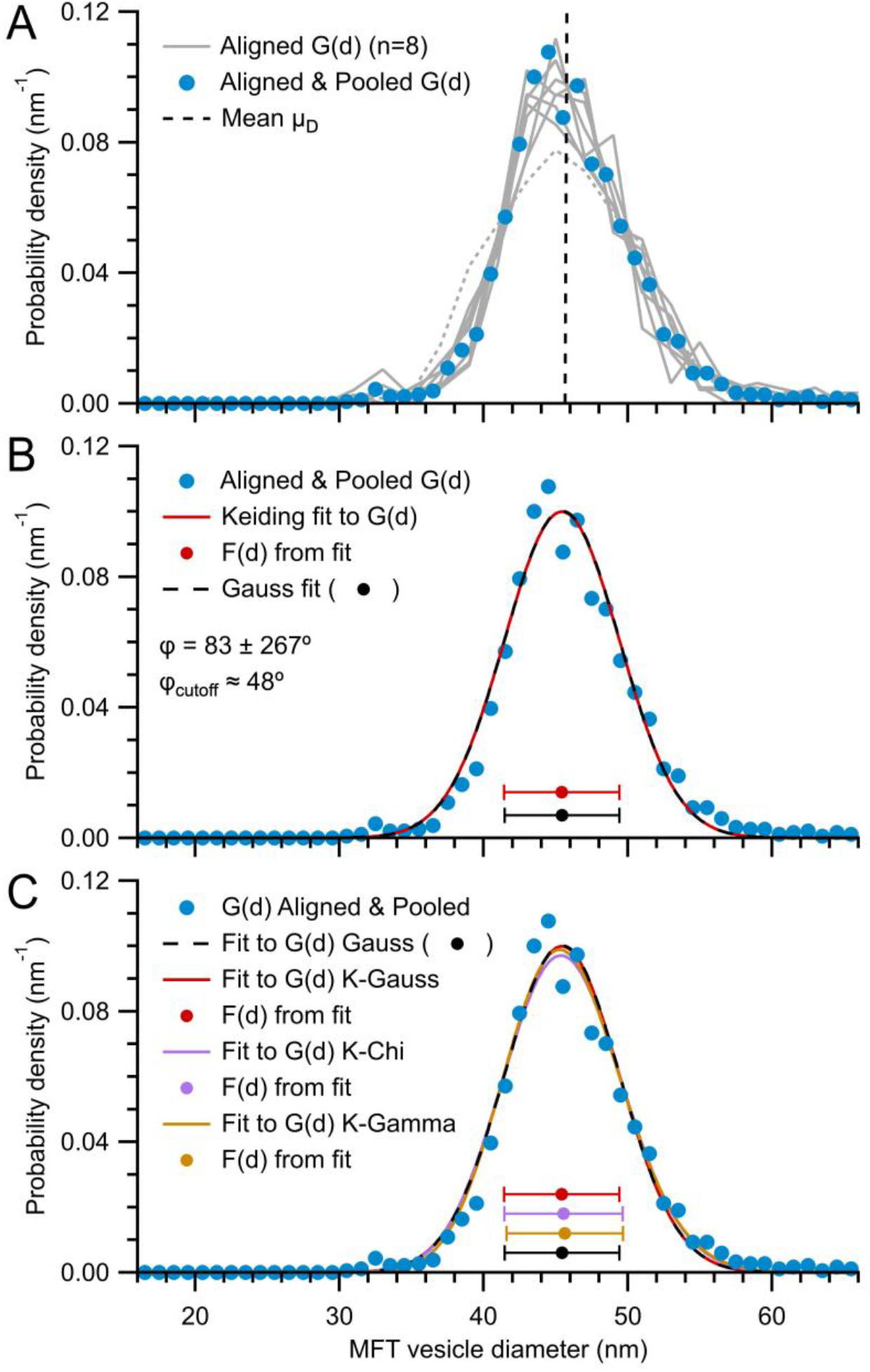
Analysis of aligned G(d) of cerebellar MFT vesicles. (A) G(d) of the 8 MFTs in Figure S15 (gray lines) aligned at their mean μ_D_ (45.7 nm; black dashed line). After alignment, only 2 of the 28 pairings of the distributions were significantly different (KS test; p < 0.02) both of which included the distribution with largest σ_D_ (gray dotted line; M15.L1.45). The aligned G(d) that were not significantly different (n = 7) were pooled into a single G(d) (blue circles; n = 1839 diameters; 1 nm bins). (B) Aligned and pooled G(d) from (A) (blue circles) curve fitted to Equation 1 (red solid line; μ_D_ = 45.4 ± 0.1 nm, σ_D_ = 4.0 ± 0.2 nm, ϕ = 83 ± 267°) with estimated F(d) (red circle and error bars) and ϕ_cutoff_. As in Figure S15, ϕ is indeterminable and G(d) ≈ F(d). A Gaussian fit to G(d) (Equation 5; black dashed line and circle; μ_D_ = 45.4 ± 0.1 nm, σ_D_ = 4.0 ± 0.1 nm) overlaps the Keiding-model fit and estimated F(d). (C) The Keiding-model fit in (B), which assumed a Gaussian function for F(d) (K-Gauss; Equation 5), compared to fits that assumed a chi distribution for F(d) (K-Chi; Equation 6; purple line; *f* = 61.8 ± 1.9, β = 33.9 ± 1.0 nm^2^, ϕ = 90 ± 49°; μ_D_ ± σ_D_ = 45.5 ± 4.1 nm) and gamma distribution for F(d) (K-Gamma; Equation 7; orange line; *f* = 127.0 ± 3.3, β = 0.36 ± 0.01 nm, ϕ = 89 ± 137°, d_0_ fixed at 0 nm; μ_D_ ± σ_D_ = 45.6 ± 4.0 nm). Comparison of F(d) from the fits show overlapping distributions (circles and error bars).

**Figure S17.**
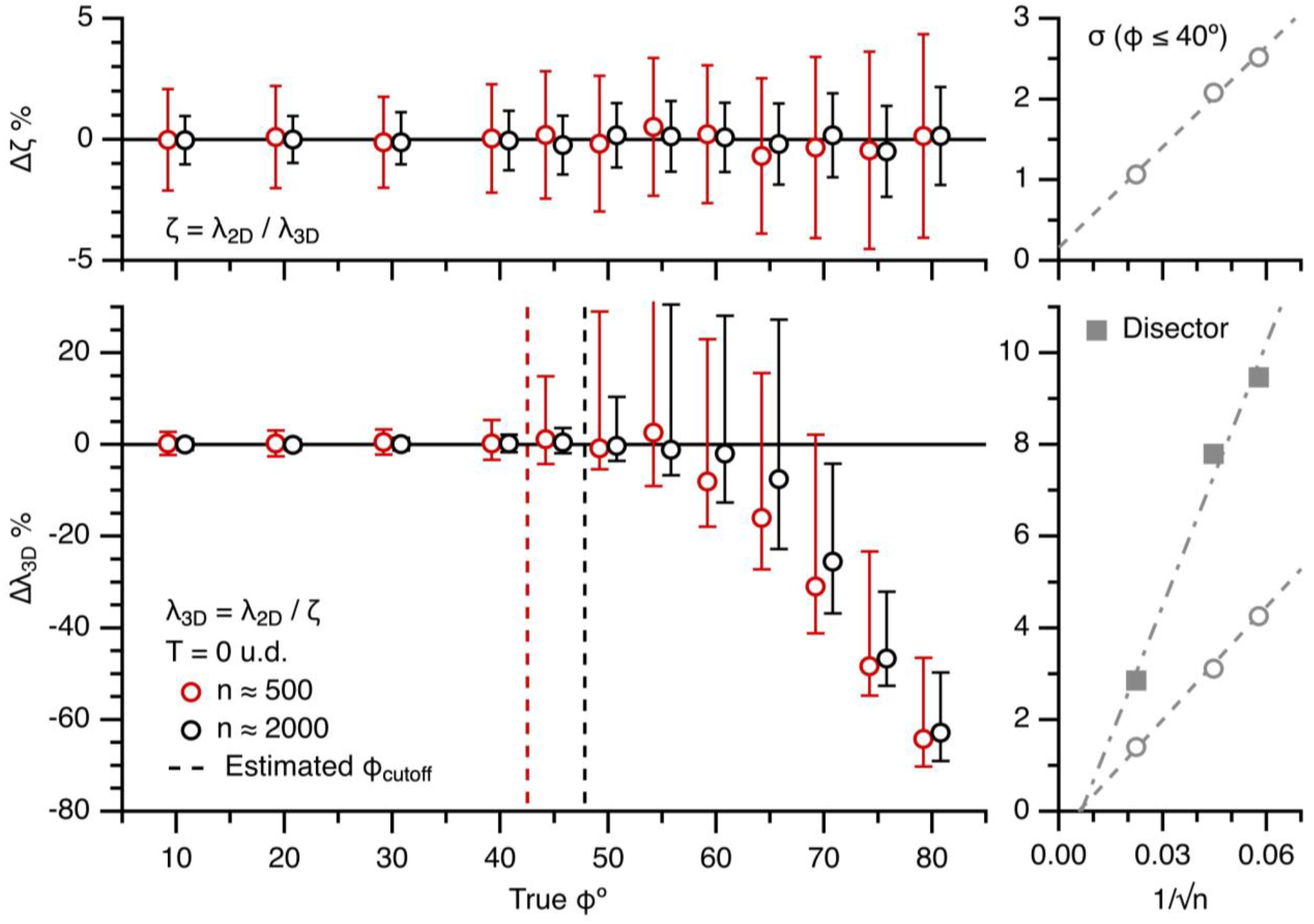
The Keiding model accurately estimates λ_3D_ from λ_2D_ for true ϕ < estimated ϕ_cutoff_ (Part II. 500 vs. 2000 simulated diameters). Left column: average Δζ (top) and Δλ_3D_ (bottom) vs. true ϕ for the simulations in Figure S5, computed from ∼500 and ∼2000 diameters (red and black circles) for T = 0 u.d. as described in Figure 10. Dashed lines denote estimated ϕ_cutoff_ (∼43 and 48°). Data x-scales shifted ±0.8° to avoid overlap. Right column: 68% confidence interval (σ_Δ_) of Δζ and Δλ_3D_ (averaged across true ϕ = 10–40°) vs. 1/√n (n = 300, 500 and 2000 diameters) curve fitted to a linear function (dashed lines). Data from disector simulations with no added bias is shown for comparison (solid squares; see Figure 12, ϕ_bias_ = 0°).

**Figure S18.**
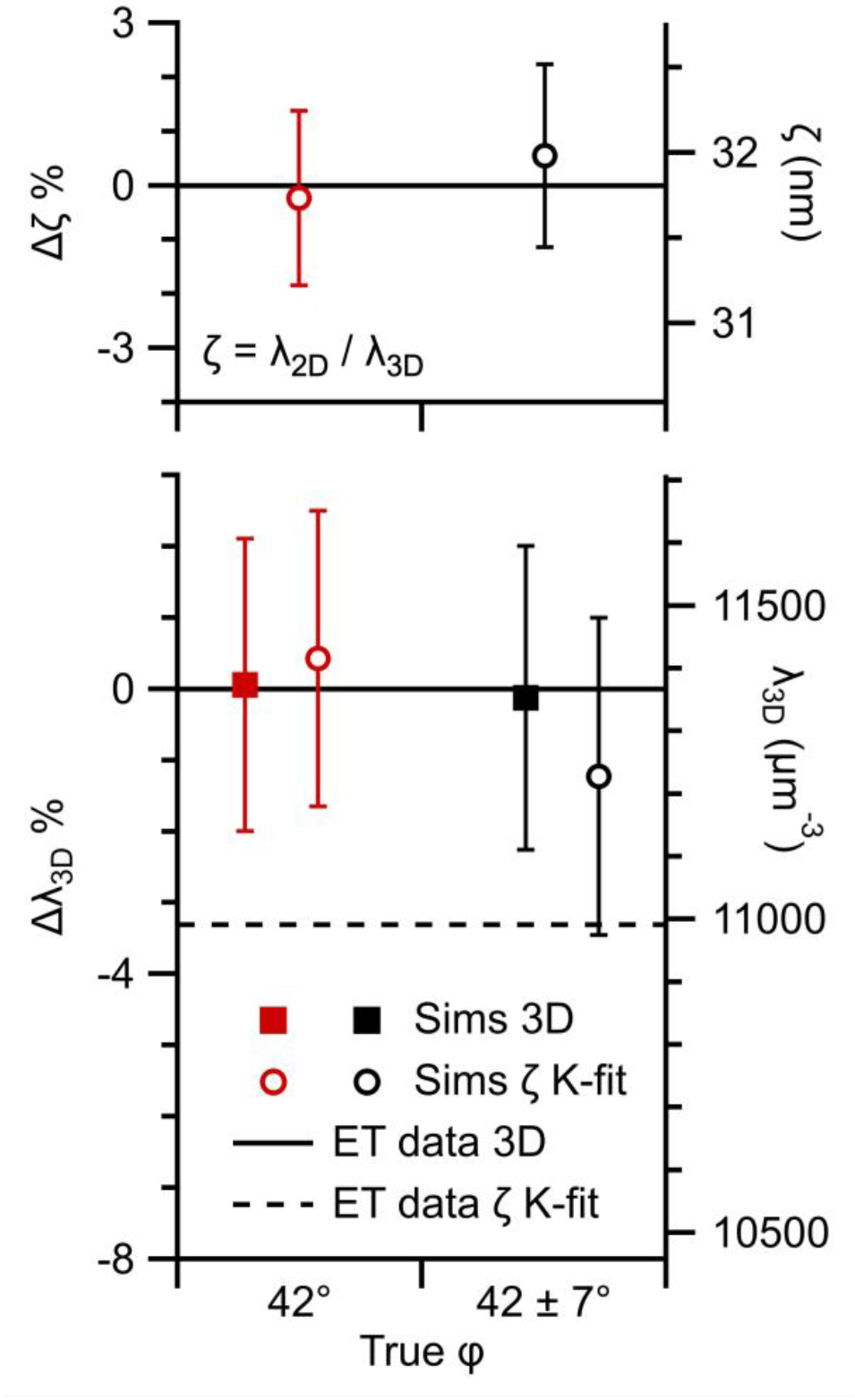
The Keiding-model fixed-ϕ assumption introduces only small errors when estimating λ_3D_ from λ_2D_ (Part I. z-stack simulations). Average Δζ (top) and Δλ_3D_ (bottom) for simulations of the z-stack analysis in Figure 11A2 (ET11) where lost caps are defined by a fixed-ϕ or Gaussian-ϕ model (CV_ϕ_ = 0 or 0.16; red vs. black symbols) and the scanning xy-plane (T = 0 nm) was z-shifted 261 times in 0.5 nm steps resulting in ∼10,500 2D diameters. Δζ and Δλ_3D_ were computed as described in Figure 10 (open circles; true ϕ = μ_ϕ_). For the volumetric analysis, estimated λ_3D_ = N_3D_/(Area_xy_·ζ) where ζ was computed via μ_D_ and μ_ϕ_ of those vesicles sampled by the z-stack (3D; N_3D_ ≈ 200; closed squares). Simulation true λ_3D_ = 10,825 μm^−3^ (VF = 0.45), μ_D_ = 42.7 nm, μ_ϕ_ = 42.0°, Area_xy_ = 0.117 μm^2^. Equivalent ζ and λ_3D_ for the ET z-stack analysis of MFT vesicles (ET11) is shown for comparison (black solid and dashed lines) which are consistent with the Gaussian-ϕ simulations. Estimated ϕ_cutoff_ ≈ 51° (Equation 9).

**Figure S19.**
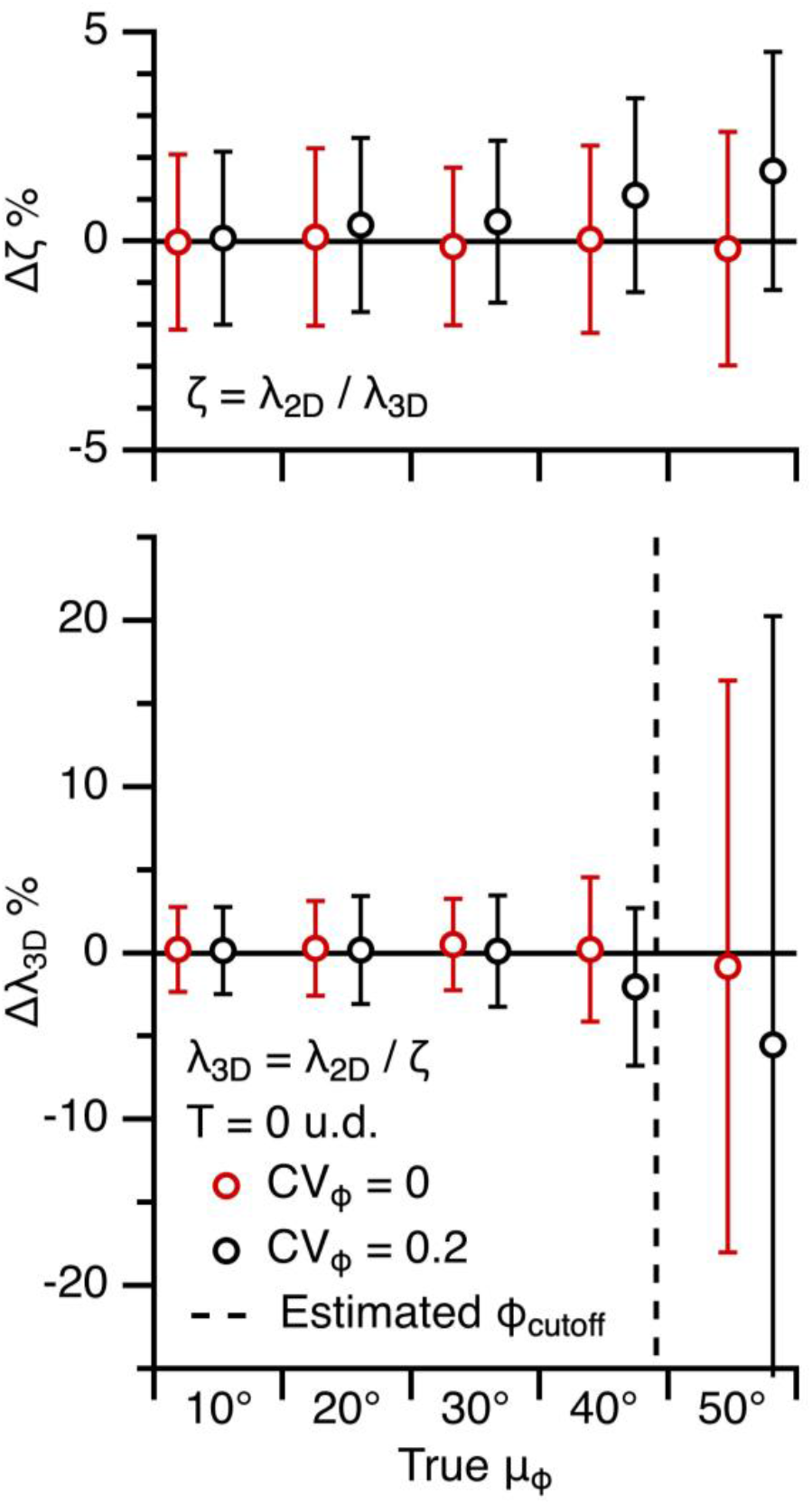
The Keiding-model fixed-ϕ assumption introduces only small errors when estimating λ_3D_ from λ_2D_ (Part II. 2D projection simulations). Average Δζ (top) and Δλ_3D_ (bottom), as described in Figures 10 and S18, for the fix-ϕ and Gaussian-ϕ simulations in Figure S11 (CV_ϕ_ = 0 or 0.2; red vs. black) and T = 0. The small positive biases in Δζ for the Gaussian-ϕ simulations indicate estimated ζ is marginally too large (i.e. Equation 2 is only approximately correct for Gaussian-ϕ conditions); however, these biases create negligible biases in λ_3D_ for μ_ϕ_ < 50°. Estimated ϕ_cutoff_ ≈ 43° (Equation 9). Fixed-ϕ data is from Figure 10.

